# A pair of congenic mice for imaging of transplants by positron emission tomography using anti-transferrin receptor nanobodies

**DOI:** 10.1101/2024.11.20.624506

**Authors:** Thomas Balligand, Claire Carpenet, Sergi Olive-Palau, Tom Jaspers, Pavana Suresh, Xin Liu, Himadri Medhi, Yoon Ho Lee, Mohammad Rashidian, Bart De Strooper, Hidde L. Ploegh, Maarten Dewilde

## Abstract

Two anti-transferrin receptor (TfR) nanobodies, V_H_H123 specific for mouse TfR and V_H_H188 specific for human TfR (huTfR) were used to track transplants non-invasively by PET/CT in mouse models, without the need for genetic modification of the transferred cells. We provide a comparison of the specificity and kinetics of the PET signals acquired when using nanobodies radiolabeled with ^89^Zr, ^64^Cu and ^18^F, and find that the chelation of the ^89^Zr and ^64^Cu radioisotopes to anti-TfR nanobodies results in radioisotope release upon endocytosis of the radiolabeled nanobodies. We used a knock-in mouse that expresses a TfR with a human ectodomain (huTfR ^+/+^) as a source of bone marrow for transplants into C57BL/6 recipients and show that V_H_H188 detects such transplants by PET/CT. Conversely, C57BL/6 bone marrow and B16.F10 melanoma cell-line transplanted into huTfR ^+/+^ recipients can be imaged with V_H_H123. In C57BL/6 mice impregnated by huTfR^+/+^ males we saw an intense V_H_H188 signal in the placenta, showing that TfR-specific V_H_Hs accumulate at the placental barrier but do not enter the fetal tissue. We were unable to observe accumulation of the anti-TfR radiotracers in the central nervous system (CNS) by PET/CT but show evidence of CNS accumulation by radiospectrometry. The model presented here can be used to track many transplanted cell types by PET/CT, provided cells express TfR, as is typically the case for proliferating cells such as tumor lines.

## Introduction

Non-invasive tracking of specific cell types *in vivo* is a desirable goal in many fields of biomedical research, particularly as it pertains to immunology and cancer. Knowledge of the biodistribution of immune cells and/or tumor cells in a disease model is essential to monitor therapeutic interventions. Real-time non-invasive *in vivo* imaging may be achieved by several methods, each with their strengths and limitations. Fluorescence and luminescence-based methods suffer from absorbance and dispersal of emitted light ^1^, which limits the depth at which images of acceptable resolution can be obtained. Fluorescence-based methods in conjunction with multi-photon microscopy provide cellular resolution, but usually require invasive surgery to gain access to the cells of interest ^2^. Methods that rely on the use of radio-isotopes, such as single photon emission computed tomography (SPECT) and positron emission tomography (PET) do not suffer those drawbacks. While SPECT and PET lack the resolution of optical microscopy, they have the advantage of being non-invasive, quantitative and enabling whole-body imaging in small animals at a resolution of ∼1 mm, or a 1 microliter volume, for PET ^3^ and even lower than 1 mm for SPECT ^4^. These methods are finding increasing use in the field of tumor immunology, as they can provide a whole-body image, unlike other approaches.

Current PET/CT cell tracing methods require the use of a cell-specific tracer. This typically relies on the detection of an endogenous surface marker that is specific for the cells of interest ^5^, or by genetically engineering cells of interest to enable their visualization, for example through expression of a viral kinase of unique specificity ^6^, or by introduction of a surface marker that can be selectively visualized ^7^.

V_H_Hs, also termed nanobodies, are enjoying increasing use for the generation of immuno-PET tracers that yield images of a quality superior to what is achieved using regular, intact immunoglobulins ^8^. Nanobodies are the recombinantly expressed variable fragments (V_H_) of heavy chain-only immunoglobulins produced by camelids ^9, 10^. Their small size (∼15 kDa), superior tissue penetration, specificity, affinity and much shorter circulatory half-life compared to intact immunoglobulins make nanobodies excellent tracers for *in vivo* imaging. We here apply nanobodies specific for the transferrin receptor to track the fate of transplanted cells non-invasively.

The transferrin receptor (TfR; CD71; encoded by *Tfrc*) is a homodimeric type-II transmembrane protein that is near-ubiquitously expressed, in particular on proliferating cells ^11^. This includes many tumor cells with some variability depending on the tumor type and differentiation status ^12–20^. The TfR binds to and endocytoses iron (Fe^+++^)-loaded transferrin (Tf). Tf remains bound to the TfR in endosomal compartments, where the resident low pH releases the Fe^+++^ cargo and thus converts Tf into apoTf, which remains TfR-bound at endosomal pH. From there, TfR-bound apoTf returns to the cell surface, where the TfR releases apoTf at neutral pH. TfR is also expressed by endothelial cells that line the blood-brain barrier (BBB) for delivery of Fe^+++^-loaded transferrin to the central nervous system by transcytosis. The nanobodies used in this study bind TfR and can traverse the BBB ^21, 22^. Mouse embryos critically depend on an iron supply in the form of Tf captured from the maternal circulation, delivered to the embryo via the TfR expressed on the syncytiotrophoblast-I ^23^. Whether the small size of nanobodies allows them -by analogy with the BBB- to traverse the placenta and reach the embryo has not been explored.

We use two nanobodies: V_H_H123 (also termed Nb62 ^21^) and V_H_H188, which recognize the murine TfR and the human TfR (huTfR), respectively. We use these V_H_Hs together with a knock-in mouse model that expresses a TfR with a human TfR ectodomain (huTfR ^+/+^) ^22^. The specificity of each nanobody for the respective TfR and the availability of huTfR^+/+^ mice as a source of primary cells allows us to track different cell types in a transplant setting. V_H_H123 allows the detection of cells of mouse origin, such as bone marrow progenitors or tumor cells, transplanted into huTfR^+/+^ mice. Conversely, we use V_H_H188 to track primary cells from huTfR^+/+^ mice after transfer into wild-type mice. Pregnancy represents a unique model akin to a transplant. We report the first immuno-PET study on localization of the TfR in mouse embryos in live mice.

We find that ^89^Zr and ^64^Cu, ligated to these TfR-specific V_H_Hs by non-covalent chelation, are released from the imaging agent upon binding the TfR, due to internalization and exposure to low endosomal pH. The release of free ^89^Zr and ^64^Cu from imaging agents are factors to consider when using these over extended periods of time. Covalent modification of V_H_Hs with ^18^F avoids the release of free radioisotope, but the short half-life of ^18^F limits the observation window to <12 hours. We compare the use of three commonly used positron-emitting isotopes (^89^Zr, ^64^Cu and ^18^F) and offer a suite of tools to track many cell types in mice, without the requirement for specific cell markers or genome-editing. At most, crossing of a mouse model of interest with the huTfr^+/+^ mouse line would be required (deposited at Jackson laboratories, strain 038212).

## Results

### V_H_H123 binds mouse TfR and V_H_H188 binds human(ized) TfR *in vitro* and *in vivo*

We characterized the specificity of V_H_H123 (anti-mouse TfR) and V_H_H188 (anti-human TfR) by biochemical methods and by PET/CT. We confirmed the specificity of each anti-TfR V_H_H for its target by immunoprecipitation from lysates of HEK293 (human) or B16.F10 (mouse) cell lines. We show that both V_H_Hs bind only to the appropriate TfR, with no obvious cross-reactivity to other surface-expressed proteins by immunoblot, LC/MSMS analysis of immunoprecipitates, SDS-PAGE of ^35^S-labelled proteins and flow cytometry (**Fig 1** ;**Table 1; Supplemental Fig 15**). Virtually all contaminants that were co-immunoprecipitated were of cytoplasmic and nuclear localization (**Table 1; supplementary file**). This is of minor concern, as these proteins are not accessible to our anti-TfR nanobodies in an *in vivo* setting.

**Figure 1.**
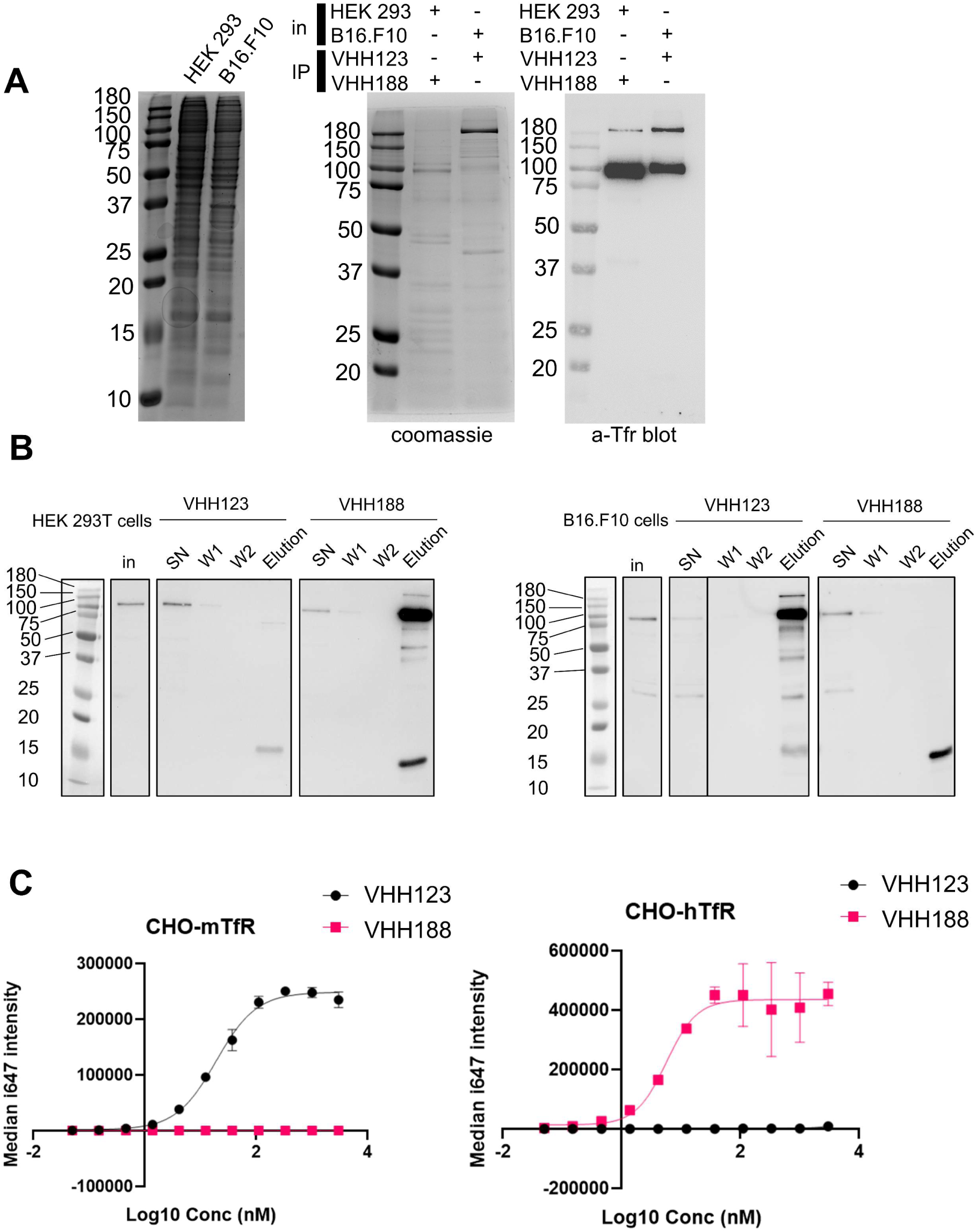
**A.** Left-most panel: Coomassie staining of an SDS-PAGE gel showing the input cell lysates used in immunoprecipitation experiments depicted in the next panels. Middle panel: lysates as shown on left-side panel were incubated with nanobody coated paramagnetic beads overnight. Beads were then washed 5 times then boiled in SDS sample buffer before loading on SDS-PAGE. This panel shows Coomassie staining of SDS-PAGE of the bead eluates. In: input lysates. IP: Nanobody used for immunoprecipitation. Right panel: the same eluates as in middle panel were run on SDS-PAGE gel then transferred to a PVDF membrane. Membrane was blocked, stained using a mouse monoclonal anti-TfR (cross-reactive for human and mouse), washed then stained with an anti-mouse-HRP secondary monoclonal. See Methods section for more details. **B.** Cell-lines were incubated with ^35^S-labelled Met before performing the same immunoprecipitation procedure as described in A, with the exception that the beads were washed only twice before elution in Laemli buffer. SDS-PAGE was run with input cell lysate (in), unbound fraction (SN), washes (W1 and W2) and eluates for each condition before transfer to a PVDF membrane. Membrane was blocked, stained using a mouse monoclonal anti-TfR (cross-reactive for human and mouse), washed then stained with an anti-mouse-HRP secondary monoclonal. **C.** Flowcytometry characterization of the specificity of V_H_H123 and V_H_H188. CHO cells overexpressing either the mouse isoform of TfR (mTfr, left panel) or human isoform (hTfr, right panel) were labelled with serial dilutions of either Flag-tagged V_H_H123 or V_H_H188. I647-fluorescently labeled anti-FLAG IgG was used to detect the presence of either VHH at the cell-surface by flow cytometry.

**Table 1.**
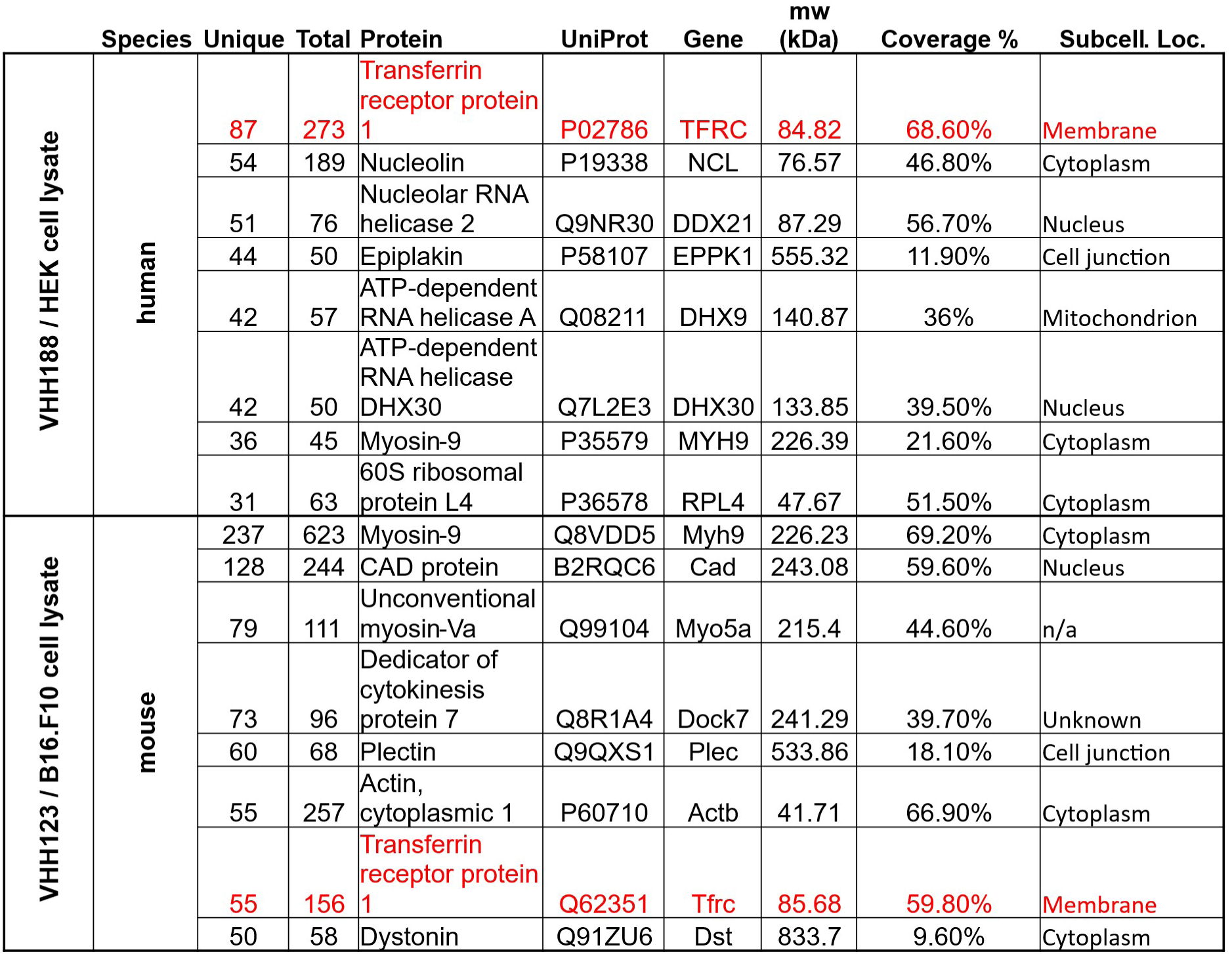
Summary of peptides identified from LC/MS/MS analysis of the gel sections from figure 1A, middle panel. Transferrin receptor 1 is highlighted in red. See supplementary Table 1 for full dataset.

For PET experiments we attached a deferoxamine (DFO)-azide moiety to each V_H_H via a sortase A-catalyzed transpeptidation reaction ^24^. The V_H_H-DFO-azide adducts were then modified with dibenzocyclooctyne-polyethyleneglycol_20kDa_ (DBCO-PEG_20kDa_) using click chemistry (**Fig 2, Supplemental Fig 1, 9, 10**). The PEG_20kDa_ moiety, hereafter referred to as ‘PEG’, serves to extend the half-life of the V_H_H in the circulation and decreases non-specific kidney uptake ^24^. The PEG-DFO-modified V_H_Hs were labeled with ^89^Zr through chelation by DFO (see methods) to generate the V_H_H-PEG-DFO-^89^Zr radiotracers. We injected the V_H_H-PEG-DFO-^89^Zr conjugates into C57BL/6 mice and huTfR^+/+^ mice and collected PET/CT images at various times after retro-orbital injection. Injection of the V_H_H123-based conjugate into huTfR^+/+^ mice yielded a signal exclusively in the kidneys, which we consider a non-specific signal. ^89^Zr radiolabeled V_H_Hs typically accumulate in the kidneys in the absence of a specific target. Similarly, injection of the V_H_H188-based conjugate into C57BL/6 mice likewise yielded a strong PET signal only in the kidneys (**Fig 3B-C**), again considered non-specific accumulation of the tracer.

**Figure 2.**
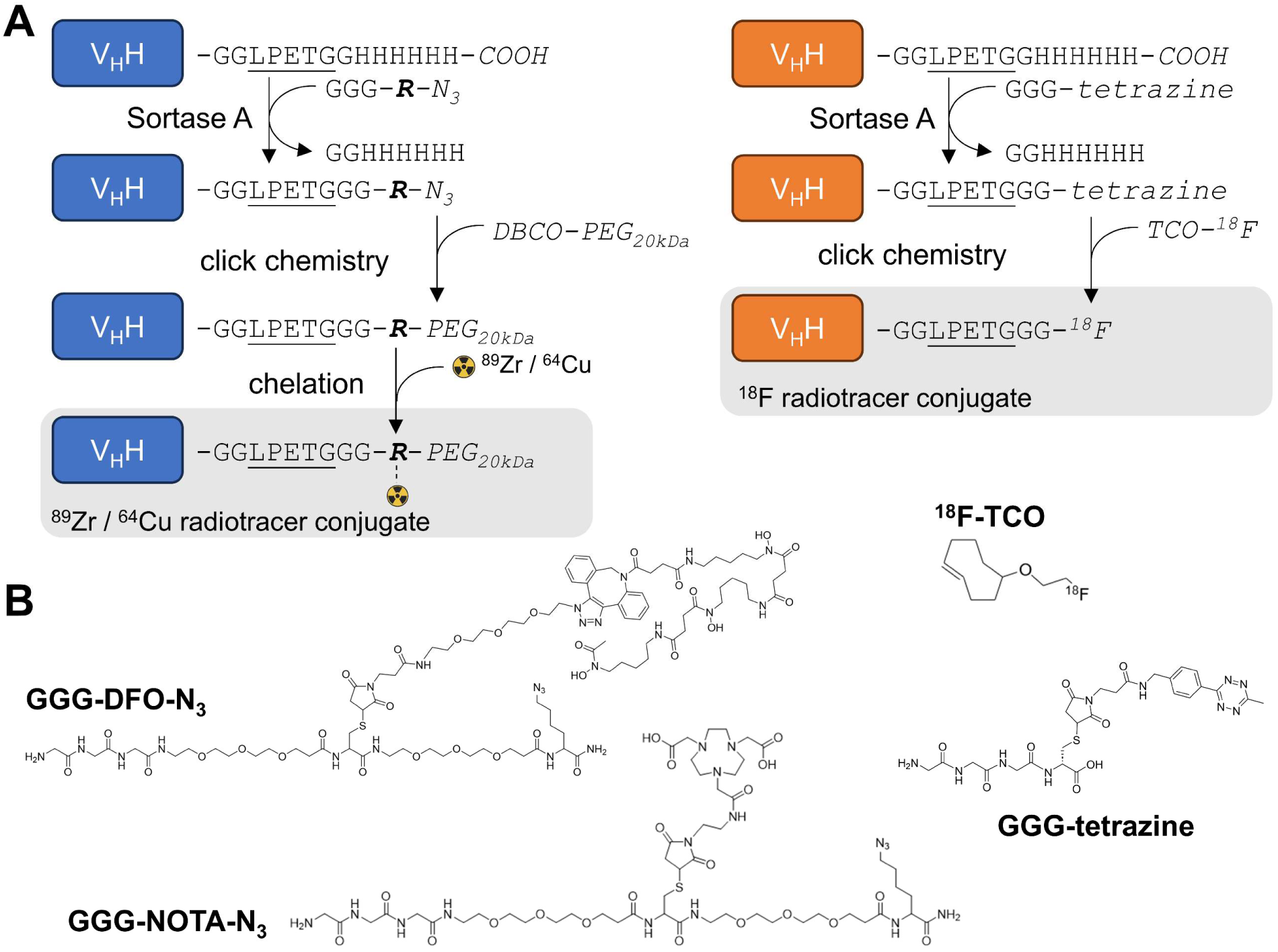
**A.** Schematic of sortase A and click-chemistry steps to generate each radio-conjugate used in this study. Capital letters denote amino-acids, unless if written in *italic* where they denote chemical groups or elements. Bold italic ‘R’ represents either DFO (deferoxamine) or NOTA (2,2′,2”-(1,4,7-triazacyclononane-1,4,7-triyl)triacetic acid). DBCO: dibenzocyclooctyne, PEG_20kDa_: poly-ethylene-glycol (20kDa mw), TCO: trans-cyclooctene. Underlined is the LPETG Sortase A cleavage site consensus motif. See suppl. Fig 1 for detailed methods. **B.** Structures of GGG-nucleophiles used in sortase A mediated conjugations (GGG-DFO-N_3_, GGG-NOTA-N_3_ and GGG-tetrazine). The ^18^F-TCO click-chemistry partner of GGG-tetrazine is also depicted. These structures were synthesized as described in supplementary methods and figures 12 and 13.

**Figure 3.**
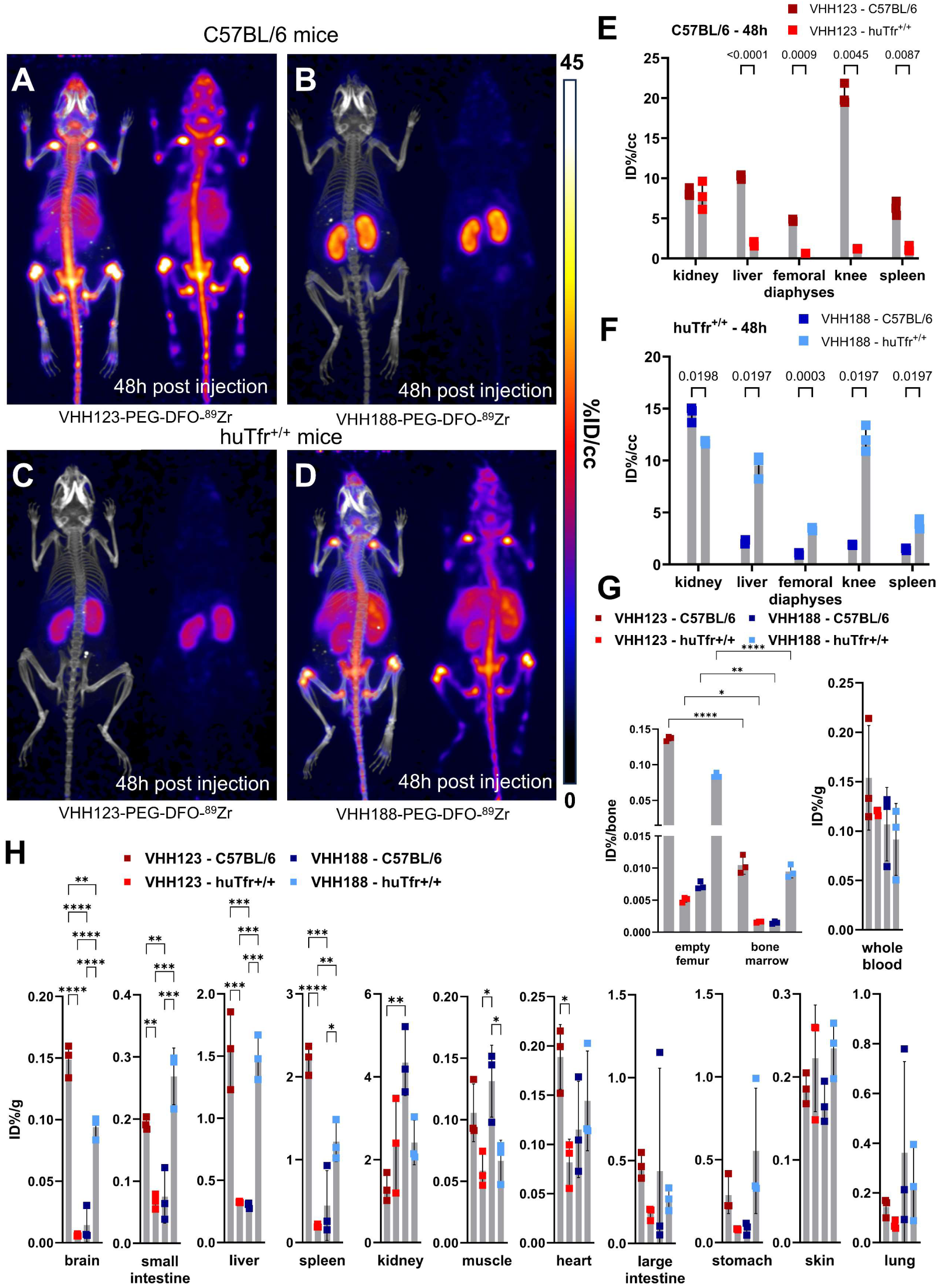
**A-D:** C57BL/6 and huTfR^+/+^ mice were injected with 3.7 MBq (100 µCi) of either V_H_H123-PEG(20kDa)-DFO-^89^Zr or V_H_H188-PEG(20kDa)-DFO-^89^Zr by retro-orbital injection. The mice were imaged by PET/CT at several timepoints post-injection. Shown here at the maximum intensity projection images acquired at 48 hours post injection of the conjugate. Each panel comprises maximum intensity projection (MIP) overlayed with CT signal on the left, and PET MIP alone on the right. PET intensity scale is displayed on the right (%ID/cc). **A.** C57BL/6 mouse injected with the V_H_H123 based conjugate. **B.** C57BL/6 mouse injected with the V_H_H188-based conjugate. **C.** huTfR^+/+^ mouse injected with the V_H_H123-based conjugate. **D.** huTfR^+/+^ mouse injected with the V_H_H188-based conjugate. Experiment performed with 3 mice per condition, with one mouse shown as representative of each condition. **E.** Region Of Interest (ROI) analysis of images acquired from mice as shown in A and C and all repeats thereof. The mean ID%/cc is plotted for each ROI and mouse repeat. **F.** Same as E, but for images acquired from mice as shown in B and D and all repeats thereof. **G**. Left graph: *Ex vivo* activity measurement of flushed femurs (thus mineral bone) and the bone marrow they contained, 72 hours post radiotracer injection as in A-D. Each dot represents measurement of one mouse on a scale of injected dose percentage per bone (ID%/bone). Right graph: ex vivo activity measurement of 20 µL of whole blood Each dot represents the activity from one mouse, on a scale of injected dose percentage per gram of tissue (ID%/g). Bars show SD. **H.** *Ex vivo* activity measurements from different tissues, performed at 72 hours post radiotracer injection. No capillary depletion was performed. Each dot represents one measurement from one mouse on a scale of ID%/g. Bars show SD.

For the properly matched V_H_H-mouse combinations (^89^Zr radiolabeled V_H_H123 conjugate injected into C57BL/6 mice; ^89^Zr radiolabeled V_H_H188-conjugate injected into huTfR^+/+^ mice), each produced an intense PET signal that localizes to bones, mainly the vertebrae, sacrum, coxal bone as well as both epiphyses of the femur and the proximal epiphysis of the humerus at 48 hours post-radiotracer injection (**Fig3A and D**). The mean activity measured by PET was significantly different between matched and unmatched V_H_H/mouse pairs in the liver, spleen, femoro-tibial articulation (knee) and spleen, by region of interest (ROI) analysis (**Fig 3E-F**). While the signals from long bone extremities and vertebrae suggest uptake of the anti-TfR radioconjugate by bone marrow ^25^, presumably due to the high demand for iron required for erythropoiesis ^26^, this pattern of label distribution is better explained by sequestration of free ^89^Zr in mineralized bone and cartilage ^27, 28^ (see also supplemental Fig. 8). The weak signal from the diaphyses of the femora relative to the intense signal in the vertebral column and bone extremities suggests that accumulation of free ^89^Zr in mineralized bone predominates under these conditions. To assess this, we measured the activity of specific tissues *ex vivo* 72 hours post radiotracer injection (**Fig. 3G-H**). Several tissues showed significant differences of activity between the matched and unmatched mouse/V_H_H pairs: particularly in the brain, spleen, small intestine and liver. Femur bones flushed of all bone marrow retained a very strong activity in the matched conditions – showing that the mineral bone retains a high activity when depleted of bone marrow (> 200,000 CPM/bone) – confirming the deposit of free ^89^Zr in the bone matrix. Nevertheless, flushed bone marrow showed activity (20000 – 30000 CPM/flushed bone) that was significantly higher in the matched vs. unmatched conditions (**Fig 3G**). We conclude that at later timepoints, the strong PET signals in the long bone extremities (femur, tibia and humerus) are due in part to bone marrow accumulation of anti-TfR radiolabeled V_H_Hs, but predominantly so to accumulation of free ^89^Zr in the mineralized bone matrix. No obvious accumulation of tracer was seen in brain parenchyma by PET, but significant differences of activity were found in the *ex vivo* analysis (**Fig 3H**), in line with the reported ability of V_H_Hs that recognize the TfR to deliver bound materials across the blood brain barrier ^21^. The failure to detect a PET signal of adequate strength in the brain are due to the comparatively small amounts of anti-TfR V_H_H that cross the BBB. When analyzing PET images obtained with V_H_Hs that recognize targets other than the TfR, which includes V_H_Hs that recognize CD8 ^24^, Ly6C/G ^29^ or CD11b ^30^, as examined previously, no accumulation of ^89^Zr in skeletal elements was seen. These previous observations, combined with the lack of any bone signal in the mismatched anti-TfR V_H_H/mouse pairs, lead us to conclude that free ^89^Zr is released from the V_H_H-conjugates *in vivo,* but only when the V_H_H binds to its respective TfR.

Because chelation of ^89^Zr to DFO depends on its charge ^31^, we hypothesized that binding of V_H_H-^89^Zr conjugates to the appropriate TfR delivers them to endosomal compartments of low pH, where ^89^Zr is then released from the DFO moiety. We examined release of ^89^Zr from the V_H_H123-PEG-DFO construct *in vitro* by exposure to low pH (**Suppl Fig 2**). ^89^Zr is freed from the adduct at pH <5.6, which is well within the pH range of late endosomal compartments ^32^. Whether the mildly acidic pH of early or recycling endosomal compartments is sufficient for isotope release *in vivo* remains to be determined.

### Comparison of ^89^Zr, ^64^Cu and ^18^F isotopes conjugated to the anti-TfR V_H_Hs

Having observed the release of ^89^Zr from the V_H_H123-PEG-DFO imaging agent, we considered the use of ^64^Cu as alternative radio-isotope. We also synthesized an adduct covalently labeled with ^18^F to circumvent any possible release of free radioisotope. The rather different half-lives of ^89^Zr (∼3.3 days), ^64^Cu (∼12 hours) and ^18^F (∼110 minutes) dictate the observation windows allowed by their use, which we arbitrarily set at 3 to 5 half-lives, thus ranging from 10 hours (^18^F) to ∼2 weeks (^89^Zr), with ∼3% of the injected dose remaining after 5 half-lives due to isotope decay.

We generated a V_H_H123-PEG-NOTA-^64^Cu version, where ^64^Cu is chelated by NOTA and would not accumulate in mineralized bone if released. The use of a ^64^Cu-labeled tracer should allow an observation window of 3-5 half-lives, i.e. ∼36-60 hours. For comparison we produced a V_H_H123-tetrazine-TCO-^18^F construct, where ^18^F is incorporated covalently through tetrazine-TCO click-chemistry, by an inverse electron-demand Diels–Alder (IEDDA) reaction between V_H_H123-tetrazine and ^18^F-TCO ^33^. This precludes isotope release other than by proteolysis. We injected these imaging agents for a side-by-side comparison with V_H_H123-PEG-DFO-^89^Zr in C57BL/6 mice (**Fig 4A**). At 1 hour post injection of the ^89^Zr, ^64^Cu or ^18^F-labeled radiotracer, we observe a strong PET signal in the diaphyses and both epiphyses of the femora and in the coxal bones and sacrum (bone marrow), thus showing that V_H_H123 accumulates specifically at those locations, and that the signal is not simply due to free radio-isotope only. ^89^Zr and ^64^Cu radiolabeled V_H_H123also accumulated in the spleen. The ^18^F based imaging agent provided a sharper picture, with some signal originating from the gut and gall bladder, typically seen when using ^18^F-based radiotracers comprising bulky hydrophobic moieties, in this case a TCO-Tetrazine clicked product ^34^. At the 12h timepoint the ^18^F signal had fully decayed due to the short half-life of ^18^F. Improved signal quality is thus offset by the shorter half-life of ^18^F. For the ^64^Cu-based agent, the bone marrow signal remains weakly visible at 24 hours post injection. The liver and gut signals increase at later timepoints for the ^64^Cu-labeled tracer, a phenomenon attributable to a slow release of ^64^Cu from NOTA to plasma proteins such as albumin, which delivers copper to the liver where it can then be conjugated to ceruloplasmin ^35, 36^. In mice that received the ^89^Zr-labeled agent we mostly observe a signal from the bone extremities and vertebrae at 24 hours post injection, which we attribute to labelling of mineralized bone with free ^89^Zr with a minor contribution of a TfR-specific bone marrow signal. For all imaging agents, we observe a pair of punctiform signals in the anterior region of the cranium. These appear very prominently with the ^18^F tracer, and at later timepoints -but less clearly-with the ^89^Zr and ^64^Cu imaging agents, potentially explained by the lower positron emission range of ^18^F. Limited by the resolution of the CT images, we tentatively attribute these signals to the accumulation of the radiotracer in the roots of the incisors, where iron uptake is required for proper amelogenesis ^37^

**Figure 4.**
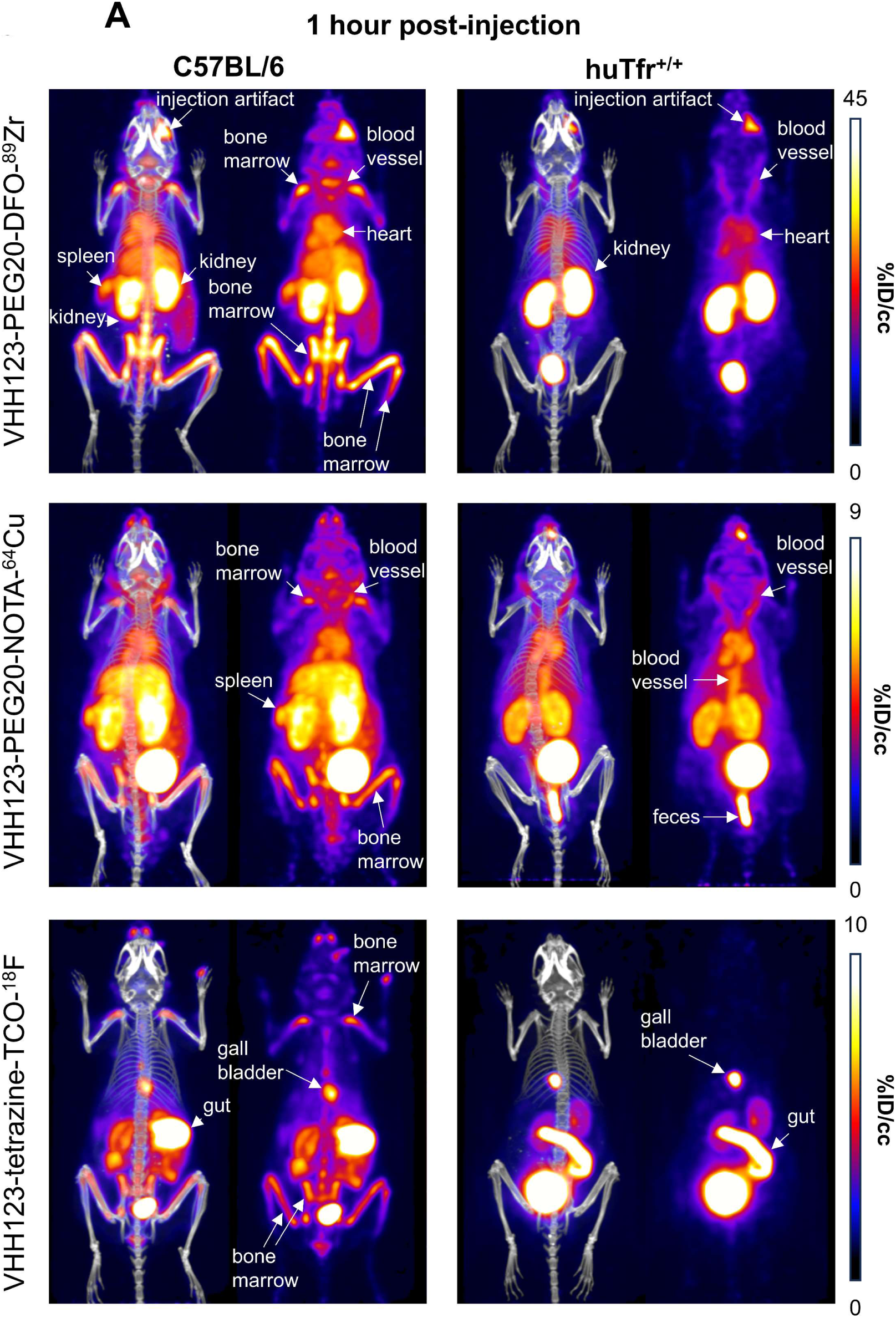
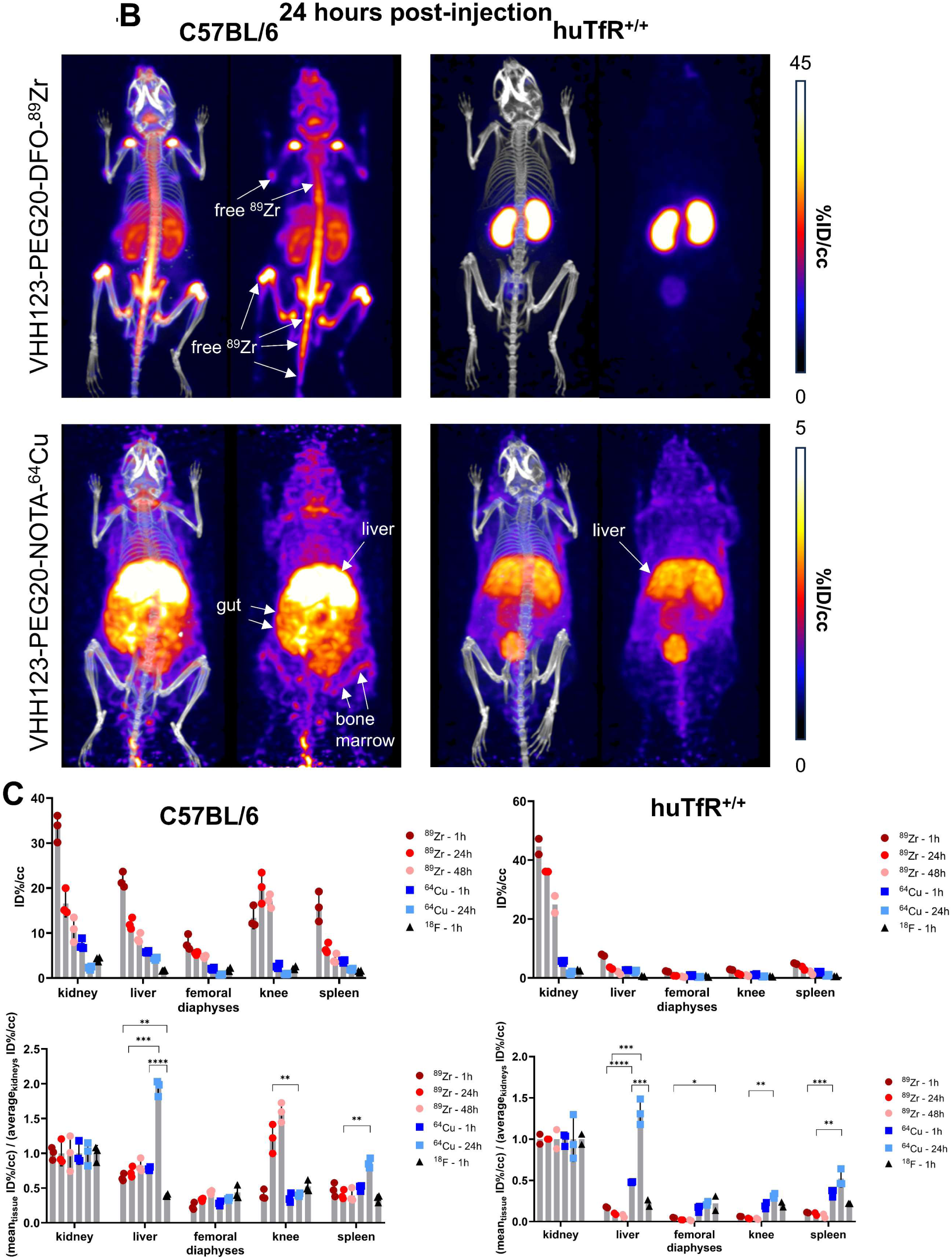
**A-B.** C57BL/6 mice (left column) and huTfR^+/+^ mice (right column) were injected with 3.7 MBq (100µCi) of different V_H_H-123-PEG(20kDa) based conjugates as indicated on the left side of the panels. PET/CT images were acquired for each condition at 1 hour post injection (A) and at 24 hours post injection (B) (barring ^18^F which was not imaged at 24 hours). Each panel comprises maximum intensity projection (MIP) overlayed with CT signal on the left, and PET MIP alone on the right. PET intensity scales are displayed on the right of each row (%ID/cc). This figure pools the representative pictures obtained from 3 independently performed sets of experiments where one specific V_H_H123-radiolabelled conjugate was tested per experiment. 3 C57BL/6 mice were imaged in each experiment for each condition, and 2 huTfR^+/+^ mice were imaged in each experiment for each condition, save for the ^64^Cu condition where 3 huTfR^+/+^ mice were imaged. **C.** ROI analysis of images acquired from mice as shown in all panels of A (left column – C57BL/6) and B (right column – huTfR^+/+^), and all repeats thereof. Top row: each point represents the mean ID%/cc for one mouse. Bottom row: same data as in the graph above, but each point represents the mean ID%/cc of a specific tissue normalized to the average ID%/cc values found in the kidneys of the same group.

We conclude that internalization of ^89^Zr and ^64^Cu V_H_H123-based imaging agents leads to release of free radioisotope (^89^Zr and ^64^Cu), which is responsible for most of the signal detected at later (24h or later) timepoints and thus does not truly reflect distribution of the TfR itself at those timepoints. This is particularly clear from a ROI analysis of the liver and knee PET signals, when normalized to the average kidney signal (**Fig 4B**), that shows a shift of distribution of PET signal towards the liver with ^64^Cu-labelled V_H_H123 and towards the knee for ^89^Zr-labelled VHHs at later timepoints post injection. Combined, these results establish specificity of recognition of the respective V_H_Hs for their intended targets. Moreover, the results show that release of ^89^Zr or ^64^Cu from the imaging agents is the unavoidable consequence of their binding to the proper targets. The choice of isotopes for conjugation to anti-TfR nanobodies must therefore be made with respect to the tissue(s) that are to be visualized, as well as the time scale of the experiment (∼3 to 5 isotope half-lives).

### ^89^Zr-conjugated V_H_H123 tracks transplanted bone marrow cells *in vivo*

Having established that V_H_H123 accumulates in the bone marrow *in vivo,* we tested whether V_H_H123 could detect bone marrow cells transplanted into lethally irradiated recipient mice. To enable the specific detection of the donor cells, we isolated bone marrow cells from C57BL/6 mice and transplanted 1.5×10^6^ cells into lethally irradiated (10 Gy) huTfR^+/+^ recipient mice by retro-orbital injection. We used V_H_H123-PEG-DFO-^89^Zr to determine whether bone marrow engraftment would suffice to reproduce the signal pattern observed in C57BL/6 wild-type mice. At 2 weeks post bone marrow transplantation, 24 hours after injection of V_H_H123-PEG-DFO-^89^Zr, the presence of the transferred cells marrow is observed in the spleen and femoral bone, together with a strong signal from free ^89^Zr accumulation in the bone matrix (**Fig 5A**). The signal in the spleen appeared more prominent than that observed when imaging wild-type C57BL/6 mice at 24 hours post injection of the same VHH, as pointed out through ROI analysis (**Fig 5F** vs. **4B**), which could reveal the presence of hematopoietic bone marrow in the spleen. The same observations were made in the reverse experiment where transplantation of huTfR^+/+^ bone marrow into C57BL/6 recipients was performed before imaging the recipients 15 days after transplantation using V_H_H188-PEG-DFO-^89^Zr (**Fig 5B and 5G**). Strikingly, the images acquired at 1 hour post V_H_H188-PEG-DFO-^89^Zr injection show a powerful signal in the spleen, something not observed when imaging huTfR^+/+^ mice with the same radiotracer, which can be easily interpreted as the presence of bone marrow engraftment in the spleen (**Fig 5C and 5G**). As a control, V_H_H188-PEG-DFO-^89^Zr was unable to show engraftment of C57BL/6 bone marrow cells in an isogenic recipient (**Fig 5D and 5G**). Because the transferred bone marrow cells proliferated in the 14 days prior to imaging, they reached numbers adequate for the release of ^89^Zr that would then accumulate in the bone matrix. Indeed, imaging of huTfR^+/+^ recipient mice immediately after C57BL/6 bone marrow transplantation using V_H_H188-PEG-DFO-^89^Zr reveals only a faint signal in the knee and liver (**Fig 5E**). Specific tracing of TfR-positive cells of mouse or human origin is thus possible when using V_H_H123 or V_H_H188 as the tracer in a huTfR^+/+^ or wild-type recipient, respectively. Depending on the mass of TfR^+^ cells present the recipients, timing of tracer injection and the PET imaging session is critical to avoid the confounding effect of the free ^89^Zr bone matrix signal. These data establish the feasibility of detecting transplanted bone marrow cells and their hematopoietic descendants amidst populations or recipient cells that remain invisible to the imaging agent used.

**Figure 5.**
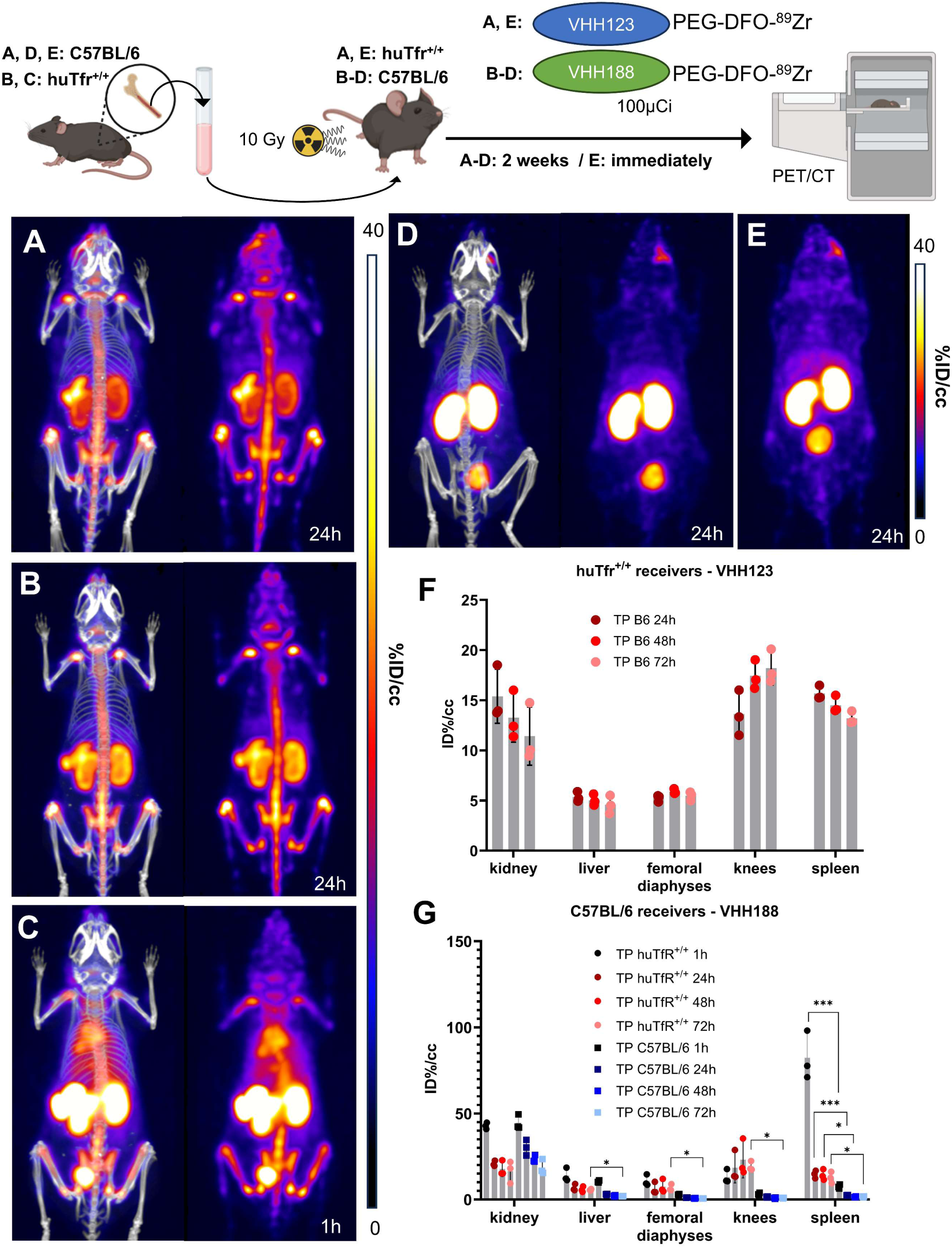
Top: cartoon depicting the experimental procedure: bone marrow was harvested from the femur of C57BL/6 (A, D and E) or huTfR^+/+^ (B and C) mice before transplantation into 3 lethally irradiated (10 Gy) huTfR^+/+^ (A and E) or C57BL/6 (B, C and D) mice. These mice were then injected with 3.7 MBq (100µCi) of V_H_H123-PEG(20kDa)-DFO-^89^Zr (A and E) or V_H_H188-PEG(20kDa)-DFO-^89^Zr (B, C and D) immediately (E) or 2 weeks post transplantation (A, B, C and D) and PET/CT images were acquired at several timepoints thereafter. Bottom: PET/CT maximum intensity projections (MIP) that were deemed the most representative of each condition are shown. Each panel comprises maximum intensity projection (MIP) overlayed with CT signal on the left, and PET MIP alone on the right**. A**: MIP of one huTfR^+/+^ recipient mouse acquired 2 weeks after C57BL/6 bone marrow transplantation and 24 hours after radiotracer injection. **B**: MIP of one C57BL/6 mouse acquired 2 weeks after huTfR^+/+^ bone marrow transplantation and 24 hours after radiotracer injection. **C.** Same as B, but imaged 1 hour after radiotracer injection. **D.** MIP of one C57BL/6 mouse acquired 2 weeks after C57BL/6 (isogenic) bone marrow transplantation and 24 hours after radiotracer injection. **E.** MIP of one huTfR^+/+^ recipient mouse injected with radiotracer immediately after C57BL/6 bone marrow and imaged 24 hours thereafter. Two cohorts were set up separately to perform the PET/CT imaging immediately after bone marrow transplantation or two weeks thereafter. PET intensity scales are displayed on the right of each panel (%ID/cc). **F.** ROI analysis of images acquired from mice as shown in panel A, and all repeats and imaging timepoints thereof. Error bars show SD. **G.** ROI analysis of images acquired from mice as shown in panels B, C and D and all repeats and imaging timepoints thereof. Error bars show SD.

### TfR-positive B16.F10 melanoma cells are detected by ^64^Cu-conjugated V_H_H123

Next, we asked whether we could trace tumor cells that express mouse TfR, but that are not derived from bone marrow hematopoietic cells. 5×10^4^ B16.F10 mouse melanoma cells were injected *i.v.* by tail vein injection to generate metastatic lung tumors in huTfR^+/+^ mice. We allowed the B16.F10 cells to engraft and establish metastases for up to 4 weeks post-transplantation. We injected the recipients with V_H_H123-PEG-NOTA-^64^Cu at weeks 2 and 4. Imaging done at 2 weeks post-inoculation of the B16F10 tumor did not yield a clear signal in the lungs (**Fig 6A**), as metastases typically arise some 3 weeks after injection for this number of cells. At 4 weeks post transplantation, we observed a clear signal originating from the lungs of the same mice (**Fig 6C**). No signal was detected in the no tumor control group (**Fig 6E**). To exclude the possibility of non-TfR specific accumulation of radiotracer in necrotic tumor tissue, we injected tumor bearing mice with a non-specific anti-GFP V_H_HEnh-PEG-NOTA-^64^Cu conjugate (**Fig 6B,D**), which showed only weak passive accumulation in the lung metastases at 4 weeks post tumor cell infusion. At necropsy we confirmed the presence of melanotic tumors in the lungs (**Fig 6F**), which were also visible on the lung CT. One mouse also had a small tumor in the liver, and another in the skin of the left flank. No mice had tumors in the heart or kidneys. By ROI analysis of the PET signals, V_H_H123 gave a significantly higher signal in the whole lungs (**Fig 6G**). The PET signal from B16.F10 metastases in the lung correlates well with what appears to be dense tumor tissue on the corresponding CT image. When performing ROI analysis restricted to the PET signal in hyperdense lung tumor tissue, V_H_H123-PEG-NOTA-^64^Cu gave an overall stronger signal when compared to V_H_HEnh-PEG-NOTA-^64^Cu, although not by a significant margin (**Fig 6H**). It is important to emphasize that ROI analysis on lung and heart is less precise, due to breathing movements and beating heart during acquisition of the PET signal. Kidney signals were significantly different between tumor-free mice that received V_H_H123-PEG-NOTA-^64^Cu and the other groups, which we attribute to a difference in clearance rate of the radioconjugates. The anti-mouse TfR nanobody V_H_H123 is thus well-suited to track TfR-expressing cells of mouse origin in huTfR^+/+^ recipient mice. Since all proliferating cells express TfR, in principle any tumor or proliferating cell of mouse origin can be detected non-invasively without the need for genetic modification of the transplanted cells. This congenic pair of mice, in combination with the species specificity of the anti-TfR V_H_Hs, is thus a unique tool for studies of this type.

**Figure 6.**
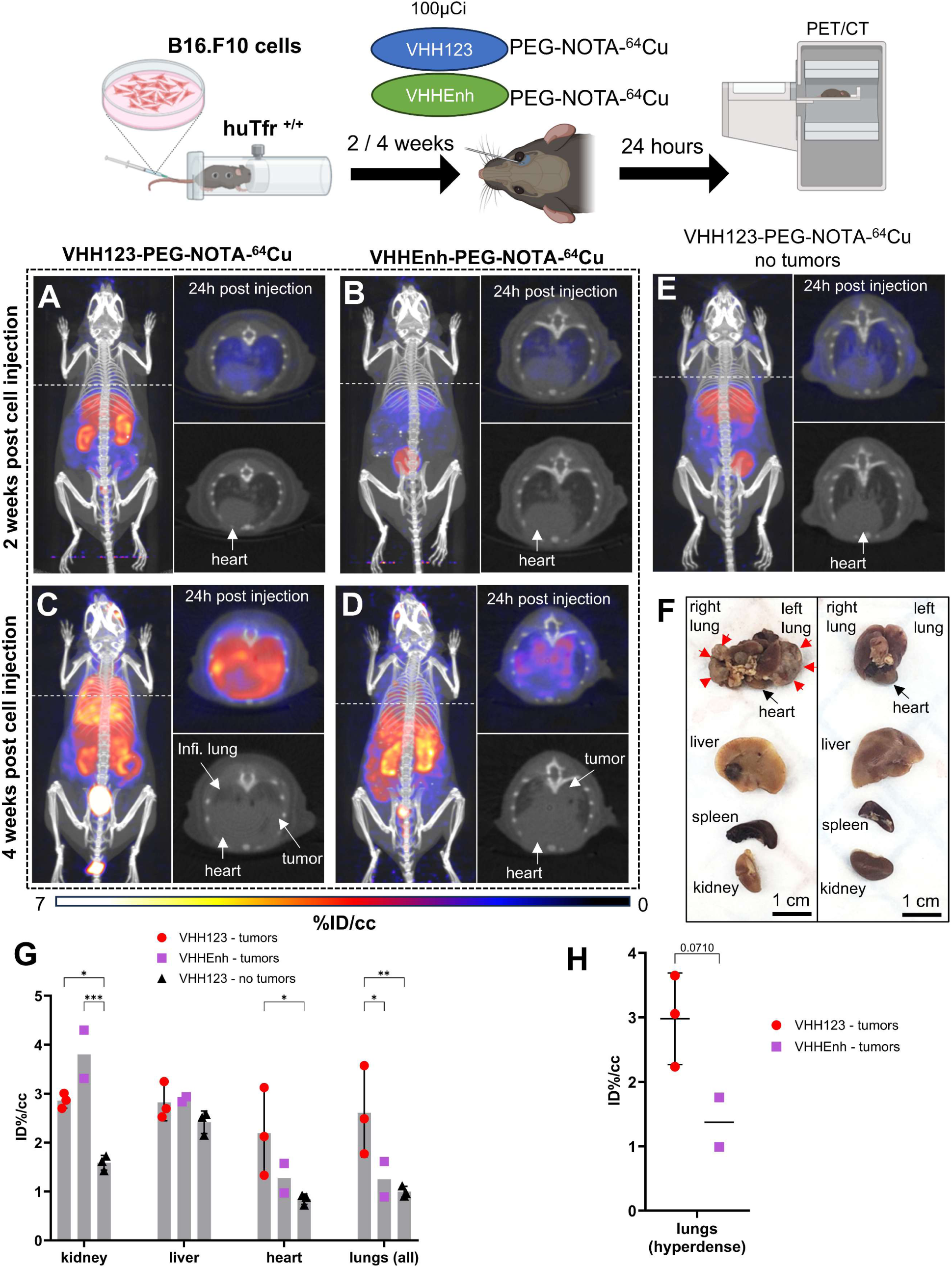
Top: cartoon depicting the experimental procedure: 5 × 10^4^ B16.F10 mouse melanoma cells were transfused to huTfR^+/+^ mice by tail-vein injection. 2 and 4 weeks later, the mice were injected with 3.7 MBq (100µCi) of either V_H_H123-PEG(20kDa)-NOTA-^64^Cu or V_H_HEnh-PEG(20kDa)-NOTA-^64^Cu radiotracers before PET/CT imaging. **A-E.** Maximum Intensity Projections (MIP) and lung traverse sections of one representative mouse out of three for each experimental condition at the 2 weeks timepoint post B16.F10 cell transfusion (except for E: no B16.F10 cells were injected). Images were acquired 24 hours post radiotracer injection. PET intensity scale is displayed on the right (%ID/cc). **A.** MIP of a huTfR^+/+^ mouse imaged with V_H_H123-PEG-NOTA-^64^Cu 2 weeks post B16.F10 cell transfusion. **B.** MIP of a huTfR^+/+^ mouse imaged with V_H_HEnh-PEG-NOTA-^64^Cu 2 weeks post B16.F10 cell transfusion. **C.** MIP of a huTfR^+/+^mouse imaged with V_H_H123-PEG-NOTA-^64^Cu 4 weeks post B16.F10 cell transfusion. **D.** MIP of a huTfR^+/+^mouse imaged with V_H_HEnh-PEG-NOTA-^64^Cu 4 weeks post B16.F10 cell transfusion. **E.** MIP of a huTfR^+/+^mouse imaged with V_H_H123-PEG-NOTA-^64^Cu that did not receive any tumor cells. **F.** Photographs of dissected organs from huTfR^+/+^ mice euthanized at 4 weeks post B16.F10 cell infusion and 96 hours post radio-tracer injection (left) and from control mice that received no cells 96 hours post radio-tracer injection (right). RL: right lung, LL: left lung, H: heart, Li: liver, Sp: spleen, Ki, kidneys. Organs are from the same respective mice as shown in C. Red arrows delimit necrotic and hyperdense tumors growing out of the right and left lung. **G.** ROI analysis of images acquired from mice as shown in panels C, D and E. Each dot represents the mean ID%/cc of a specific ROI for one mouse. Error bars show SD. No error bars are shown for the V_H_HEnh – tumor group as n=2 (one mouse died before imaging). **H.** ROI analysis of hyperdense lung tissue as visualized by CT on images acquired from mice as in panels C and D. Each point shows the mean ID%/cc of one mouse. Error bars show SD. No error bars are shown for the V_H_HEnh – tumor group as n=2.

### ^89^Zr-conjugated VHH188 binds to TfR at the blood-placenta barrier as observed by PET/CT

Pregnancy presents a unique situation akin to a transplant setting. Maternal Tf is taken up at the blood-placenta-barrier (BPB) to deliver iron to the developing embryo. We asked whether the V_H_H188 nanobody would detect TfR expressed by syncytiotrophoblasts-I at the BPB. C57BL/6 females were mated with a huTfR^+/+^ male or with a C57BL/6 male as a control. Embryos will thus be heterozygous for the presence of the two TfR isoforms. We hypothesize that proper mouse and human TfR homodimers will be formed, in addition to the possible formation of the interspecific hybrid TfR heterodimer. Two weeks post-fertilization, the pregnant females were injected retro-orbitally with V_H_H188-PEG-DFO-^89^Zr. We observed rapid uptake of the V_H_H188-PEG-DFO-^89^Zr radiotracer in the individual placentas of the huTfR^+/−^ embryos (**Fig 7A**). Wild-type C57BL/6 placentas were barely detectable in females crossed with a C57BL/6 male, showing a much weaker signal (**Fig 7B**). This difference is significant through ROI analysis (**Fig 7C**). We attribute the low but detectable signal for V_H_H188 to possible cross-reactivity to the mouse TfR when expressed at the very high levels on syncytiotrophoblast-I: placental tissue expresses a much higher than average level of Tfr1^23, 38^. Post-euthanasia dissection and separation of the placenta from the embryo was done, followed by a separate round of PET/CT imaging of the extirpated uterus and the embryos it contained. This confirmed that the intense V_H_H188-PEG-DFO-^89^Zr signal originates from the placenta and not from the embryo (**Fig 7D and E**). V_H_H188-PEG-DFO-^89^Zr thus detects huTfR^+/−^ placental tissue. Notwithstanding intense labeling of the placenta, we did not see a signal that corresponds to free ^89^Zr in either embryos or in the pregnant female. This results therefore departs from what was seen for any of the other imaging experiments, all of which showed release of ^89^Zr when the appropriate target TfR was expressed. Of note, the signal in the placenta was already quite intense at 1h post-injection of the radiotracer (**Fig 7F**).

**Figure 7.**
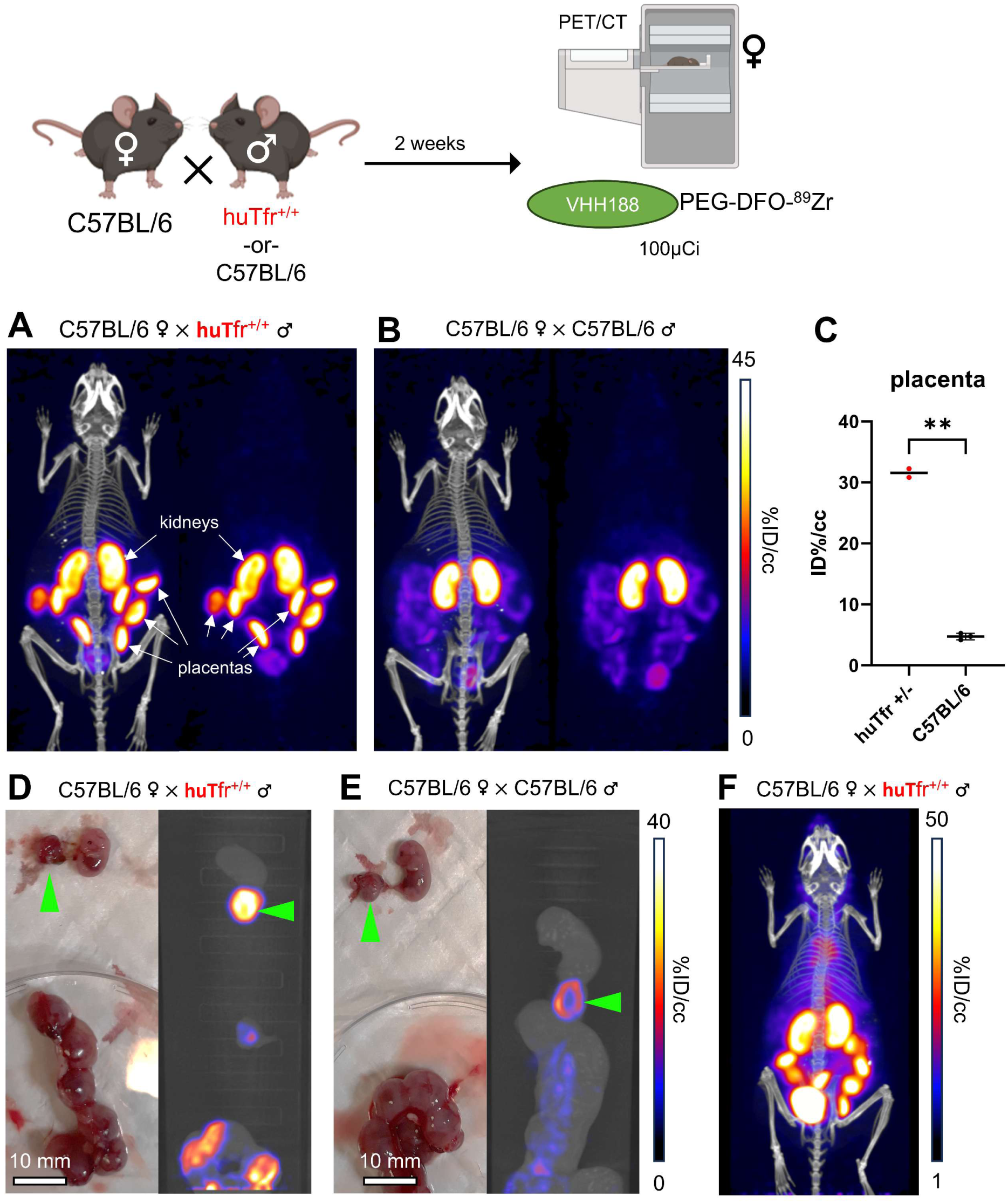
Top: cartoon depicting the experimental procedure: 8–12-week-old C57BL/6 females were mated in pairs with a single 12-week-old huTfR^+/+^ or C57BL/6 male. 2 weeks post fertilization (confirmed by observing vaginal plugs), the females are injected retro-orbitally with 3.7 MBq (100µCi) of V_H_H188-PEG(20kDa)-DFO-^89^Zr. **A.** PET/CT (left) and PET (right) maximum Intensity Projections (MIP) of one female carrying huTfR^+/−^ embryos, imaged 24 hours after injection of the radiotracer. **B.** PET/CT (left) and PET (right) MIP of one female carrying C57BL/6 wild-type embryos, imaged 24 hours after injection of the radiotracer. PET intensity scale for A and B is displayed on the right (%ID/cc). **C.** ROI analysis of images acquired from mice as shown in panels A and B. Each dot represents the mean ID%/cc of the placenta for one mouse. Error bars show SD. No error bars are shown for the huTfR^+/−^ condition as n=2 (2 out of 4 females had plugs 2 weeks prior but were not gestating at time of imaging). **D.** Left side panel: photograph of dissected embryos from the euthanized female carrying huTfR^+/−^ embryos, 72 hours post radiotracer injection. Right side panel: medial PET/CT section of a 50 mL tube containing the dissected embryos as shown on the left-panel. The green arrows highlight the placenta of one embryo. **E.** Left side panel: photograph of dissected embryos from the euthanized female carrying wild-type C57BL/6 embryos, 72 hours post radiotracer injection. Right side panel: medial PET/CT section of a 50 mL tube containing the dissected embryos as shown on the left-panel. The green arrows highlight the placenta of one embryo. PET intensity scale for D and E is displayed on the right (%ID/cc). **F.** PET/CT MIP of one female carrying huTfR^+/−^ embryos, imaged 1 hour after injection of the radiotracer. PET intensity scale is displayed on the right (%ID/cc). Experiment performed twice. Total of n=2 mice in B6 x huTfR^+/+^ group and n=3 in B6 x B6 group.

## Discussion

The ability to track the fate of transplanted cells non-invasively over time is a useful asset for several applications. These include the monitoring of tumor growth in response to various forms of therapy in a pre-clinical setting, or to follow the fate of transplanted cells of hematopoietic origin. Methods currently in use include luminescence-based approaches, in which the transplanted cells are engineered to express a suitable reporter, such as a luciferase ^1, 39^. Luminescence methods do not provide cellular resolution.

Alternatively, to achieve single cell resolution, fluorescence-based methods have been applied in multi-photon microscopy ^2^, but this typically involves invasive surgical interventions to expose the target tissue or organ, because absorption by surrounding tissue limits the depth of penetration of the excitation beam and emitted fluorescence.

Non-invasive methods such as NMR lack the specificity to detect particular cell populations, although tumors that exceed a certain size are readily visualized ^3^. Single photon emission computed tomography (SPECT) and positron emission tomography (PET) can detect specific cell populations based on the use of radiolabeled ligands that recognize them, but lack single cell resolution. The most widely used radioisotopes for SPECT include ^123^I and ^99^Tc, with half lives of ∼13 hours and ∼6 hours, respectively ^40^. Radiolabeled immunoglobulins used as imaging agents show excellent specificity, but their molecular mass (∼150 kDa) impedes efficient tissue penetration and imposes a long circulatory half-life. When using immunoglobulin-based imaging agents for PET, the use of long-lived radioisotopes such as ^89^Zr (t_1/2_=∼3.3 days) is therefore indicated.

The introduction of nanobodies for SPECT and PET overcomes many of the limitations of immunoglobulin-based imaging agents. The small size of nanobodies improves tissue penetration and drastically reduces the circulatory half-life of nanobody-based imaging agents. Free nanobodies are typically excreted via the kidneys and have a circulatory half-life of ∼30 minutes, versus a half-life (in mice) of several days for intact immunoglobulins ^41^. This means that unbound nanobodies are rapidly cleared from the circulation to yield a much-improved signal-to-noise ratio. Conjugation of a PEG_20kDa_ moiety further improves contrast by reducing somewhat the renal clearance rate of the nanobodies ^42^. Chemo-enzymatic methods for the modification of nanobodies allow their site-specific and reproducible modification with substituents of interest, including the installation of chelators for radiometals such as ^89^Zr and ^64^Cu, as well as click handles for site-specific covalent modification with ^18^F ^8^.

For detection of tumors, xenografts in immunocompromised mice are widely used in combination with tumor-specific imaging agents. Typically, they are based on antibodies that recognize human-specific surface markers ^43^. Methods for the detection of tumors of mouse origin in their natural host are few and far between. It is for this reason that we explored the possibility of using a congenic pair of mice in combination with nanobodies that distinguish the congenic marker, in this case the transferrin receptor (TfR). We show that a combination of anti-TfR nanobodies and mice that express either a wild-type (mouse) or a humanized TfR may be used to trace different cell types *in vivo* that express the TfR. To that end, we transferred hematopoietic bone marrow cells and B16.F10 melanoma cells into the respective mouse recipients. It is thus possible to track cells of mouse origin in humanized huTfR^+/+^ mice and humanized huTfR^+/+^ mouse bone marrow in C57BL/6 mice. Mice carrying huTfR^+/−^ fetuses showed a strong signal for VHH188 in the placenta, indicative of trans-endothelial transport of the imaging agent. Except for the TfR on syncytiotrophoblast-I, at 14 days of gestation no other embryonic tissues showed any accumulation of label.

The combination of human and mouse-specific anti-TfR V_H_Hs with mice that were engineered to express a TfR with the human TfR ectodomain constitutes a congenic pair of mice ideally suited for a variety of transplant experiments. The growth of tumors of mouse origin, when transplanted into huTfR^+/+^ mice, can be followed non-invasively, for example in response to various treatments. Conversely, human patient-derived tumor xenografts (PDX models) can be followed upon transplantation into mice without a requirement for genetic modification of the transplant, even if no tumor-specific antibodies are available.

Transplantation of C57BL/6-isolated tissues and tumor cells (-lines) in huTfR^+/+^ is straightforward, as both recipient and graft are of the same genetic background. The only required neo-epitope of the graft would be the ectodomain of the TfR, which we consider a negligible risk for immune rejection : the huTfR^+/+^ mouse expresses a chimeric TfR protein where only amino-acids 196-381 of the mouse TfR are replaced with the homologous amino-acids of the human TfR, that have a 74% pairwise identity by alignment using Clustal Omega ^22, 44^. The lack of an obvious PET signal in the CNS when imaging mice may be due to the fact that proteins that traverse the BBB, such as the TfR nanobody conjugates, accumulate in the CNS at a concentration far lower (around 1-5%) than that in blood plasma ^45, 46^. Furthermore, ^89^Zr radioconjugates are prone to radioisotope release post-internalization, which is the first step of transcytosis required for plasma proteins to traverse the BBB. Finally, both the ^89^Zr and ^64^Cu radioconjugates were PEGylated, which may impede transcytosis. While the use of non-PEGylated ^18^F radioconjugates might circumvent these issues, their short circulatory and isotopic half-life may not allow the visualization at adequate sensitivity of a signal in the CNS. Even so, gamma ray spectrometry on various organs from mice that received ^89^Zr-conjugated anti-TfR tracers confirmed accumulation of both nanobodies in the CNS of the appropriate genotype compared to control conditions (**Fig 4H**).

Release of free ^89^Zr and ^64^Cu by the nanobody conjugates was a surprising observation, unique to the anti-TfR V_H_Hs. For no other ^89^Zr labeled, PEGylated V_H_H have we seen such a striking and rapid release of free ^89^Zr. The osteophilic properties of ^89^Zr have been well-documented ^27, 28^ and confirmed in our hands (**Suppl. Fig. 8**). The signal generated by free ^89^Zr should therefore not be confused with the localization of anti-TfR nanobody to the spinal cord, bone marrow or skeletal elements more generally. Careful interpretation is required when examining accumulation of tracer in bone marrow when using anti-TfR-^89^Zr conjugates or any other internalizing ^89^Zr-labeled tracer. The fact that the observed release was unique to the anti-TfR V_H_Hs, seen only in the presence of the appropriate target, may also be consistent with a mechanism that specifically targets the DFO-^89^Zr chelate in compartments to which the TfR localizes. Perhaps release of Fe^+++^ from Tf requires not only acidic pH but also the presence of some as yet unidentified co-factor that can act on the ^89^Zr-DFO chelate as well. Imaging experiments performed on pregnant C57BL/6 mice that carry huTfR^+/−^ embryos may shed further light on this question. Notwithstanding the very strong accumulation of label seen in the placenta, which we ascribe to the presence of the huTfR at the surface of the embryonic syncytiotrophoblast-I, we did not see any sign of release of free ^89^Zr. Either the TfR upon internalization into the syncytiotrophoblast-I is never exposed to the low pH responsible for ^89^Zr release in other tissues, but still sufficiently low to allow release of Fe^+++^ from transferrin, or the hypothesized co-factor that mediates release of ^89^Zr from the DFO chelator is absent from the syncytiotrophoblast-I. We favor the former explanation, because it would allow delivery of Fe^+++^ to embryonic tissues and satisfy their demand for iron. Another possibility is that ^89^Zr is indeed released at the level of syncytiotrophoblasts-I, but would remain trapped at the interface between syncytiotrophoblasts-I and -II as it would then not be able to penetrate the fetal circulation via ferroportins ^23^. Release of free ^89^Zr from DFO after epitope-binding should also be a concern in the development of ^89^Zr-DFO radio-conjugates destined for clinical use.

The issue of the ‘free ^89^Zr signal pattern’, defined as an intense signal originating from long bone extremities, vertebrae and coxal bone, induced by ^89^Zr release may be traded for an increased signal in the liver by shifting to the use of ^64^Cu as a PET isotope, which is attributed to the passive release of ^64^Cu from NOTA – a phenomenon that has been well characterized in the literature ^35, 36^. The use of ^18^F-labeled anti-TfR V_H_Hs conjugated through click-chemistry avoids the complications of isotope release. However, ^18^F comes with its own drawbacks, such as its short half-life, necessitating less convenient and far more expensive synthetic routes for tracer production, and the typical accumulation of label seen in organs of elimination such as the gall bladder and gastro-intestinal tract when using TCO-tetrazine click-chemistry^34^.

In conclusion, this work demonstrates that the ubiquitously expressed TfR may be used as a cell marker to track virtually any cell-type of choice *in vivo*, provided they are transferred to a mouse model that expresses a different isoform of TfR. The huTfR^+/+^ mouse model has been deposited at the Jackson Laboratories and is therefore easily accessible (strain number 038212). Production and sortagging of nanobodies to generate PET tracers is a relatively simple process. These nanobodies and the huTfR^+/+^ model may thus benefit the field of *in vivo* imaging.

## Materials and Methods

### Production of nanobodies

cDNAs encoding V_H_H123 (anti-mouse TfR nanobody) and V_H_H188 (anti-human TfR nanobody) were cloned into a pHEN6 plasmid backbone that encodes the ‘GGLPETGGHHHHHH’ sortase A motif and histidine tag at the C-terminus of the expressed construct. WK6 *E. coli* were transformed with each plasmid vector and grown to saturation at 37°C in Terrific Broth (Millipore Sigma) prior to induction with 1mM IPTG and continued incubation overnight at 16°C. Extraction of each V_H_Hwas performed by osmotic shock as described ^47^. Purification of each V_H_Hwas achieved through Ni^2+^-NTA affinity chromatography followed by size-exclusion FPLC (Superdex 16/600 75pg, Cytiva).

### Sortase A mediated conjugation

10-30µM of V_H_H-GGLPETGG-His_6_ were incubated overnight at 8°C in the presence of 500µM – 1mM GGG-nucleophile, 30 µM of penta-mutant Sortase A-His_6_ (produced in-house as previously described ^48^, Addgene #51140), 2mM CaCl_2_ in 1mL total volume of PBS. 300µL Ni^2+^-NTA beads were then added to the reaction to capture Sortase A and unreacted V_H_H. The unbound fraction was desalted on a gravity-fed PD-10 size-exclusion column (Cytiva) to separate the V_H_H-conjugate from the excess of free GGG-nucleophile. Conjugation of each V_H_H was monitored by SDS-PAGE of each individual step of the reaction and by LC/MS of the purified product (QDa, Waters). Yields of sortase-mediated conjugations were > 75% or higher (total output protein mass vs. total input protein mass – measured by A280 using a NanoDrop, TermoFisher).

### Immunoprecipitation

Conjugation of V_H_H123 or V_H_H188 to biotin was performed by Sortase A conjugation and GGG-biotin as the nucleophile, as described above. The biotin-conjugated V_H_Hs were then incubated with 1 mg of streptavidin-coated paramagnetic beads (streptavidin Dynabeads T1, ThermoFisher) for 1 hour at 4°C in PBS. Unbound V_H_H-biotin was removed by washing the beads 3 times in lysis buffer: 1% NP-40, 150 mM NaCl, 20 mM Tris-HCL pH 7.4, Halt Protease Inhibitor 1X (ThermoFisher). HEK293 and B16.F10 cells were grown to confluency before detaching using Versene solution. Cells were washed in PBS to remove excess medium and FBS before lysis in 100 µL lysis buffer / 1 × 10^6^ cells. Lysates were incubated on a rotator for 30 minutes at 4°C before pelleting debris by centrifugation at 21,130 g for 5 minutes at 4°C on a benchtop centrifuge. Supernatants were then pre-cleared with 100µg / 300 µL of lysate of streptavidin Dynabeads for 1 hour at 4°C. The unbound fraction was removed after placing the tube on a magnetic rack, and incubated with 1 mg of V_H_H123-biotin or V_H_H188-biotin pre-coated Streptavidin Dynabeads (see above) overnight at 4°C. Beads were then washed 5 times in lysis buffer and then 2 times in PBS. Elution was done by boiling the beads in SDS-PAGE sample buffer. Eluted proteins were resolved by SDS-PAGE.

### LC/MS/MS analysis

Sample preparation and analyses were performed by the Taplin Mass Spectrometry Core at Harvard Medical School. Excised SDS-PAGE gel sections were cut into approximately 1 mm^3^ pieces. Gel pieces were then subjected to a modified in-gel trypsin digestion procedure ^49^. Gel pieces were washed and dehydrated with acetonitrile for 10 min. followed by removal of acetonitrile and lyophilization in a speed-vac. Gel pieces were then rehydrated with 50 mM ammonium bicarbonate solution containing 12.5 ng/µl modified sequencing-grade trypsin (Promega, Madison, WI) at 4°C. After 45 min., the trypsin solution was removed and replaced with sufficient 50 mM ammonium bicarbonate solution to cover the gel pieces. Samples were then placed in a 37°C room overnight. Peptides were recovered by removing the ammonium bicarbonate solution, followed by one wash with a solution containing 50% acetonitrile and 1% formic acid. The extracts were combined and dried in a speed-vac (∼1 hr). Samples were reconstituted in 5 - 10 µl of HPLC solvent A (2.5% acetonitrile, 0.1% formic acid) and were applied to a nano-scale C18 reverse-phase HPLC capillary column. Peptides were eluted with increasing concentrations of solvent B (97.5% acetonitrile, 0.1% formic acid) and were subjected upon elution to electrospray ionization and then injected into a Velos Orbitrap Pro ion-trap mass spectrometer (Thermo Fisher Scientific, Waltham, MA). Peptides were detected, isolated, and fragmented to produce a tandem mass spectrum of specific fragment ions for each peptide. Peptide sequences (and hence protein identity) were determined by matching protein databases with the acquired fragmentation pattern by the software program, Sequest (Thermo Fisher Scientific, Waltham, MA) ^50^. All databases include a reversed version of all the sequences and the data was filtered to between a one and two percent peptide false discovery rate. Subcellular localization annotation was retrieved by matching the Uniprot entry number of each protein with its Uniprot sub-cellular localization annotation, using the CellWhere database ^51^.

### ^35^S-Cysteine/Methionine labelling of cells

HEK 293T and B16.F10 cells were grown to 80% confluency in 2x T75 plates each before careful washing 2x with Cys/Met free DMEM media with 10% dialyzed FBS. Cells were then starved in Met/Cys-free medium at 37°C for 30 minutes with dialyzed FBS before replacing the medium with ^35^S-Cys/Met enriched medium (11 µCi/µL, EasyTag™ EXPRESS35S Protein Labeling Mix, Perkin Elmer) for 5 hours at 37°C. ^35^S-labelled cells were then processed for immunoprecipitation as described above.

### Flow cytometry

A dilution range of different V_H_H concentrations prepared in PBS 2% FBS, were incubated with 0.1 million CHO cells overexpressing either the human or mouse TfR for 30 min at 4 °C. The binding of V_H_Hs was next followed by a 30 min incubation at 4 °C with an anti-FLAG-iF647 antibody (A01811, Genscript, Piscataway, NJ, USA), diluted 1:500. Dead cells were stained with the viability dye eFluor™780 (1:2000; 65-0865-14, Thermofisher Scientific, Waltham, MA, USA) for 30 min at 4 °C. Flp-In™-CHO™ cells, used as unstained control and single stain controls, were used to determine the cutoff point between background fluorescence and positive populations. UltraComp eBeads™ Compensation Beads were used (01-2222-42, Thermofisher Scientific, Waltham, MA, USA) to generate single stain controls for the anti-FLAG-iF647 antibody. The data was acquired by using an Attune Nxt flow cytometer (Thermofisher Scientific, Waltham, MA, USA) and analyzed by FCS Express 7 Research Edition.

### Mice

C57BL/6 mice were purchased from Jackson Laboratories (strain 000664) and huTfR^+/+^ mice were provided by the groups of M. Dewilde and B. De Strooper (VIB and KU Leuven, Belgium) and bred in-house. The huTfR^+/+^ strain has been deposited at Jackson Laboratories (strain 038212). Mice were housed and handled according to the institution’s IACUC policy #00001880 – with a 12h day/night cycle, min/max temperature of 20/23.3°C and water/food access *ad libitum*. Unless otherwise specified in figure legends, mice used in the experiments were between 8-12 weeks old. Both male and female mice were included in our experimental groups, at an approximate 50/50 percent ratio.

### Transplantation of bone marrow

Femora and tibiae from 6–8-week-old C57BL/6 mice were flushed using a syringe equipped with a 23G needle to harvest bone marrow in Iscove’s Modified Dulbecco’s Medium (IMDM). Cells were pelleted and resuspended in PBS, re-pelleted and then resuspended in 5 mL RBC lysis buffer (15mM NH_4_Cl, 1mM KHCO_3_, 1µM disodium EDTA in water) and left for 10 minutes at room temperature. Cells were then washed twice in PBS to remove lysed RBCs prior to injection of 1.5×10^6^ cells into a lethally irradiated (2 x 5Gy, 4 hours apart) recipient huTfR^+/+^ mouse. Recipient mice were kept in individual restraining chambers during irradiation to maintain a uniform total body irradiation.

### Implantation of B16.F10 lung metastases

B16.F10 melanoma cells were grown to confluency, then detached with Versene solution 1X, washed twice and resuspended in sterile PBS at a concentration of 2.5 × 10^5^ cell/mL. 5 × 10^4^ cells were transfused *i.v.* by tail-vein injection in huTfR^+/+^ mice. Lung metastases began to appear ∼ 3-4 weeks later, as confirmed by CT imaging of the lung and necropsy at the end of the experiment. Control mice that beared no tumors received PBS instead by tail-vein injection.

### Radioactive conjugate preparation

Conjugation to ^89^Zr was based on previously published methods ^24^. V_H_Hs were conjugated to GGG-Deferoxamine (DFO)-Azide by Sortase A transpeptidation (see above). The V_H_H-DFO-Azide conjugate was then incubated with a 5-fold molar excess of polyethylene-glycol_20kDa_-dibenzylcyclooctyne (PEG_20kDa_-DBCO) overnight at 4°C on a shaker in PBS (pre-treated with Chelex beads in order to remove divalent cations) in order to generate V_H_H-PEG_20kDa_-DFO through click-chemistry. Completion of the reaction was assessed by SDS-PAGE. For radio-labelling, a stock solution of 129.5 MBq (3.5 mCi) of ^89^Zr^4+^ in a 1M oxalate solution (purchased from the Madison-Wisconsin University Cyclotron Lab) was adjusted to a pH of 6.8-7.5 with a 75% (vol/vol) of 2M Na_2_CO_3_ and 400 % (vol/vol) of 0.5M HEPES buffer, pH 7.5. 37 MBq (1mCi) of pH-adjusted ^89^Zr was then mixed with 100µg of V_H_H-PEG_20kDa_-DFO for 1h at room temperature, followed by removal of unbound ^89^Zr using a PD-10 gravity desalting column (Cytiva) pre-equilibrated with Chelex-treated PBS. The column was eluted in fractions of 600µL, the activity of which was measured using a dose calibrator (AtomLab 500, Biodex). The fraction corresponding to the peak activity (typically fraction 6 with an activity of typically 37 kBq/µL (1µCi/µL)) was used for injection. The free ^89^Zr remaining in the desalting column was typically < 10% of input: 37 kBq (100 µCi) suggesting a radioelement chelation efficiency of ∼90%. For conjugation of ^64^Cu, GGG-NOTA-Azide was conjugated to the V_H_H through sortase A transpeptidation (NOTA: 2,2′,2”-(1,4,7-triazacyclononane-1,4,7-triyl)triacetic acid). The resulting conjugate was then further conjugated to PEG_20kDa_ by click-chemistry as described above. For radio-labelling, a stock solution of 1.48 GBq (40mCi) of ^64^CuCl_2_ (purchased from the Madison Wisconsin University -Cyclotron Lab) was mixed with 150µg of V_H_Hconjugate in PBS for 1 hour at room temperature on a shaker. Unbound ^64^Cu was removed from the mixture by passage onto a PD-10 gravity-fed desalting column (Cytiva) pre-equilibrated with PBS. The elution of the column, peak activity measurements and injection doses were the same as with ^89^Zr, as described above.^18^F-based V_H_H conjugates were ready for injection post click-chemistry of the tetrazine-conjugated V_H_Hs with ^18^F-TCO at the Molecular Cancer Imaging Facility at Dana-Farber Cancer Institute, Boston, M.A. (Fig. 2 and suppl. Fig 1 and 10). Synthesis of ^18^F-TCO is described in supplementary methods. In short, V_H_H123-tetrazine (25 μL, 230 μM) was diluted with 668 μL of 1x PBS. To this solution, 754.8 MBq (20.4 mCi) of ^18^F-TCO were added in 37 μL of EtOH. The reaction mixture was placed on a Thermomixer at 300 rpm at 25 °C. V_H_H188-tetrazine (21 μL, 275 μM) was diluted with 843 μL of 1x PBS and reacted with 943.5 MBq (25.5 mCi) of ^18^F-TCO in 45 μL of EtOH. Progress of click reactions was monitored by spotting iTLC-SG strip (Agilent, SGI0001) with 0.5 μL of reaction mixture at 5 and 10 min until about 20% of clicked product was detected in the mixture (10 min). For purification, 200 μL of TCO-agarose slurry (50% slurry in 20% EtOH, Click Chemistry Tools, 1198-5) was added to the top of a pre-equilibrated PD-10 column. The column was then washed with 30 mL of sterile 1x PBS. Each ^18^F-TCO-tetrazine-V_H_H click reaction mixture was loaded to a column. When the solution reached bed level, 1.6 mL of 1x PBS was added and the ^18^F-radiolabeled V_H_H was then eluted with 2×1 mL fractions of PBS. For ^18^F-V_H_H123, the first 1 mL fraction contained 13.98 MBq (378 μCi) of product and the second 1 mL fraction measured 82.88 MBq (2.24 mCi). The first fraction was discarded, and after checking pH and radiochemical purity of the second fraction (iTLC-SG, **Supplemental Figure 11**), the final formulation contained 78.07 MBq (2.11 mCi) of ^18^F-V_H_H123 in 0.95 mL of 1x PBS. For ^18^F-V_H_H188, the first 1 mL fraction contained 30.97 MBq (837 μCi) of product and the second 1 mL fraction measured 138.38 MBq (3.74 mCi). The first fraction was discarded, and after checking pH and radiochemical purity of the second fraction (iTLC-SG, **Supplemental Figure 11**), the final formulation contained 128.02 MBq (3.46 mCi) of ^18^F-V_H_H188 in 0.91 mL in 1x PBS. Each were used immediately after synthesis for radio-imaging. All final preparations of conjugates were confirmed to be ∼pH 7.4 by testing with pH paper.

### PET/CT imaging

Mice were anaesthetized using 2.0% isoflurane in O_2_ at a flow rate of ∼1 liter per minute. For all radiotracers, 1.85 to 3.7 MBq (50 to 100 µCi) of radiotracer was injected retro-orbitally (typically in a 50 to 100 µL volume, depending on final activity/mL of radiotracer). PET/CT images were acquired using a G8 PET/CT machine (Sofie biotech – Perkin Elmer) with a 10-minute PET signal detection window at several timepoints post-injection: 1-2h, 12h, 24h then every 24 hours until 96 hours post-injection or until the radio-isotope had decayed. Each PET acquisition was followed by a 1.5-min CT scan. Raw acquired images were processed by the manufacturer’s automatic image reconstruction software to generate DICOM files. Images were then visualized, rendered and analyzed using VivoQuant 3.5 software, patch 2 (Invicro). For ID%/cc, images were produced using ID%/g scales that were converted to ID%/cc by postulating that 1 mL of tissue = 1 g. Mice were kept anesthetized by continuous inhalation of 2.5 % isoflurane during the acquisition of PET/CT images.

### Ex vivo measurement of activity

Mice injected with ^89^Zr-radiolabelled conjugates were euthanized by CO_2_ inhalation. No capillary depletion was performed. Organs were harvested post-mortem and collected in pre-weighed 5mL assay tubes. Each organ was weighed prior to gamma-counting using a Packard E5003 instrument.ID%/g (injected dose percentage per gram of tissue) for each sample was calculated using the following formula: (activity of sample (MBq)/total injected activity (MBq))/sample weight (g) × 100. For whole blood, sample weight was calculated by considering that 1mL of whole blood = 1.06 g. Bone marrow was harvested by extensive flushing of harvested femurs with 4 mL total volume of PBS per femur. For comparing the activity of bone marrow vs emptied bone, weight of both sample types was considered as equal to 1, in order to generate a ID%/bone scale

### Antibodies

Mouse monoclonal anti-TfR IgG (mouse and human cross-reactive), clone H68.4, Abcam catalog #ab269513

### Cells

HEK 293 and B16.F10 cells were provided by the laboratory of Dr. Stephanie Dougan – Dana Farber Cancer Institute, Boston, MA, USA. B16.F10 cells were tested and found negative for presence of specific mouse pathogens.

### Statistics

For every comparison of one condition between two populations, unpaired two-tailed t-tests using Welch’s correction for non-equal standard deviations was performed. For every comparison of multiple conditions between two populations, unpaired two-tailed t-tests were performed, with Holm-Šídák’s correction for multiple comparisons. For every comparison of one condition across 3 or more populations, one-way ANOVA with Tukey’s correction for multiple comparisons was performed. For every comparison of two or more conditions across 3 or more populations, two-way ANOVA was performed, with Holm-Šídák’s correction for multiple comparisons by considering all comparisons. All these analyses were done using GraphPad Prism v. 10.2.3. When considering experiments containing repeats at different timepoints – comparisons were done only between populations within the same timepoint. For all graphs: p <= 0.05 : *, p < 0.01 : **, p < 0.001 : ***, p < 0.0001 : ****

## Acknowledgements

The authors thank the Molecular Cancer Imaging Facility at Dana-Farber Cancer Institute, Boston, M.A. for the conjugation of ^18^F to the VHH-tetrazine constructs and the Taplin Mass Spectrometry Core at Harvard Medical School, Boston, M.A. for the LC/MSMS analysis. T.B. acknowledges support of a Belgian American Educational Foundation post-doctoral fellowship and of a WBI. World fellowship from Wallonie-Bruxelles International. B.dS and M.D. acknowledges a Grand Challenges grant from the Vlaams Instituut voor Biotechnologie (VIB), Ghent, Belgium. MD & TJ acknowledge Interne Fondsen KU Leuven/Internal Funds KU Leuven for its financial support. BDS is grateful for Methusalem support from the KU Leuven.

## Author Contributions

All authors participated in the study design. T.B., C.C., S.O., P.S. T.J. and X.L. performed the experiments. X.L. synthesized and prepared the GGG-conjugates. H.M. and Y.L. performed the experiment described in supplemental figure 8. T.B. and H.P. wrote the manuscript.

## SUPPLEMENTAL DATA AND FIGURES

**Supplemental Figure 1.**
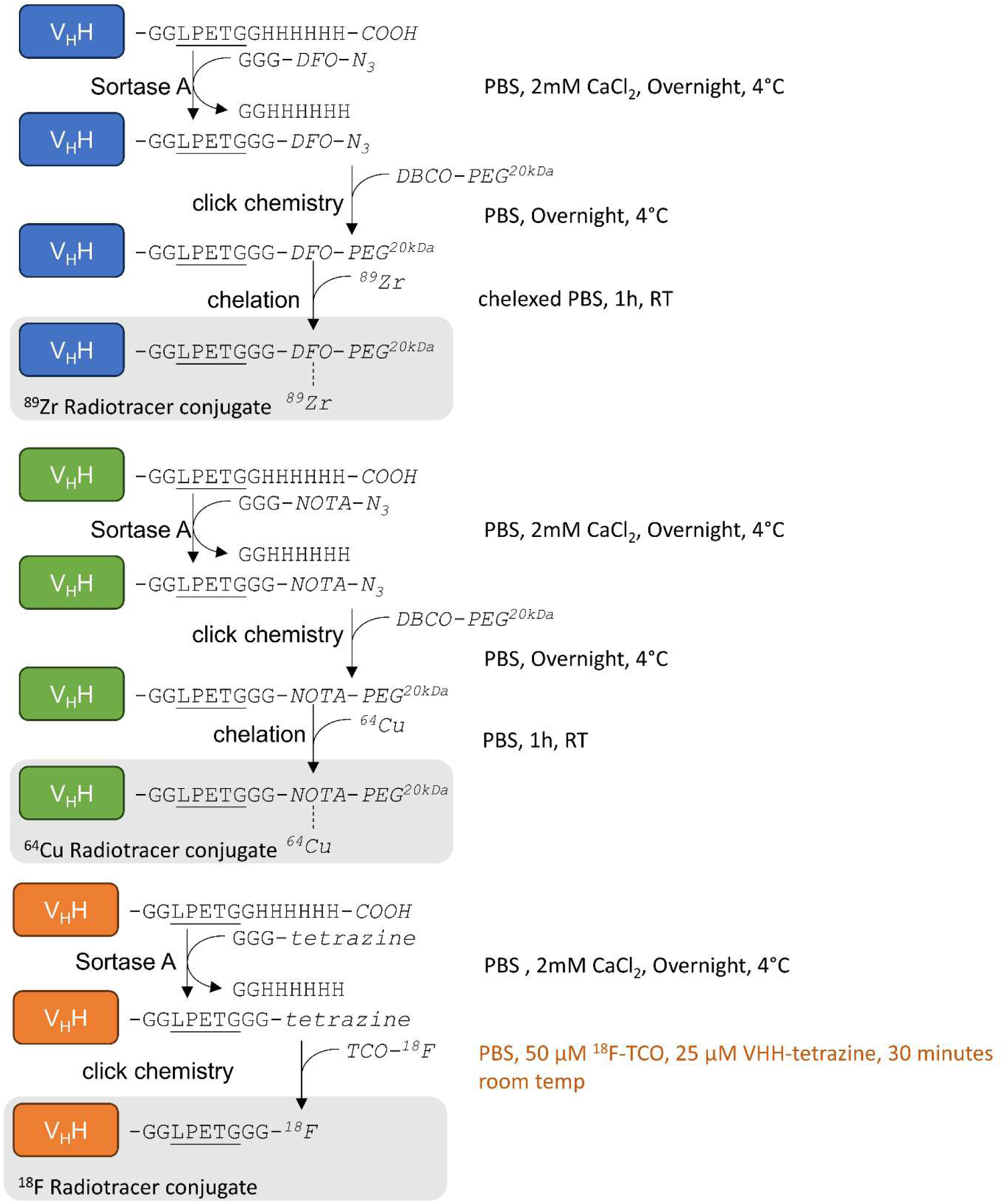
Schematic representation of each step required to generate ^89^Zr-based (blue), ^64^Cu-based (green) and ^18^F-based (orange) V_H_Hconjugates. Capital letters denote amino-acids, unless written in *italic* where they denote chemical groups or elements. Each reaction condition is noted on the right hand of each reaction. DFO: deferoxamine, NOTA: 2,2′,2”-(1,4,7-triazacyclononane-1,4,7-triyl)triacetic acid, PEG: poly-ethylene-glycol, TCO: trans cyclo-octene, DBCO: dibenzocyclooctyne. See methods for details.

**Supplemental Figure 2.**
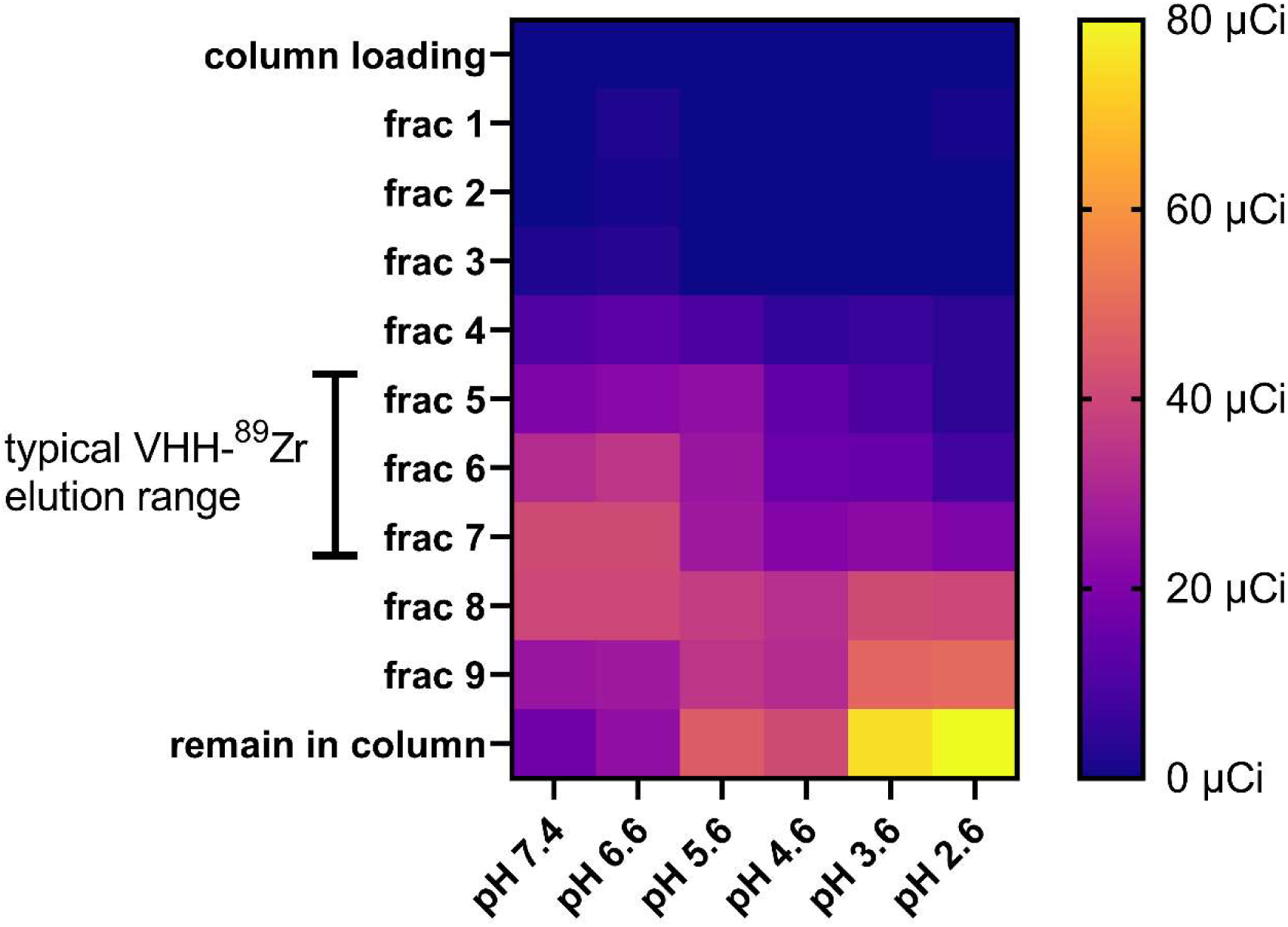
185 MBq (5mCi) of ^89^Zr-oxalate stock solution was obtained from the Cyclotron Lab at UW Madison, USA. Stock solution was put at a neutral pH of 7.5 by successive addition of 90% of stock volume of 2.0 M Na_2_CO_3_ and 400% stock volume of HEPES 0.5M. V_H_H123-PEG-DFO was conjugated to ^89^Zr at a neutral pH of 7.4 by adding 44.4 MBq (1.2mCi) of pH neutralized ^89^Zr to 120µg of V_H_H123-PEG-DFO in chelexed PBS, put in a microcentrifuge tube for 1 hour at room temperature on an agitator. The mixture was then split into 6 tubes (200 µCi each), and diluted 1/15 (v/v) in citrate-sodium phosphate buffer of varying pH: 7.4, 6.6, 5.6, 4.6, 3.6 and 2.6. The conjugation mixture was then immediately passed onto a PD-10 gravity size exclusion chromatography column (Cytiva) pre-equilibrated with citrate-sodium phosphate buffer of same pH used to dilute the VHH conjugate. Fractions of 600µL were collected by addition of citrate-sodium phosphate buffer of same pH and each fraction was measured for radioactivity using a dosimeter (AtomLab 500, Biodex). Shown is a heat map of measured radioactivity from each fraction collected. ‘column loading’: flowthrough from column displaced by application of the reaction mixture to the column. ‘remain in column’: residual radioactivity measured from the whole column post-elution. Brackets on the left indicate the typical elution fraction range of a VHH-radiometal conjugate post successful conjugation.

**Supplemental Figure 3-7.** Images for each individual repeat and timepoint of each PET/CT image panel shown in figures 3-7 (respectively) :

**PLEASE SEE ANNEXED FILE (large size)**

**Supplemental Figure 8:**
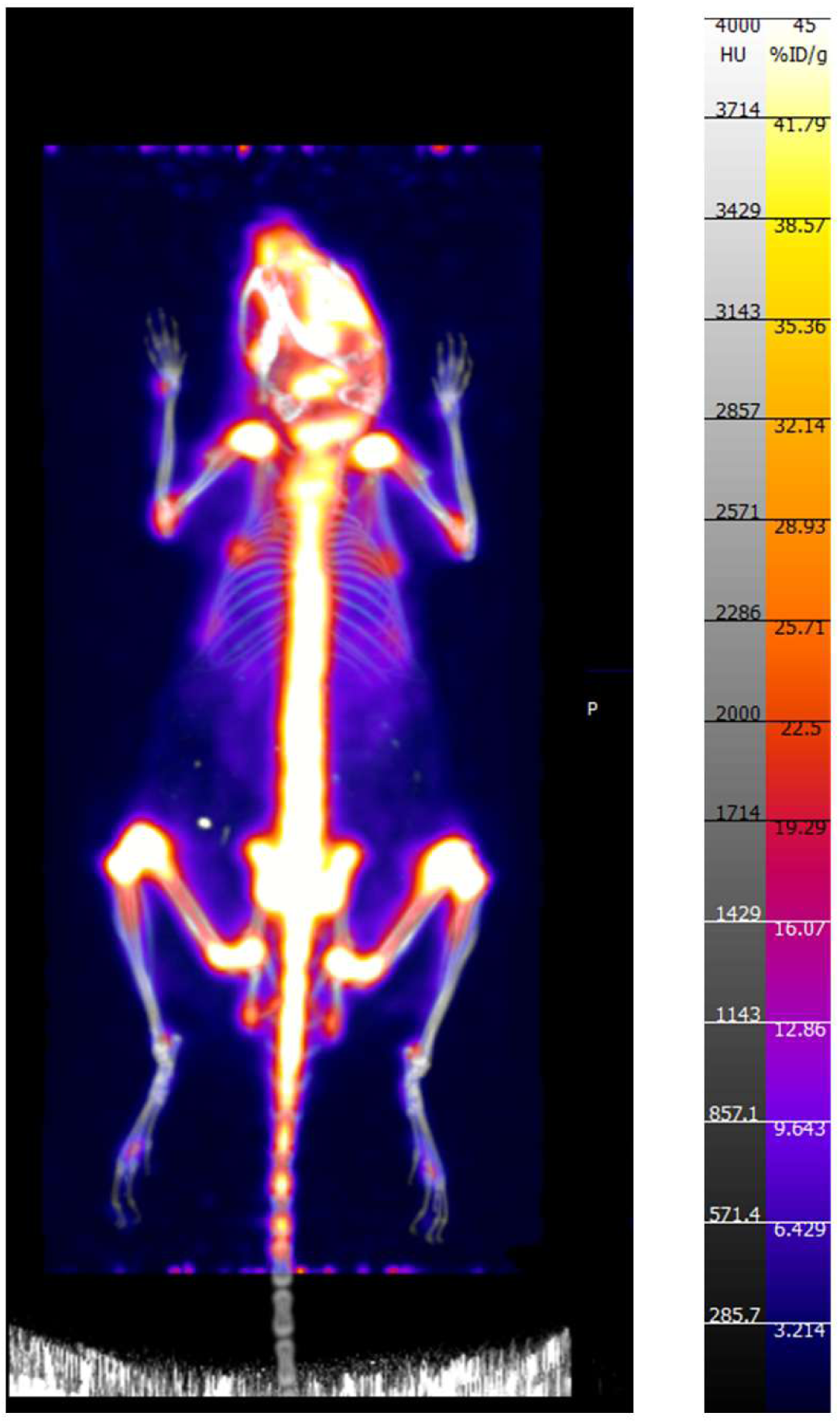
PET/CT of one female C57BL/6 mouse that received 2.775 MBq (75 µCi) of ^89^Zr as a free element (un-chelated/un-conjugated) alongside 75 × 10^8^ gold chiral nano-particles. This image exclusively shows the signal generated by the accumulation of free ^89^Zr in a mouse. The co-injection of gold nano-particles was done in relation to another work published by our group, but where this data was not included ^52^.

**Supplemental Figure 9:**
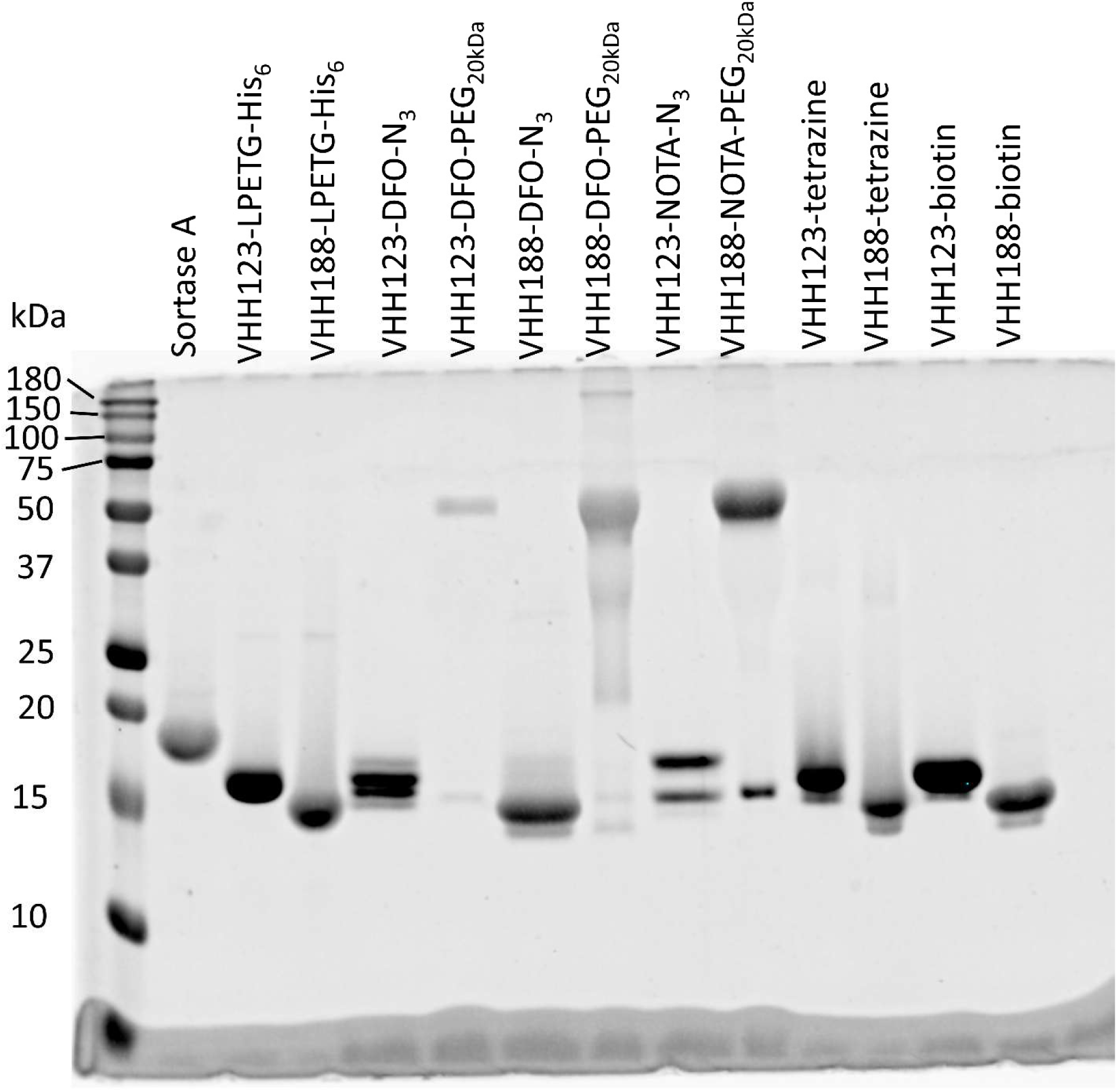
SDS-PAGE followed by Coomassie Blue stain, showing the individual constructs used throughout this work, as purified and concentrated post-sortase A transpeptidation and post DBCO-PEG_20kDa_ click-chemistry conjugation when performed. Sortase A expected mw: 17.8 kDa. For expected mw of other constructs, see Suppl. Figure 10.

**Supplemental Figure 10:**
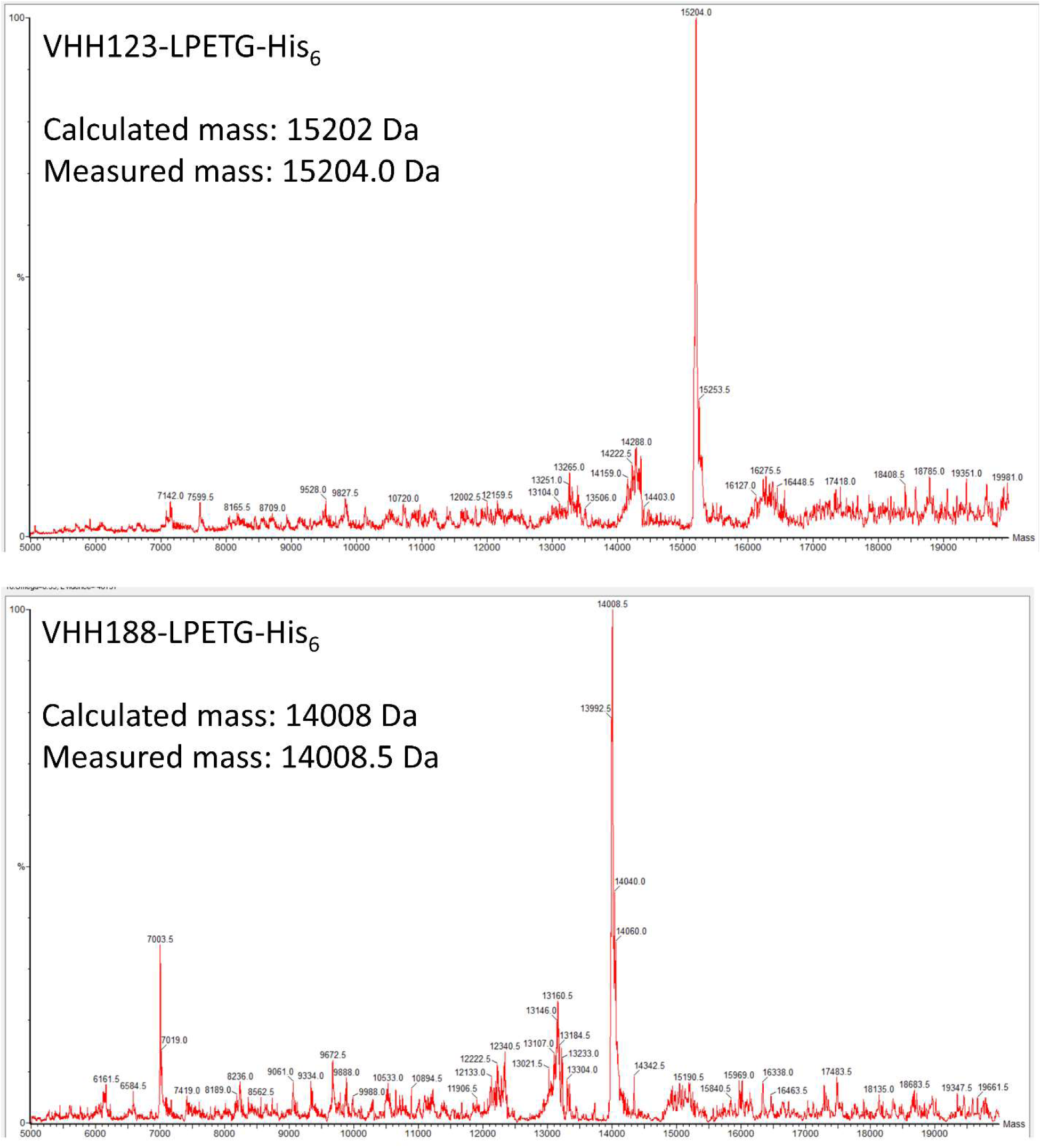
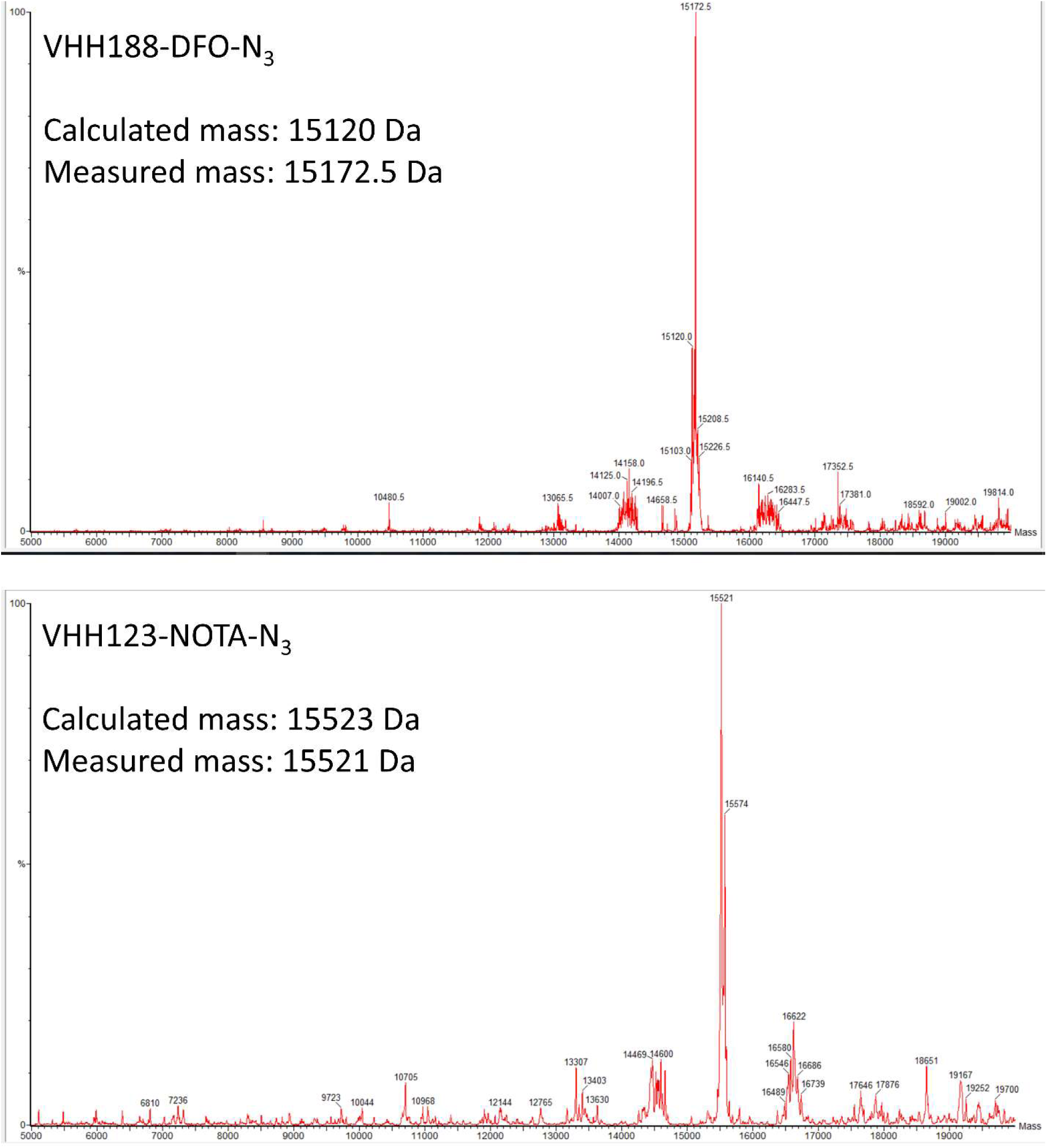
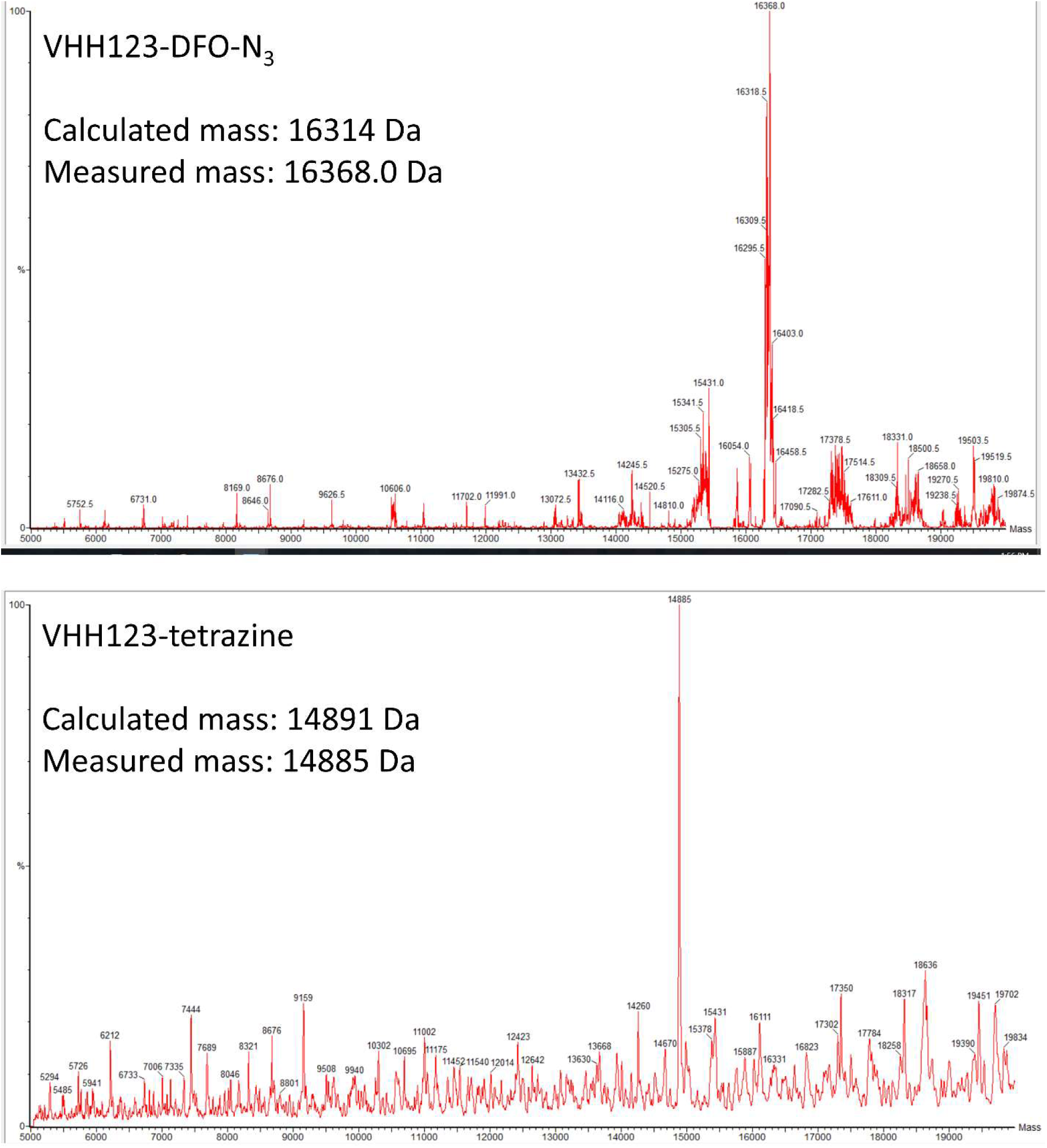
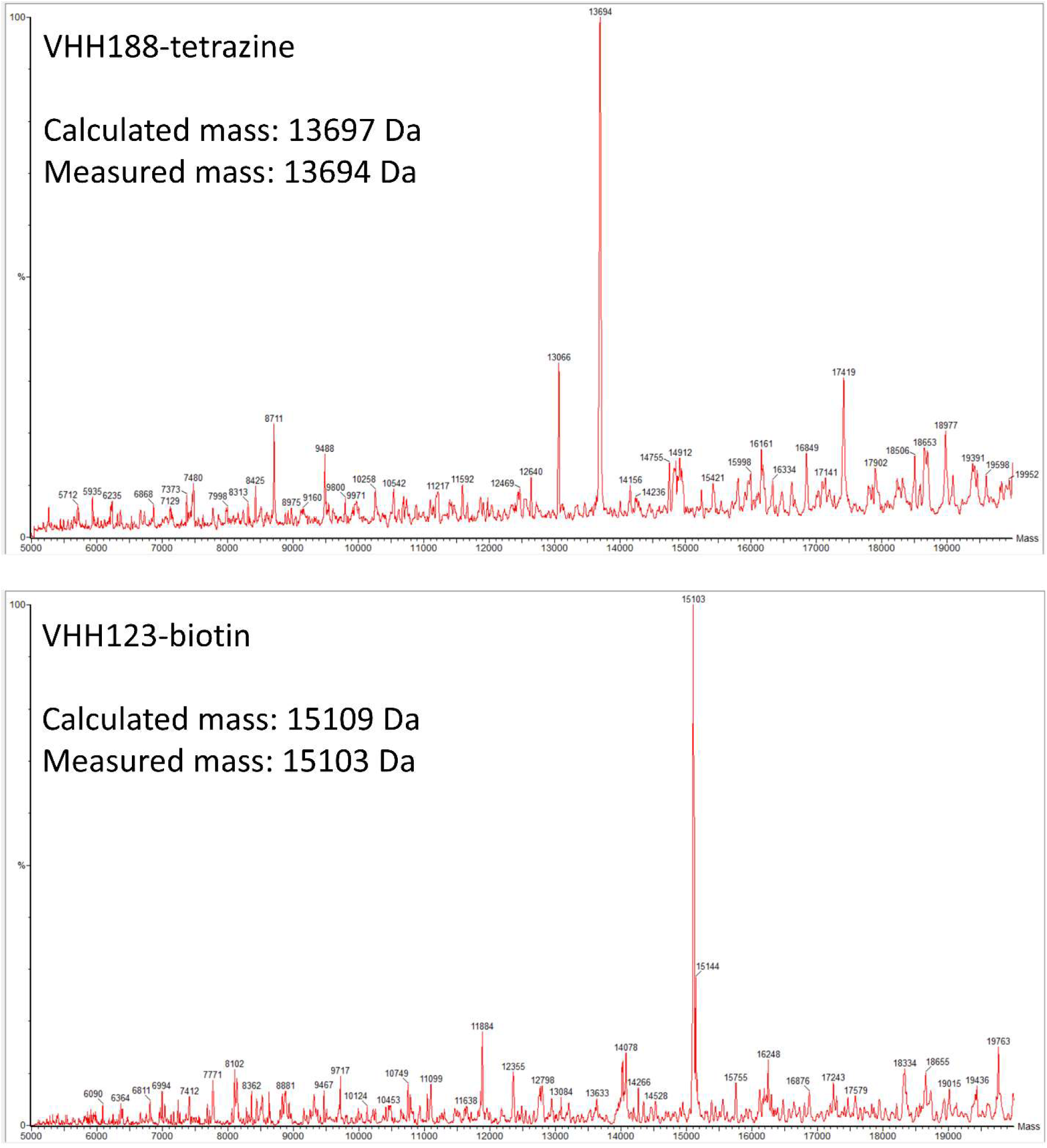
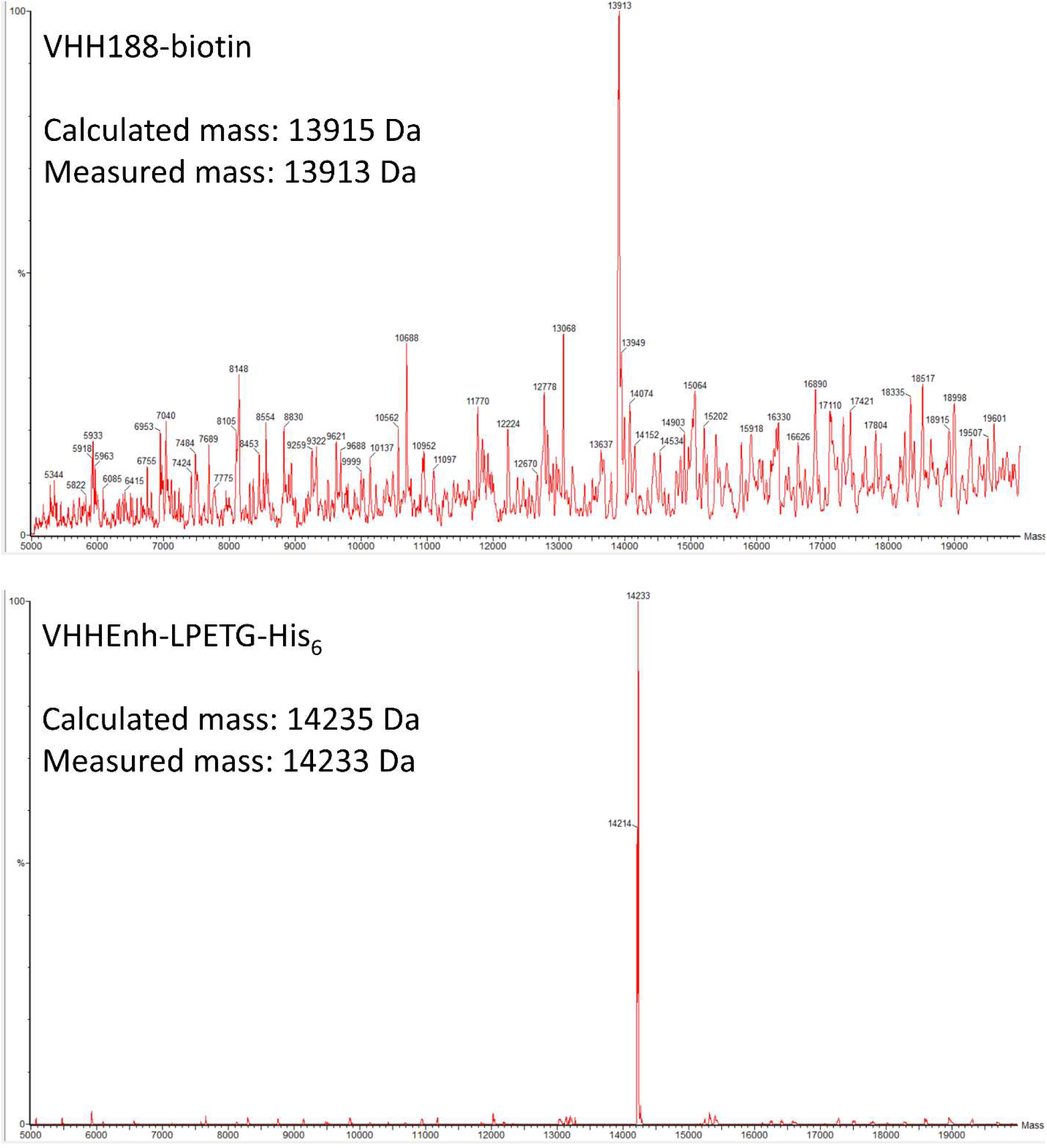
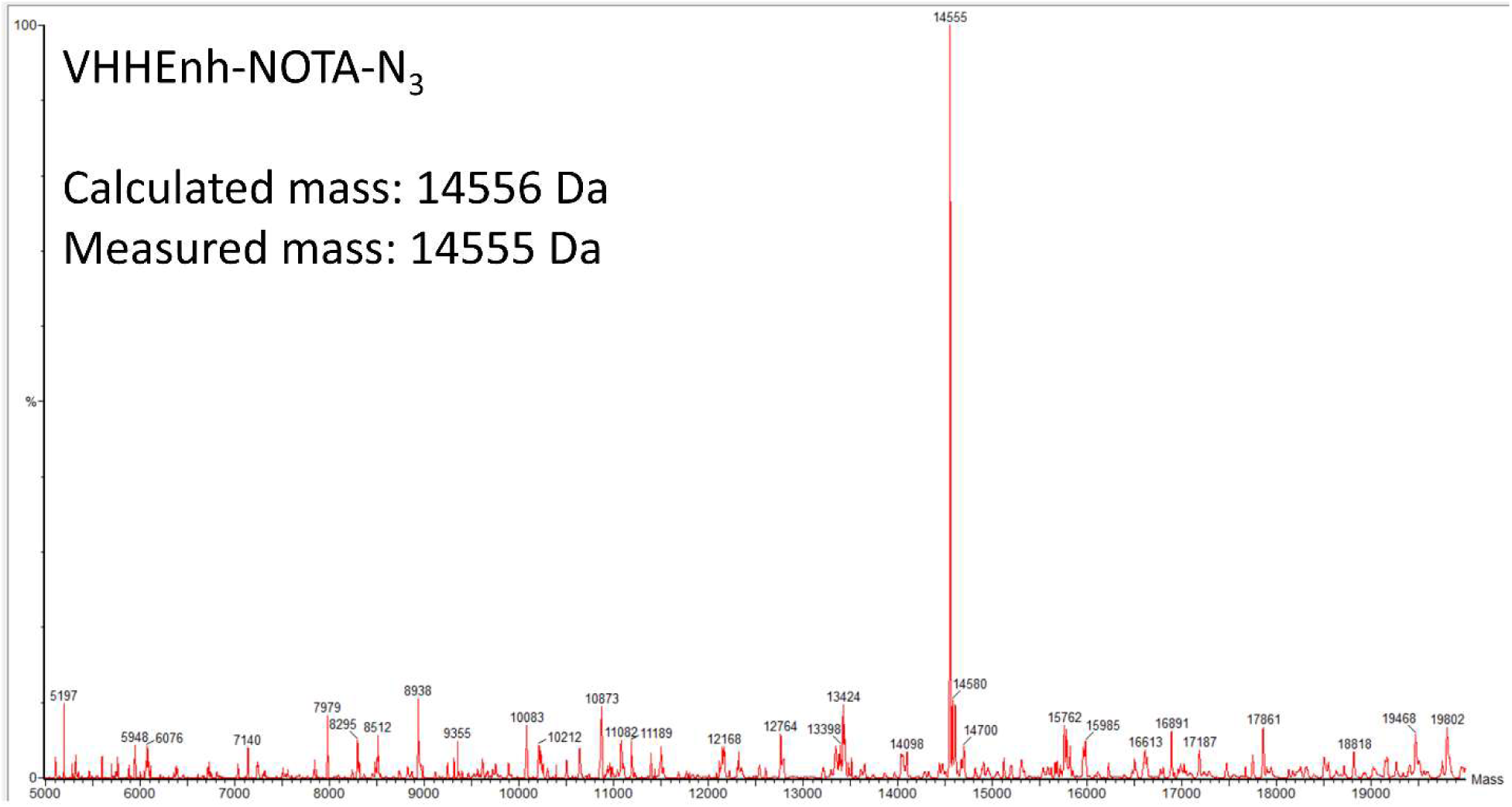
LC/MS mass measurements of each V_H_H construct used in this work. PEG_20kDa_ modified constructs are not shown as their masses cannot be precisely measured due to the fact that the DBCO-PEG_20kDa_ compound has an <u>average</u> mass of 20kDa.

**Supplemental Figure 11:**
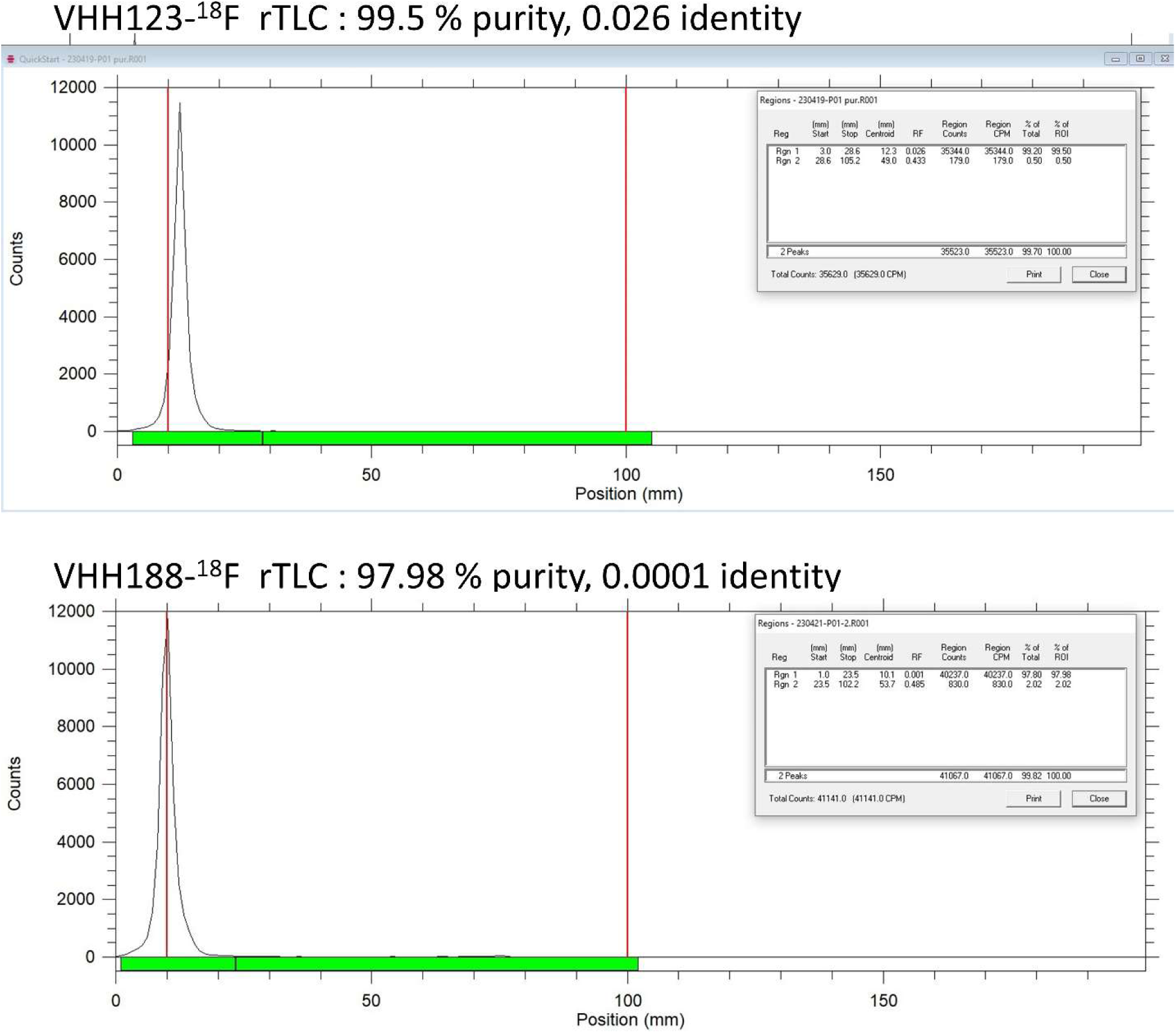
Radio-Thin Layer Chromatography QC data for VHH-^18^F constructs.

**Supplemental Figure 12.**
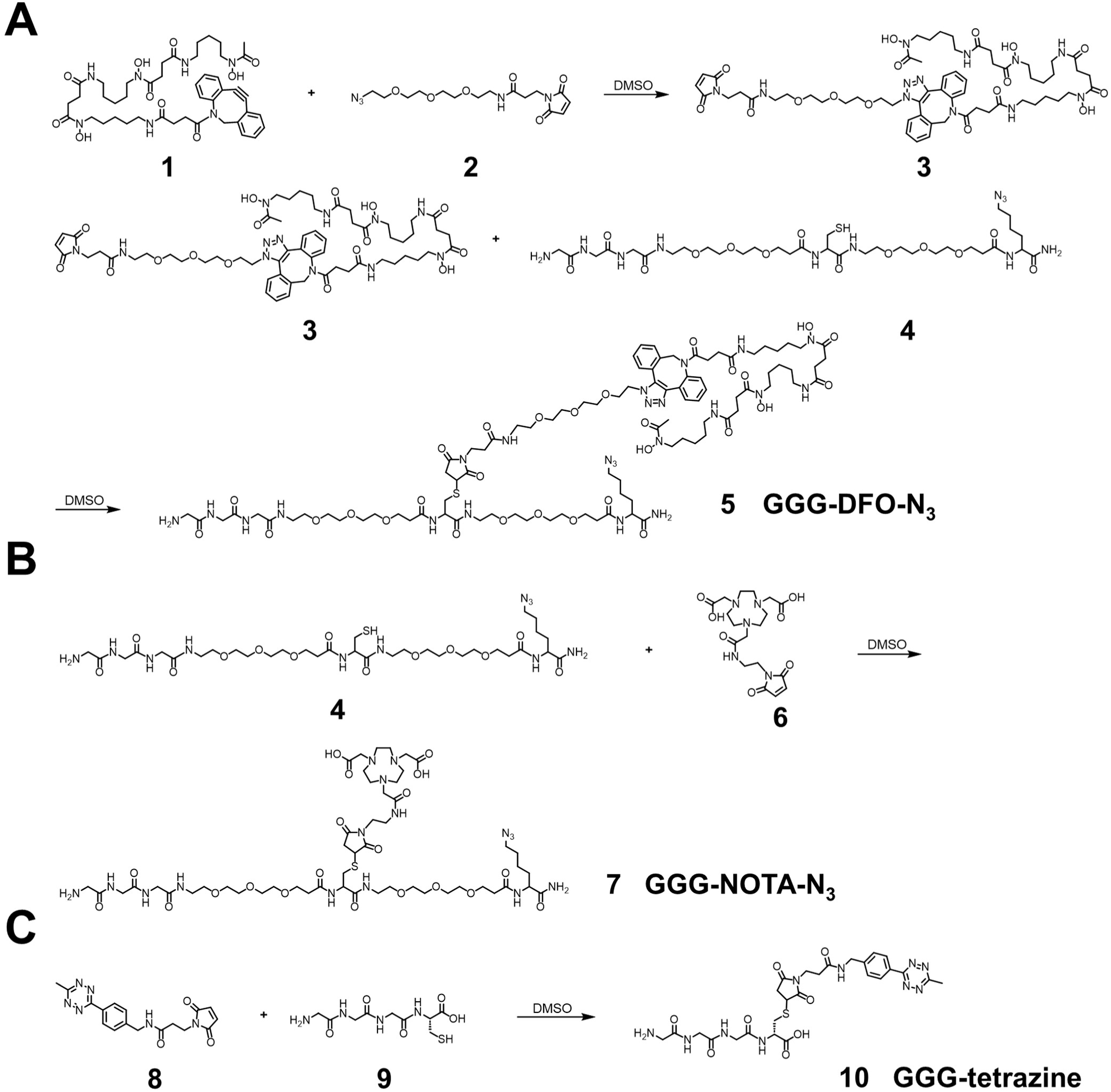
Synthetic schemes of labeling reagents. (A) Synthesis of a triglycine- and azido-modified deferoxamine (DFO) (GGG-DFO-N3) as a sortase-ready radioisotope zirconium-89 (89Zr) chelator. Polyethylene glycol (PEG) spacers were incorporated to enhance molecular flexibility and water solubility while minimizing potential interactions, such as steric hindrance, among different moieties. (B) Synthesis of a triglycine- and azido-modified NOTA (GGG-NOTA-N3) as a sortase-ready radioisotope Copper-64 (64Cu) chelator. (C) Synthesis of a triglycine-modified tetrazine as a sortase-ready click chemistry handle.

**Supplemental Figure 13.**
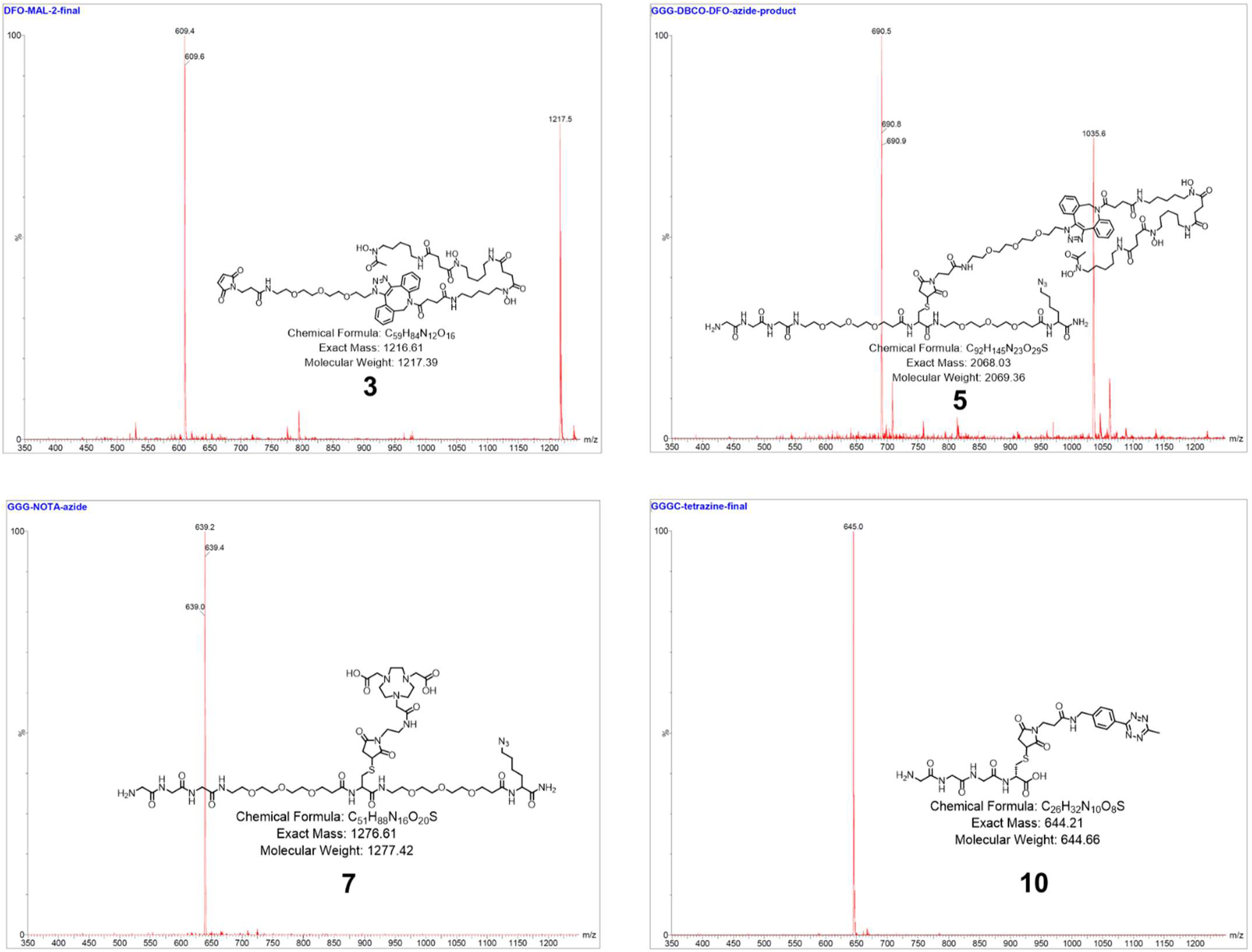
Mass spectra of the labeling reagents shown in Supplemental Figure 12.

**Supplemental Figure 14.**
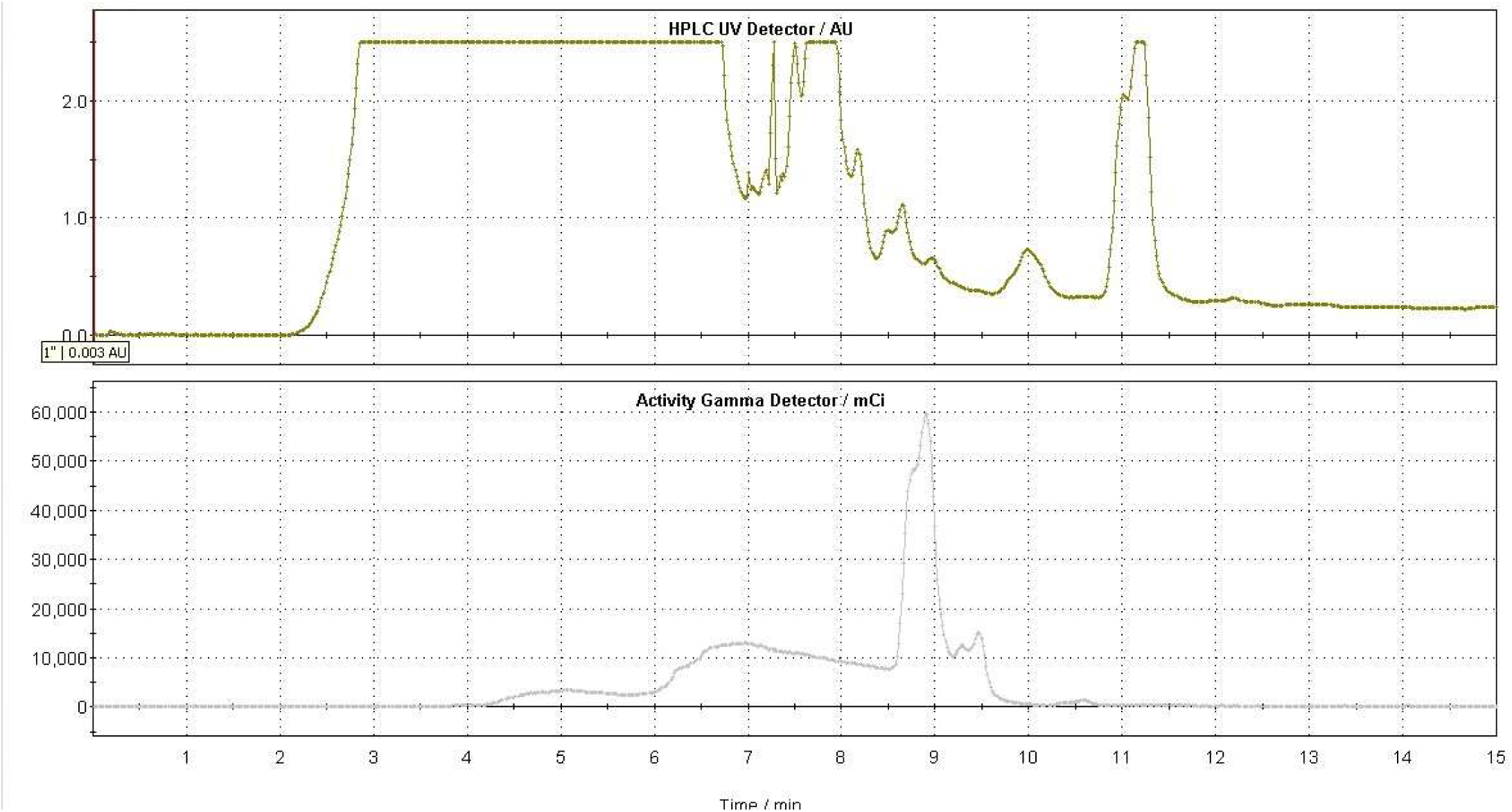
Semi-preparative HPLC gamma chromatogram of 18F-TCO (collected from 8.5 to 9.2 min).

**Supplemental Figure 15.**
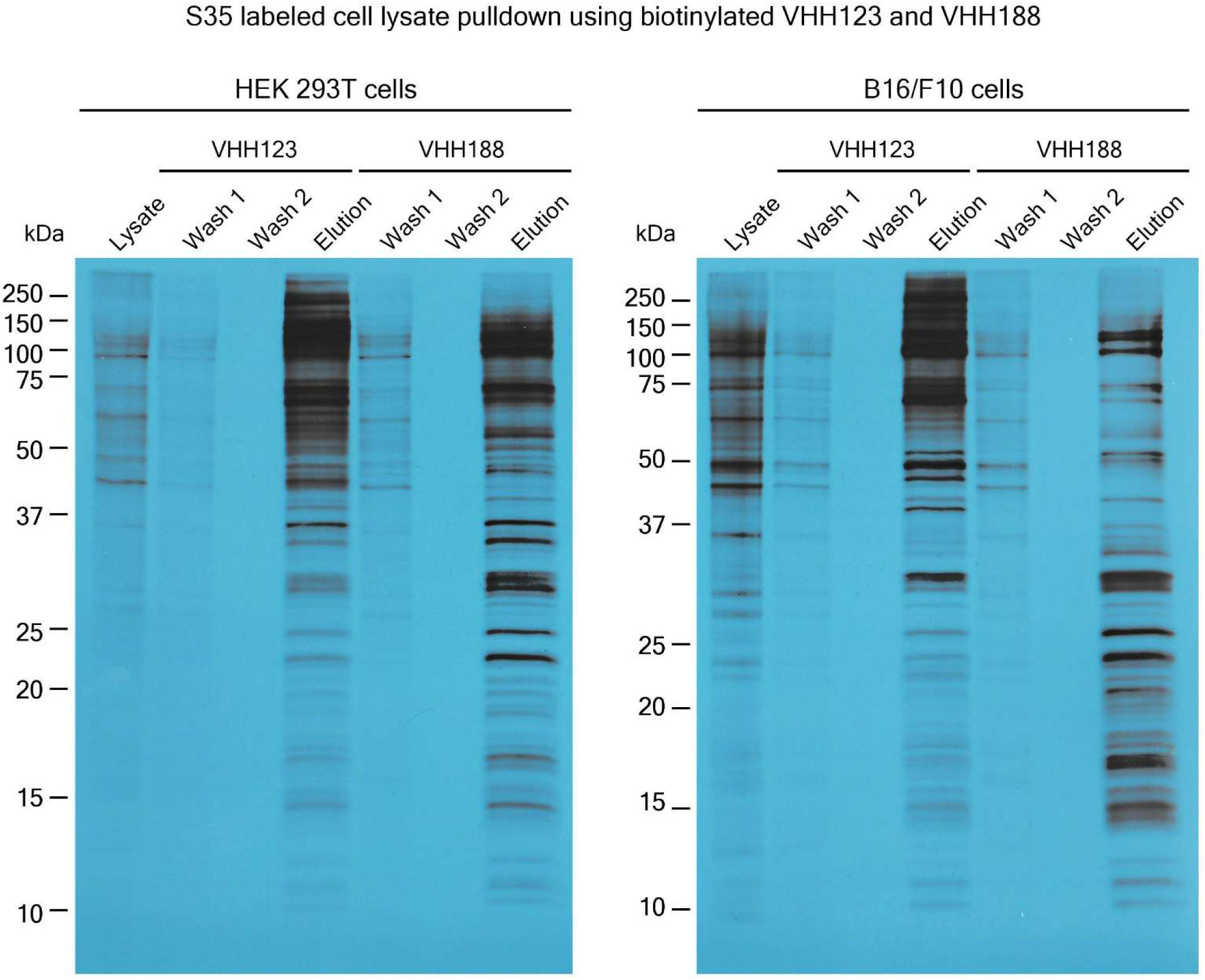
Autoradiograph of ^35^S labelled lysates of HEK 293T cells and B16.F10 cells that were used in pull-down experiments for figure 1B. Cell-lines were incubated with ^35^S-labelled Met before incubation (see methods) with nanobody coated paramagnetic beads overnight (VHH123 or VHH188 as shown above corresponding lanes). Beads were then washed 2 times then boiled in Laemli buffer before loading on SDS-PAGE. SDS-PAGE was run with input cell lysate (lysate), washes 1 and 2 and eluates for each condition. Gel was then prepared for autoradiography (see supplementary methods) before being placed on top of an X-ray film. Gel and film were stored together at −80°C for 24h before developing.

## Supplementary Methods

### Labeling reagent synthesis

The synthetic routes of labeling reagents are shown in supplemental figure 12, and their corresponding mass spectra are shown in supplemental figure 13.

### Materials

Deferoxamine-DBCO (**1**) was purchased from Macrocyclics (cat. no. B-773). Azido-PEG3-Maleimide (**2**) was purchased from Vector Laboratories (cat. no. CCT-AZ107-100). Maleimido-mono-amide-NOTA (**6**) was obtained from Macrocyclics (cat. no. B-622). Methyltetrazine-Maleimide (**8**) was purchased from Conju-Probe (cat. no. CP-6608-25mg). All the other chemical reagents and solvents were purchased from Sigma-Aldrich. The synthesis of the peptide GGG-PEG_3_-Cys-PEG_3_-Lys(azide) (**4**) and Gly-Gly-Gly-Cys (**9**) were synthesized as described ^24, 53^. The molecular mass of the labeling reagents was determined using an LC-MS system (Waters QDa_Arc). All labeling reagents were purified by HPLC (Shimadzu) equipped with an XBridge BEH C18 OBD Prep Column (130Å, 5 µm, 19 mm × 150 mm).

### Synthesis of GGG-DFO-N_3_ (5)

To a solution of deferoxamine-DBCO (**1**) (25 mg, 0.029 mmol) in anhydrous DMSO (0.5 mL), azido-PEG₃-maleimide (**2**) (13 mg, 0.035 mmol) dissolved in anhydrous DMSO (0.5 mL) was added and stirred for 1 hour at room temperature. The reaction mixture was directly injected into HPLC (0–100% acetonitrile in H₂O containing 0.1% TFA), yielding intermediate (**3**) as a white powder (29 mg, 82%). LC-MS: [M + H]⁺ = 1217.5.

A mixture of intermediate (**3**) (29 mg, 0.024 mmol) and peptide GGG-PEG₃-Cys-PEG₃-Lys(azide) (**4**) (41 mg, 0.048 mmol), dissolved in DMSO (1.5 mL), was stirred overnight at room temperature. The reaction mixture was then directly injected into HPLC (0–100% acetonitrile in H₂O containing 0.1% TFA), yielding GGG-DFO-N₃ (**5**) as a colorless oil (32 mg, 64%). LC-MS: [M + 2H]²⁺ = 1035.6.

### Synthesis of GGG-NOTA-N_3_

A mixture of GGG-PEG₃-Cys-PEG₃-Lys(azide) (**4**) (78 mg, 0.092 mmol) and maleimido-mono-amide-NOTA (**6**) (25 mg, 0.046 mmol), dissolved in DMSO (2 mL), was stirred overnight at room temperature. The reaction mixture was then directly injected into HPLC (0–100% acetonitrile in H₂O containing 0.1% TFA), yielding GGG-NOTA-N₃ (**7**) as a colorless oil (41 mg, 70%). LC-MS: [M + 2H]²⁺ = 639.2.

### Synthesis of GGG-tetrazine

A mixture of methyltetrazine-maleimide (**8**) (25 mg, 0.071 mmol) and Gly-Gly-Gly-Cys (**9**) (62 mg, 0.213 mmol), dissolved in DMSO (1.5 mL), was stirred overnight at room temperature. The reaction mixture was then directly injected into HPLC (0–100% acetonitrile in H₂O containing 0.1% TFA), yielding GGG-tetrazine (**10**) as a red powder (23 mg, 50%). LC-MS: [M + H]⁺ = 645.0.

### Radiosynthetic Method for generating ^18^F-TCO

Approximately 3000 mCi of [18F]fluoride was produced on a GE PETtrace 800 cyclotron and delivered to the 18F target delivery vial of a GE FX2 N radiosynthesis module. The irradiated target water was passed through a pre-conditioned Sep-PAK Light QMA Cartridge to trap [18F]fluoride, followed by elution from the cartridge into reactor 1 using 2.5 mg K2CO2 and 18 mg Cryptand 222 in 0.1 mL of H2O (HyClone) and 0.9 mL acetonitrile (HPLC). The contents of reactor 1 were dried via azeotropic distillation using a combination of heating, helium flow, and vacuum. An additional 1 mL of acetonitrile (anhydrous) was introduced to reactor 1 and the azeotropic drying process was repeated. The reactor temperature was then reduced to 40 °C, followed by introduction of TCO-nosylate (4 mg, Peptech) in 0.9 mL acetonitrile (anhydrous). The reactor was then sealed and heated to 75 °C and allowed to react with stirring for 10 minutes. Temperature of the reactor was cooled down to 40 °C and the reaction was quenched with the addition of 5 mL of sodium ascorbate solution (6.5 mg/mL). The reaction mixture was filtered through an Alumina N Sep-Pak cartridge (pre-conditioned with 15 mL of H2O) and loaded onto a 5 mL HPLC Loop which had been previously filled with semi-preparative HPLC mobile phase (80% acetonitrile, HPLC, in water, HPLC) to minimize injection of air onto the HPLC column. The crude mixture was then injected on a C18 semi-preparative HPLC column (flow rate = 3.5 mL/min). The 18F-TCO product peak (∼8.5-9.2 minutes – supplemental figure 14) was collected into a round bottom flask containing 20 mL sterile water. The contents of the round bottom flask were then passed through a C18 Plus Light Sep-PAK cartridge, where 18F-TCO was trapped. The cartridge was then washed with 5 mL water (USP, sterile for irrigation), followed by elution of 18F-TCO to the FX2 N product vial using 0.5 mL ethanol. 291 mCi of 18F-TCO was obtained in 0.42 g of EtOH for generation of VHH123-^18^F; 237 mCi of 18-TCO was obtained in 0.34 g of EtOH for generation of VHH188-^18^F. See methods for click-chemistry of 18F-TCO onto VHH-tetrazine conjugates.

### Autoradiography

To perform autoradiography, after running, the SDS-PAGE gel was soaked in DMSO for 30 min, twice, to remove all aqueous solutions. Gel was then incubated in DMSO with 20% w/v 2,5 diphenyloxazole (PPO, insoluble in water) for 1h to allow PPO incorporation. Gel was then washed multiple times in water and dried using a gel dryer. Then, the gel was placed against an X-ray film in a cassette and stored at −80°C for 24 hours before developing the film.

