## Supplemental Figures 3-7 for "A pair of congenic mice for imaging of transplants by positron emission tomography using anti-transferrin receptor nanobodies"

### Slide 1
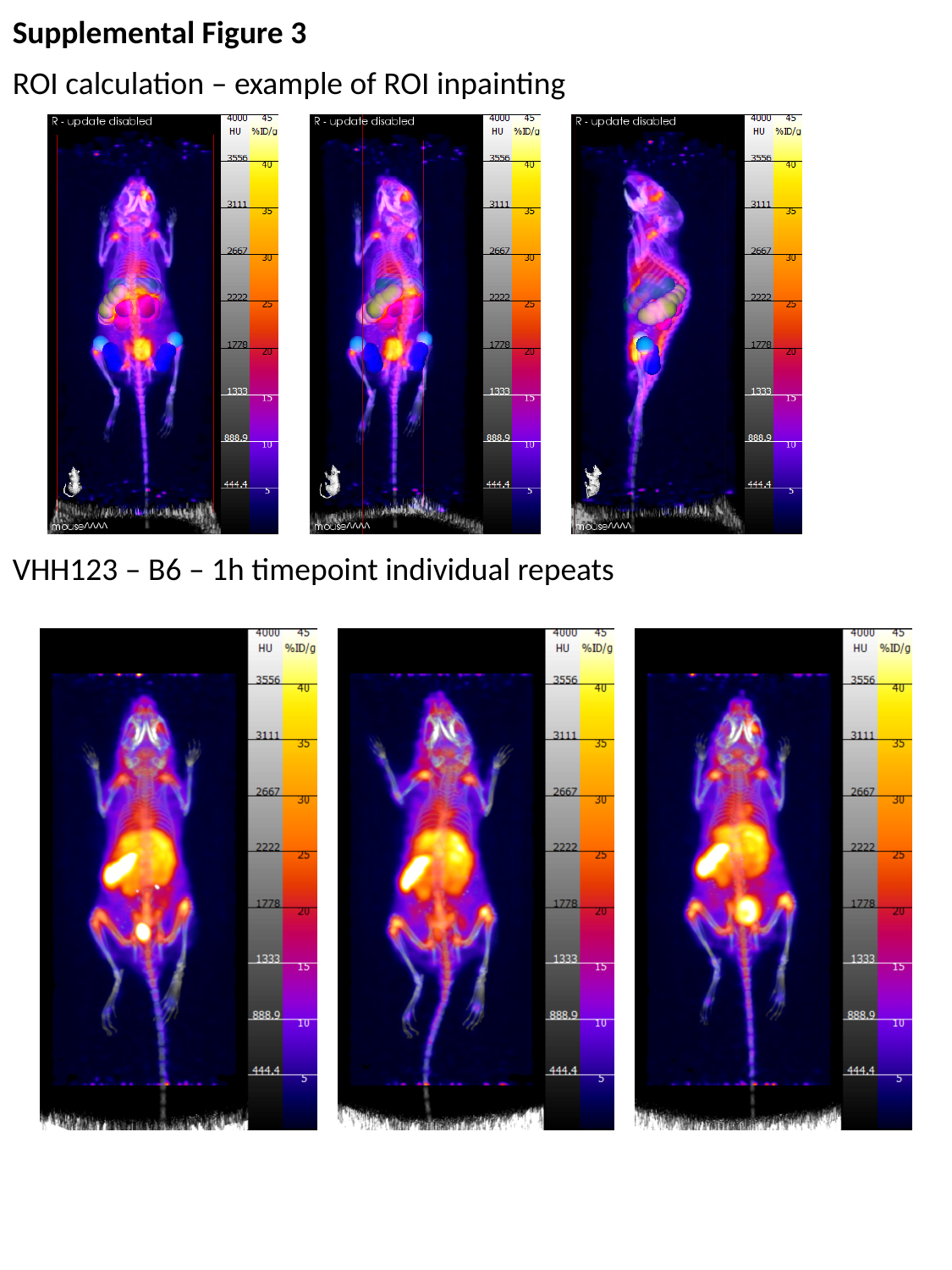

Supplemental Figure 3
ROI calculation – example of ROI inpainting
VHH123 – B6 – 1h timepoint individual repeats

### Slide 2
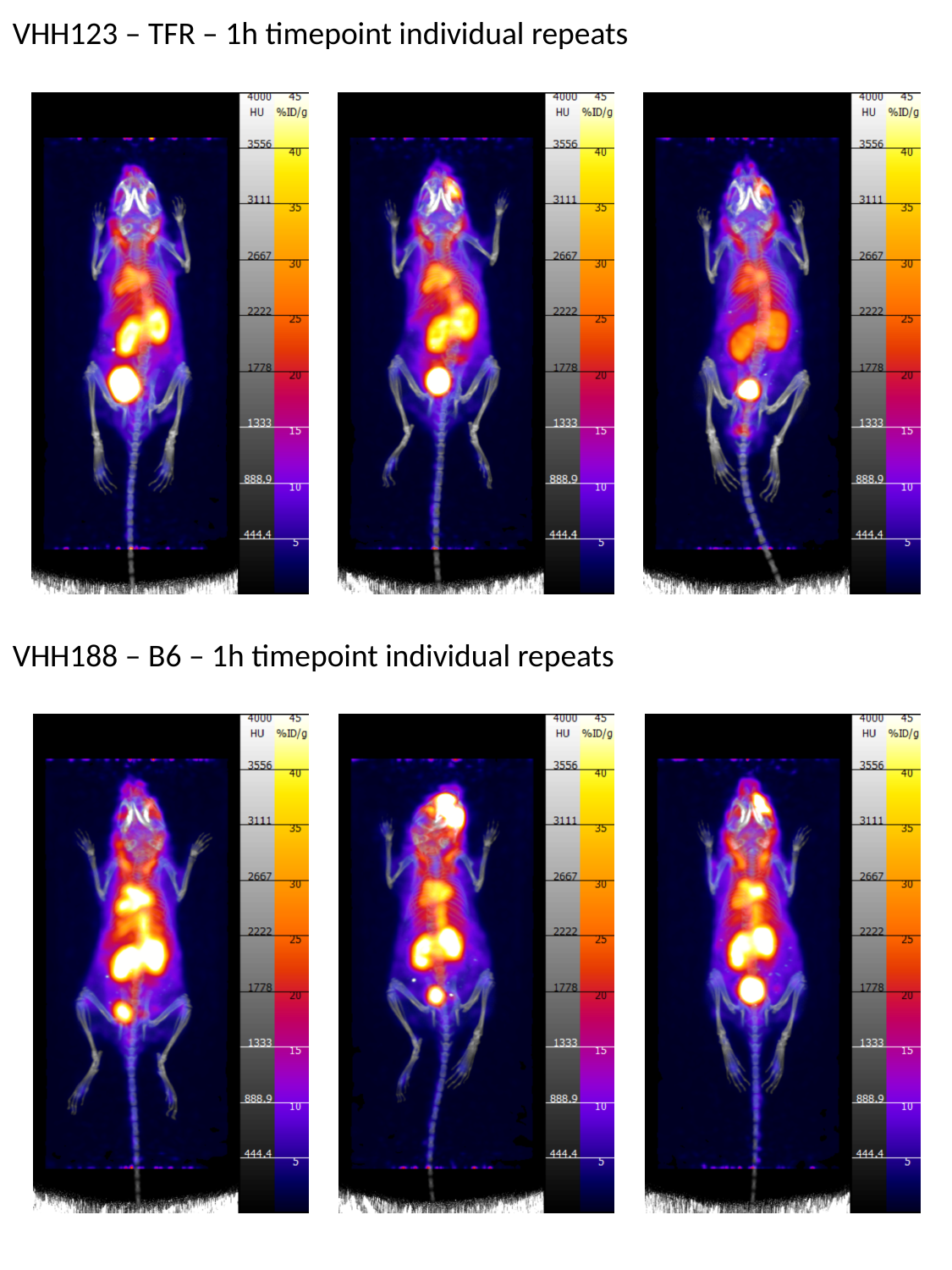

VHH123 – TFR – 1h timepoint individual repeats
VHH188 – B6 – 1h timepoint individual repeats

### Slide 3
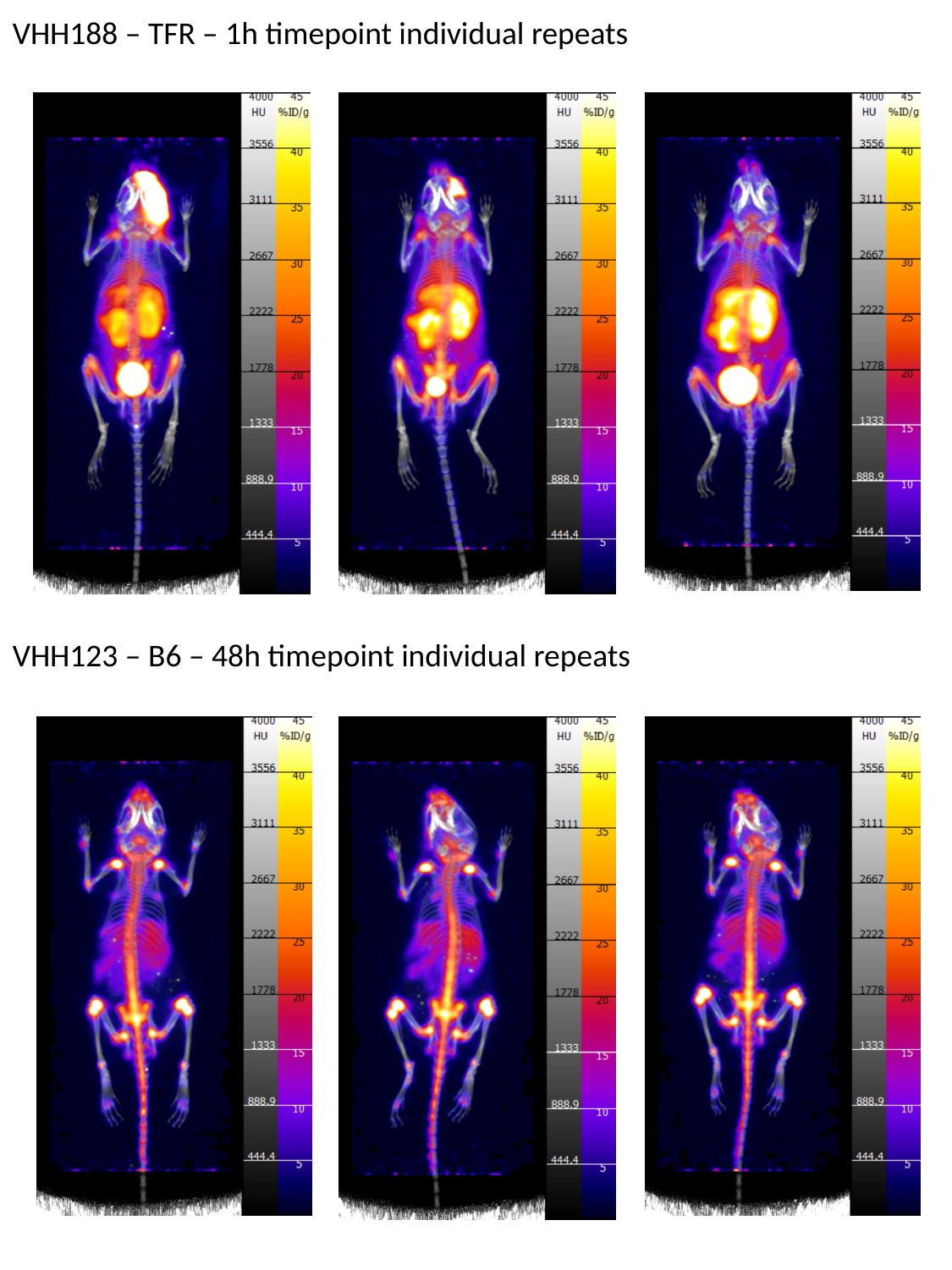

VHH188 – TFR – 1h timepoint individual repeats
VHH123 – B6 – 48h timepoint individual repeats

### Slide 4
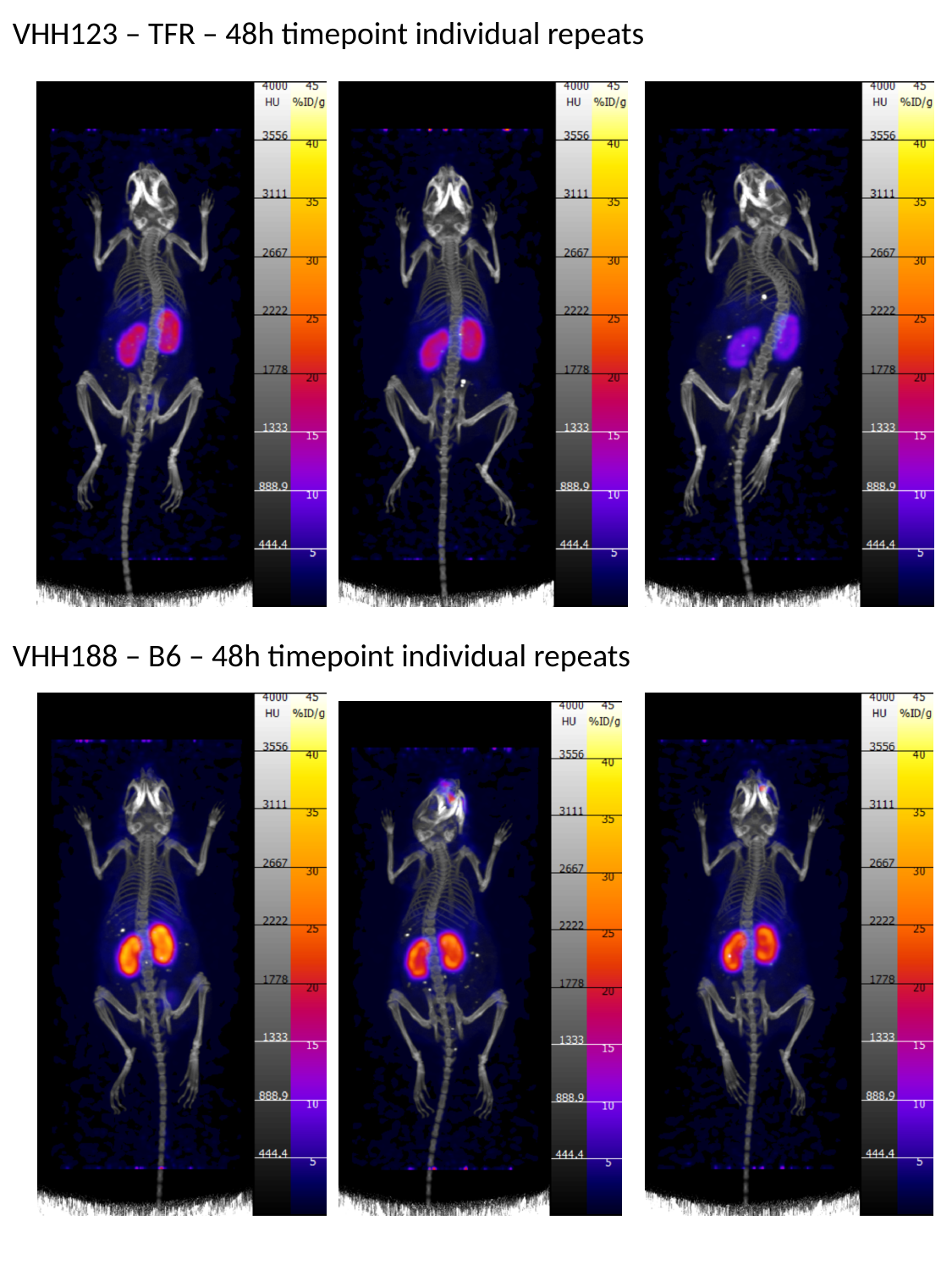

VHH123 – TFR – 48h timepoint individual repeats
VHH188 – B6 – 48h timepoint individual repeats

### Slide 5
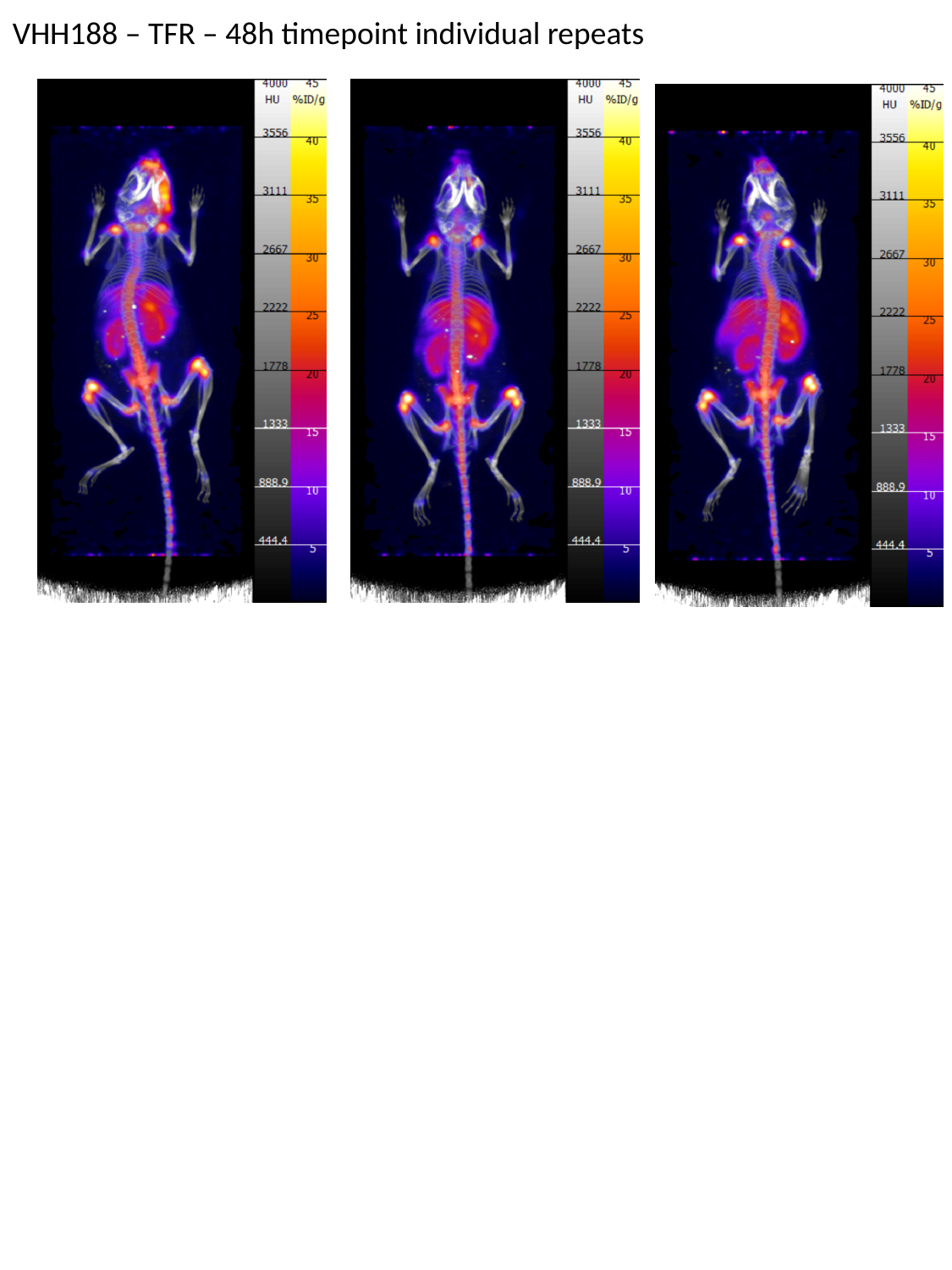

VHH188 – TFR – 48h timepoint individual repeats

### Slide 6
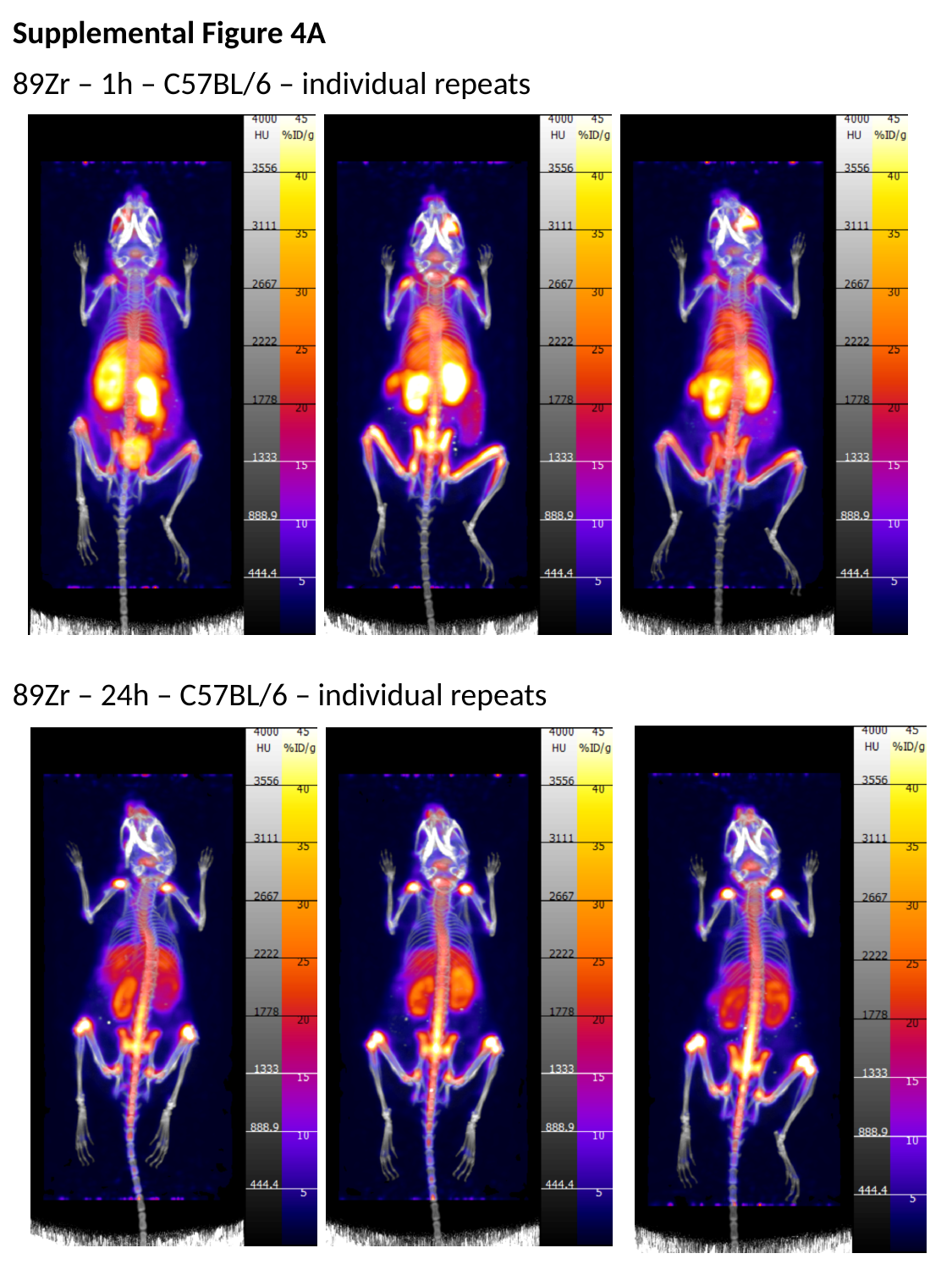

Supplemental Figure 4A
89Zr – 1h – C57BL/6 – individual repeats
89Zr – 24h – C57BL/6 – individual repeats

### Slide 7
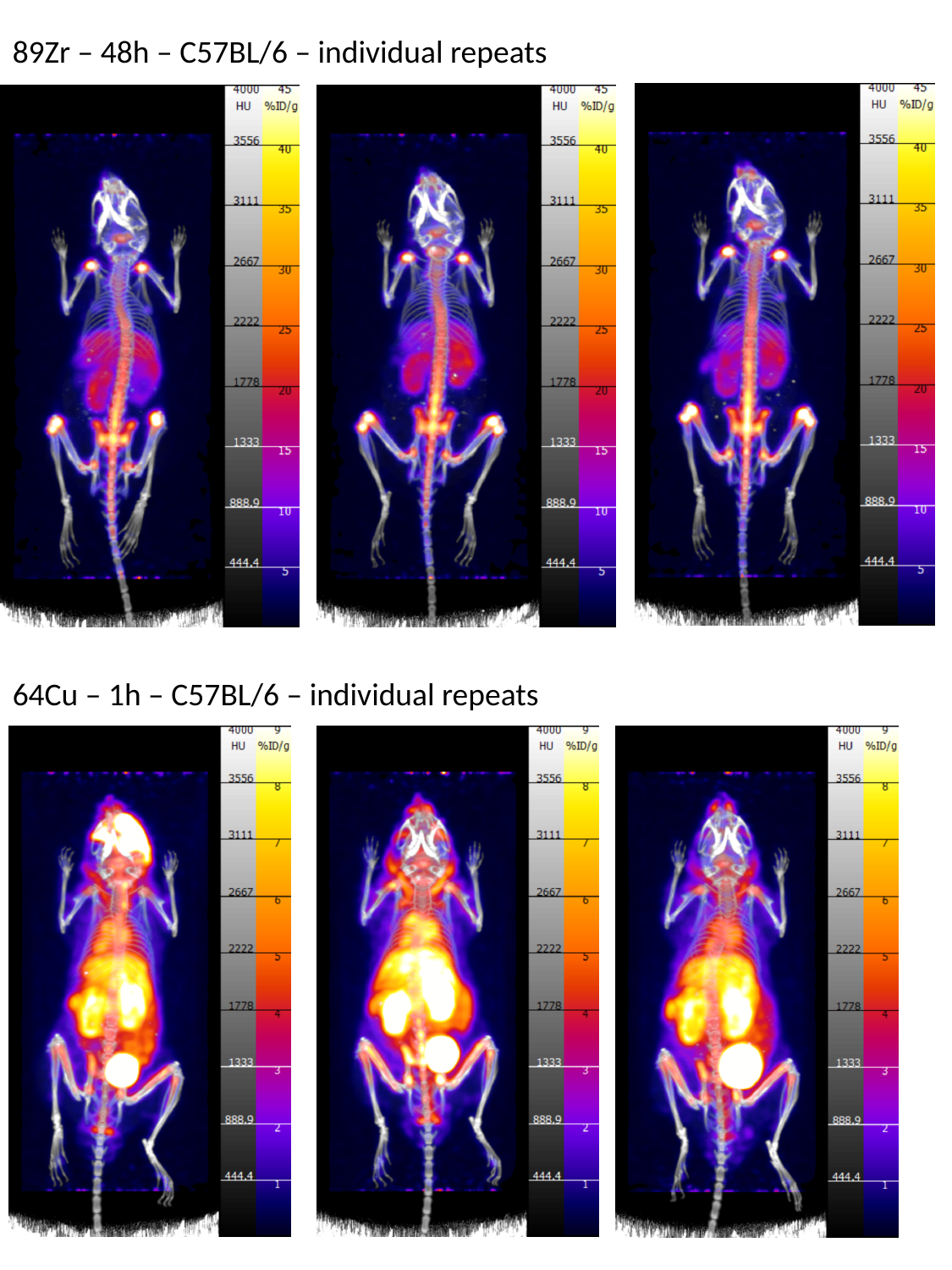

89Zr – 48h – C57BL/6 – individual repeats
64Cu – 1h – C57BL/6 – individual repeats

### Slide 8
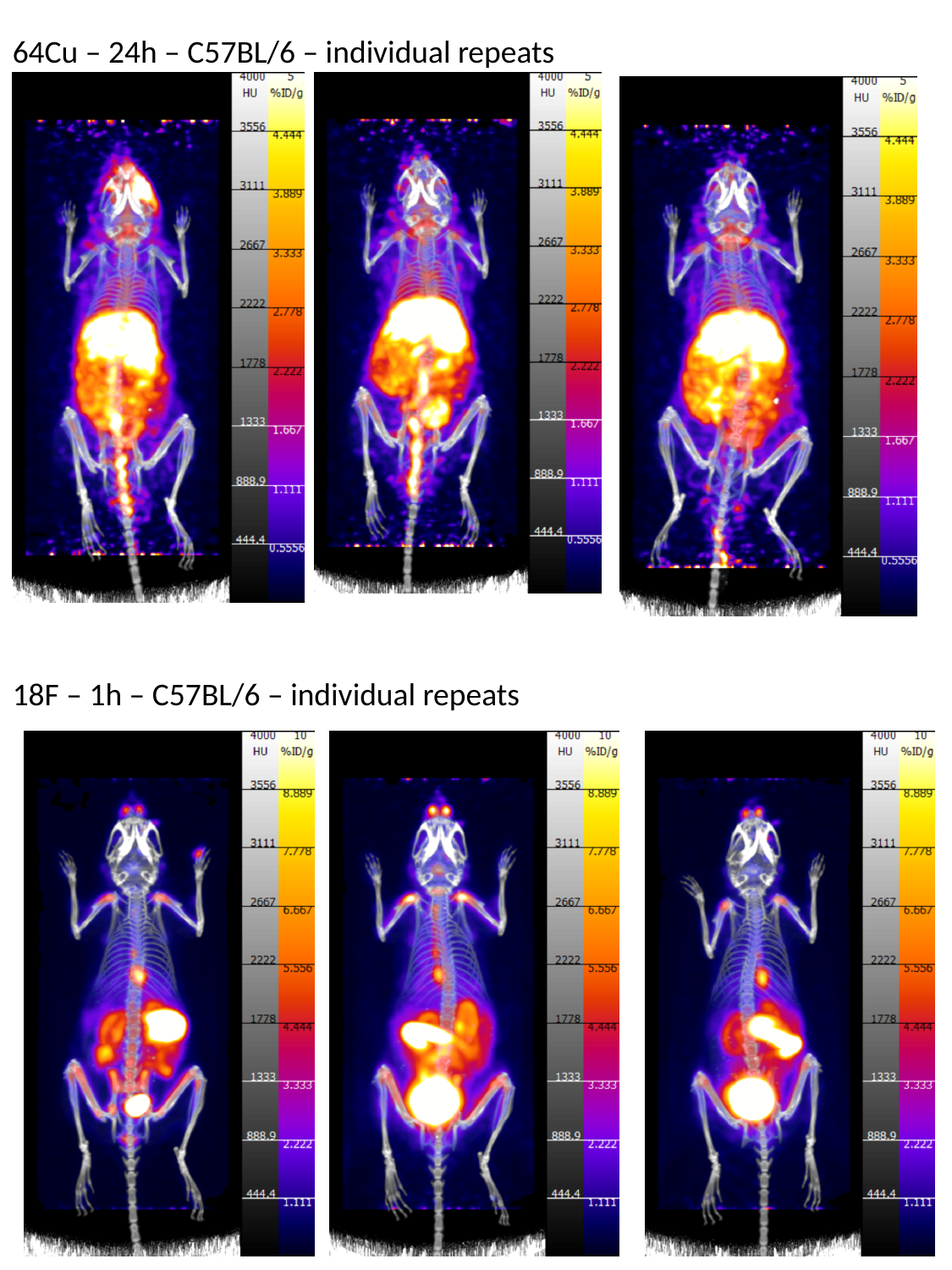

64Cu – 24h – C57BL/6 – individual repeats
18F – 1h – C57BL/6 – individual repeats

### Slide 9
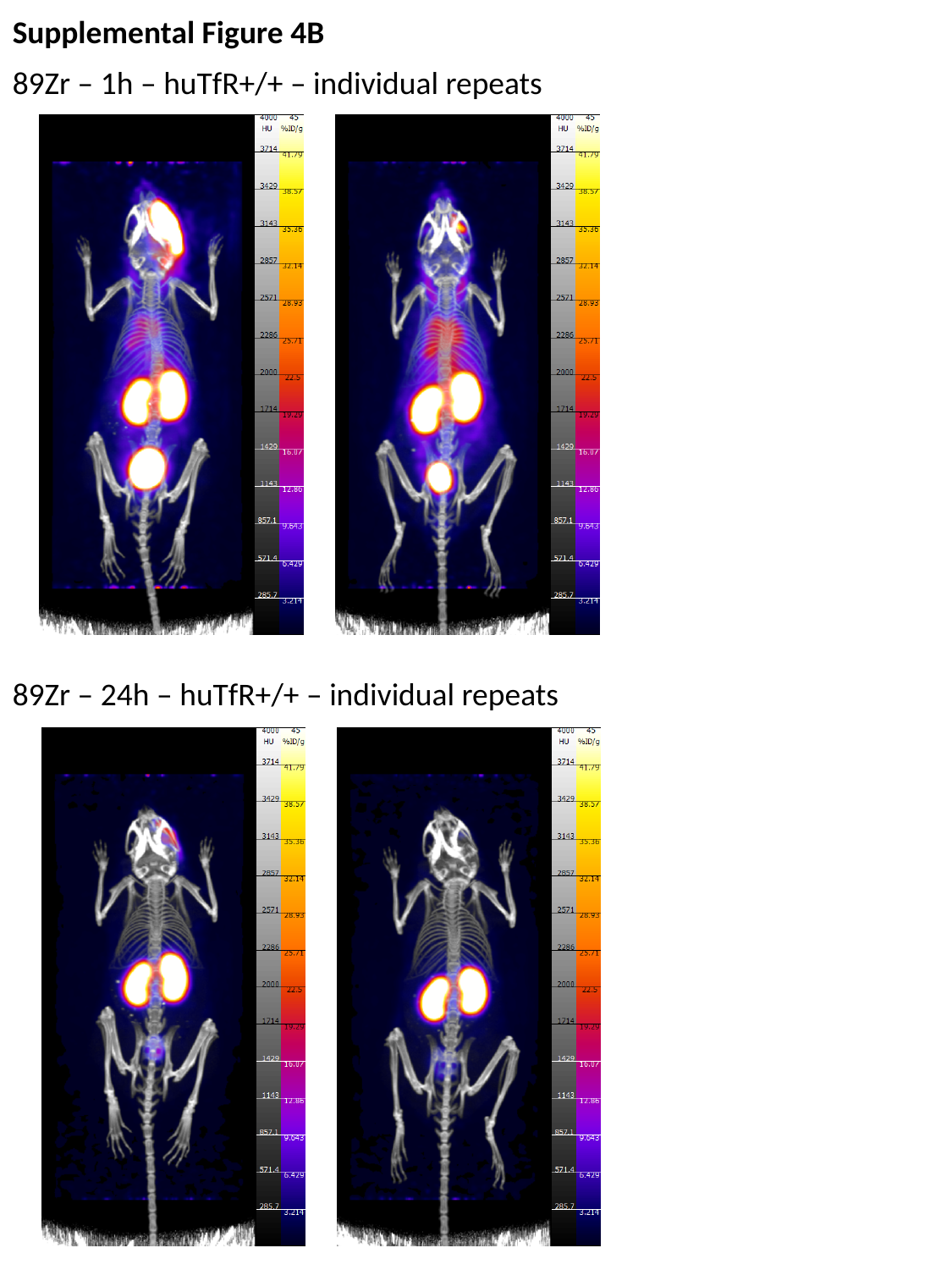

Supplemental Figure 4B
89Zr – 1h – huTfR+/+ – individual repeats
89Zr – 24h – huTfR+/+ – individual repeats

### Slide 10
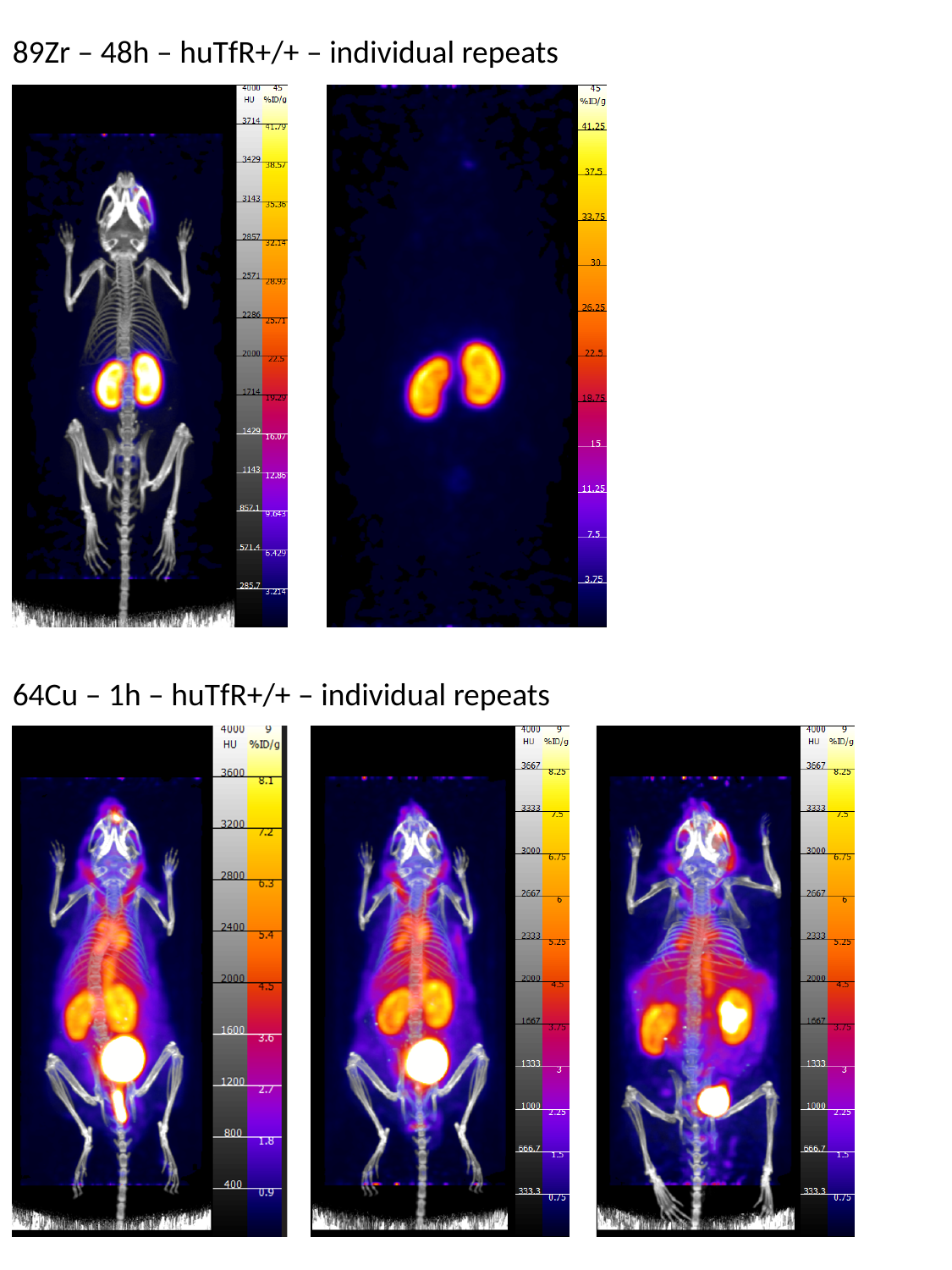

89Zr – 48h – huTfR+/+ – individual repeats
64Cu – 1h – huTfR+/+ – individual repeats

### Slide 11
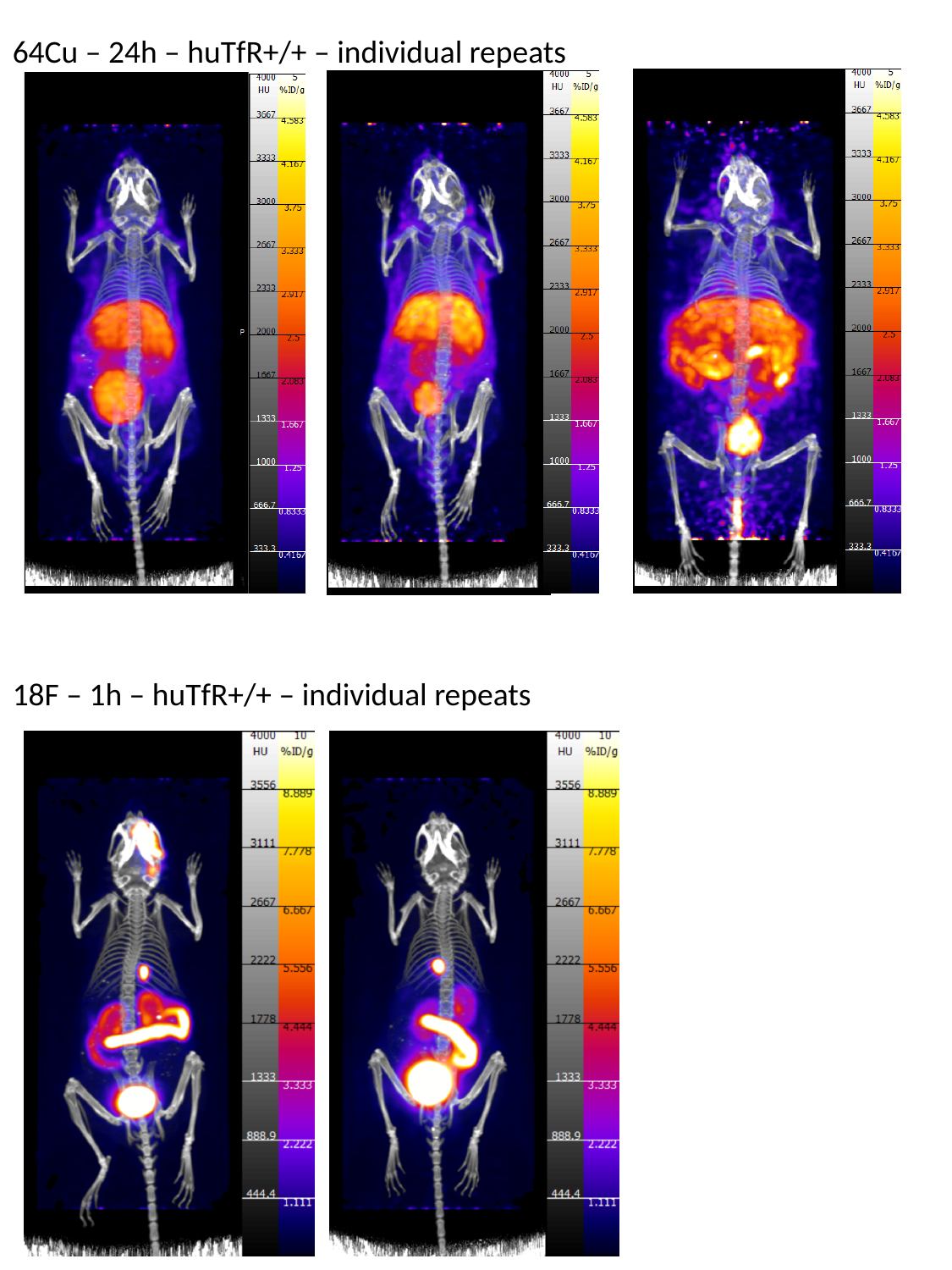

64Cu – 24h – huTfR+/+ – individual repeats
18F – 1h – huTfR+/+ – individual repeats

### Slide 12
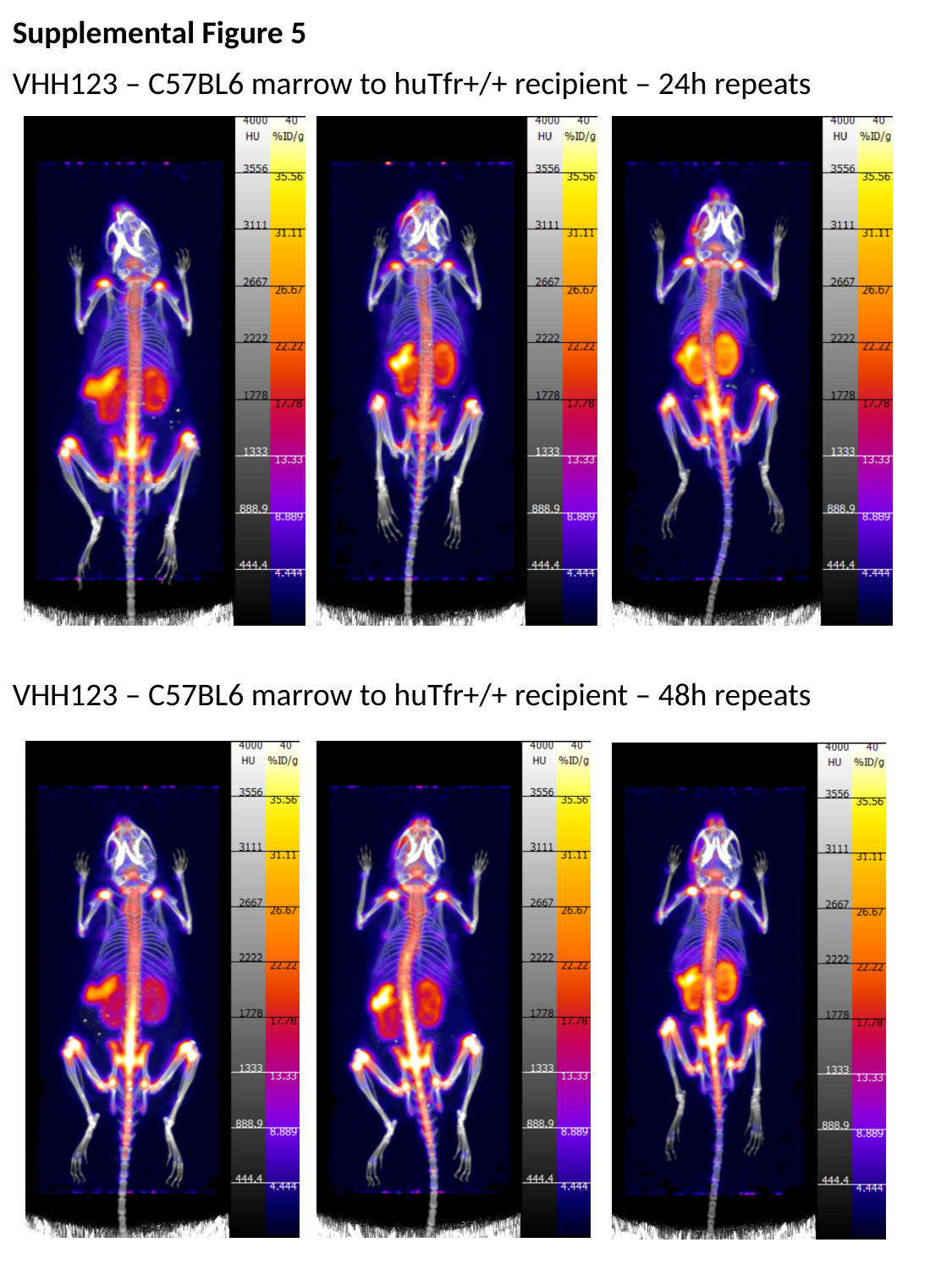

Supplemental Figure 5
VHH123 – C57BL6 marrow to huTfr+/+ recipient – 24h repeats
VHH123 – C57BL6 marrow to huTfr+/+ recipient – 48h repeats

### Slide 13
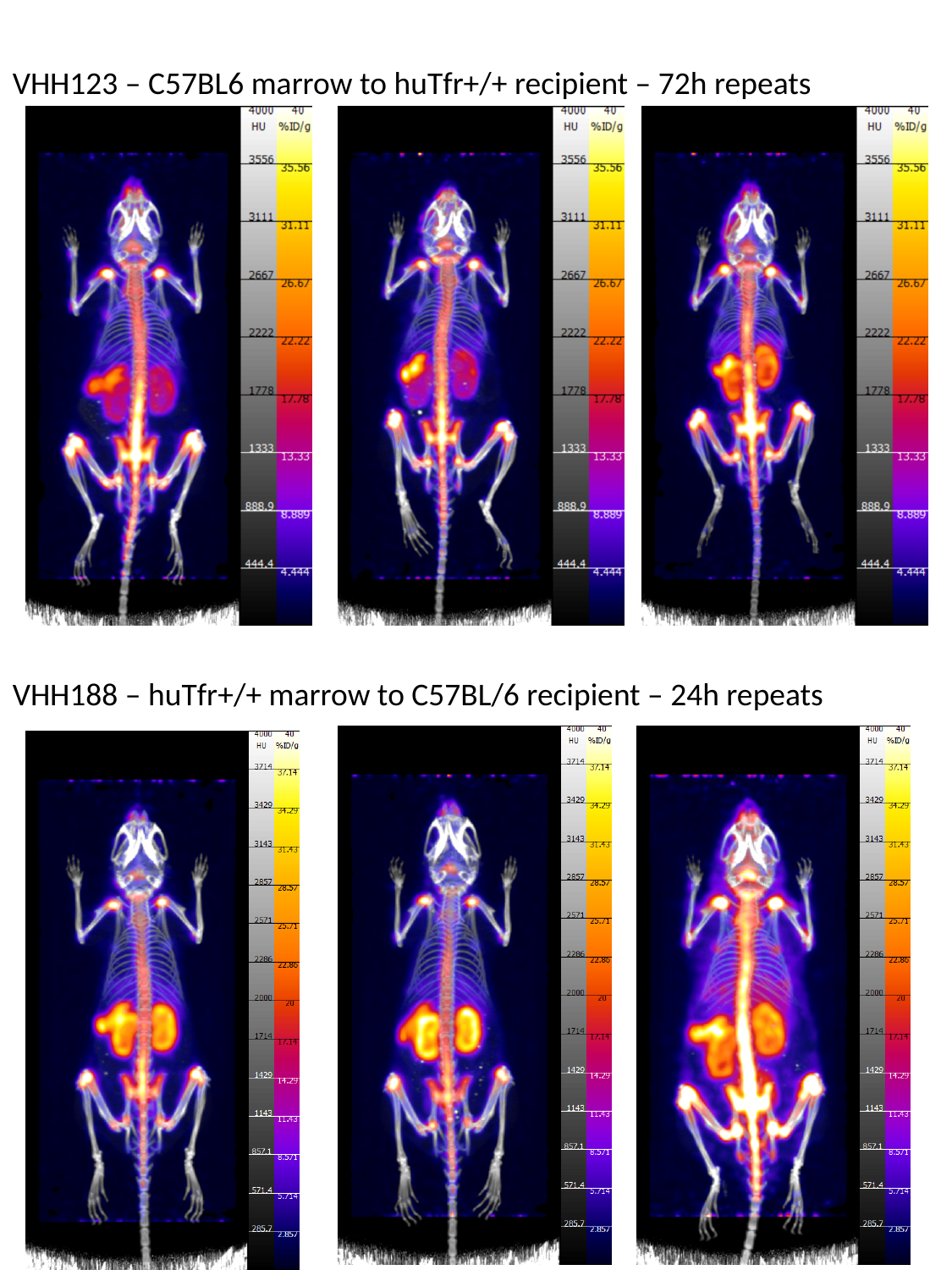

VHH123 – C57BL6 marrow to huTfr+/+ recipient – 72h repeats
VHH188 – huTfr+/+ marrow to C57BL/6 recipient – 24h repeats

### Slide 14
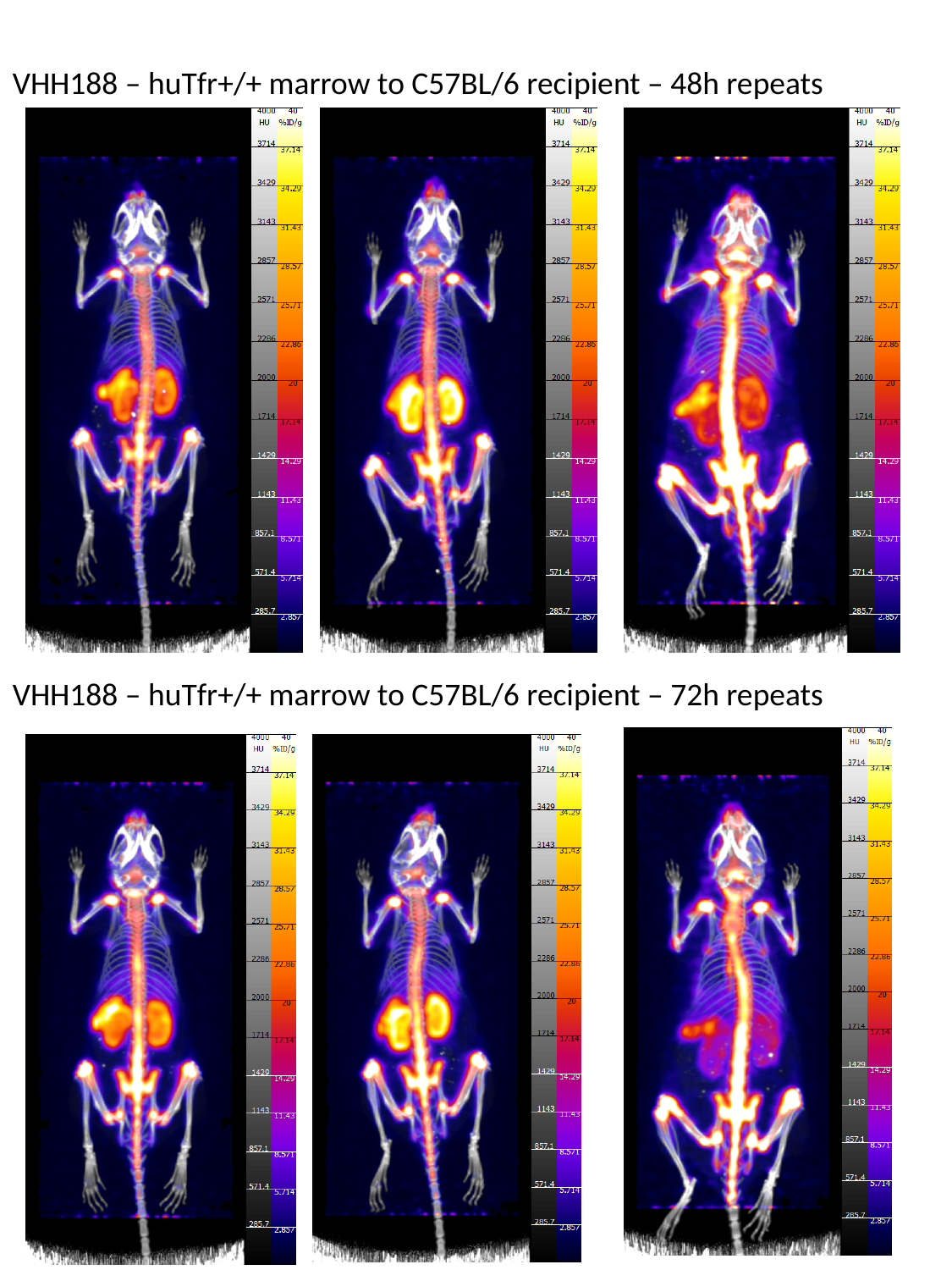

VHH188 – huTfr+/+ marrow to C57BL/6 recipient – 48h repeats
VHH188 – huTfr+/+ marrow to C57BL/6 recipient – 72h repeats

### Slide 15
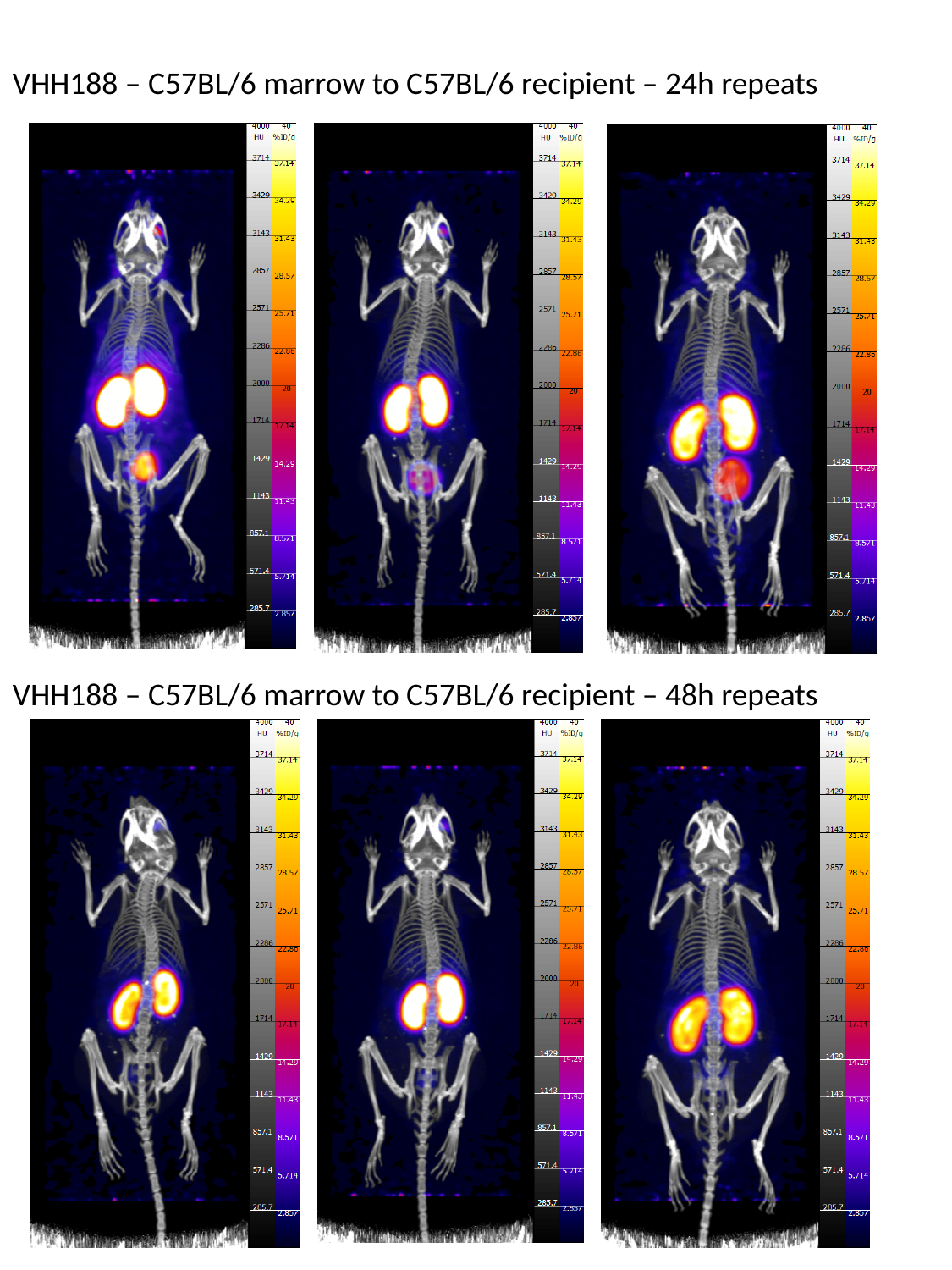

VHH188 – C57BL/6 marrow to C57BL/6 recipient – 24h repeats
VHH188 – C57BL/6 marrow to C57BL/6 recipient – 48h repeats

### Slide 16
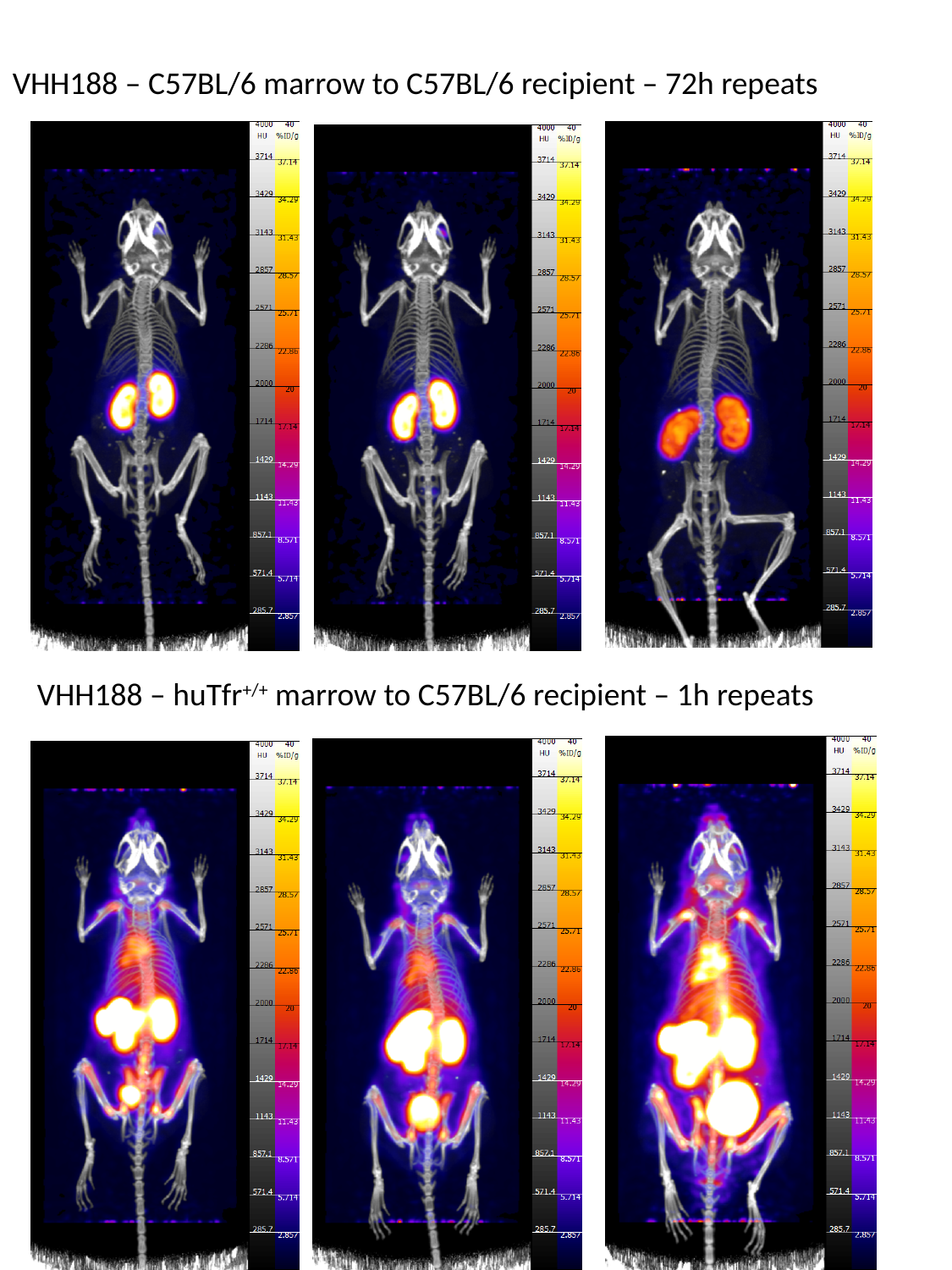

VHH188 – C57BL/6 marrow to C57BL/6 recipient – 72h repeats
VHH188 – huTfr^+/+^ marrow to C57BL/6 recipient – 1h repeats

### Slide 17
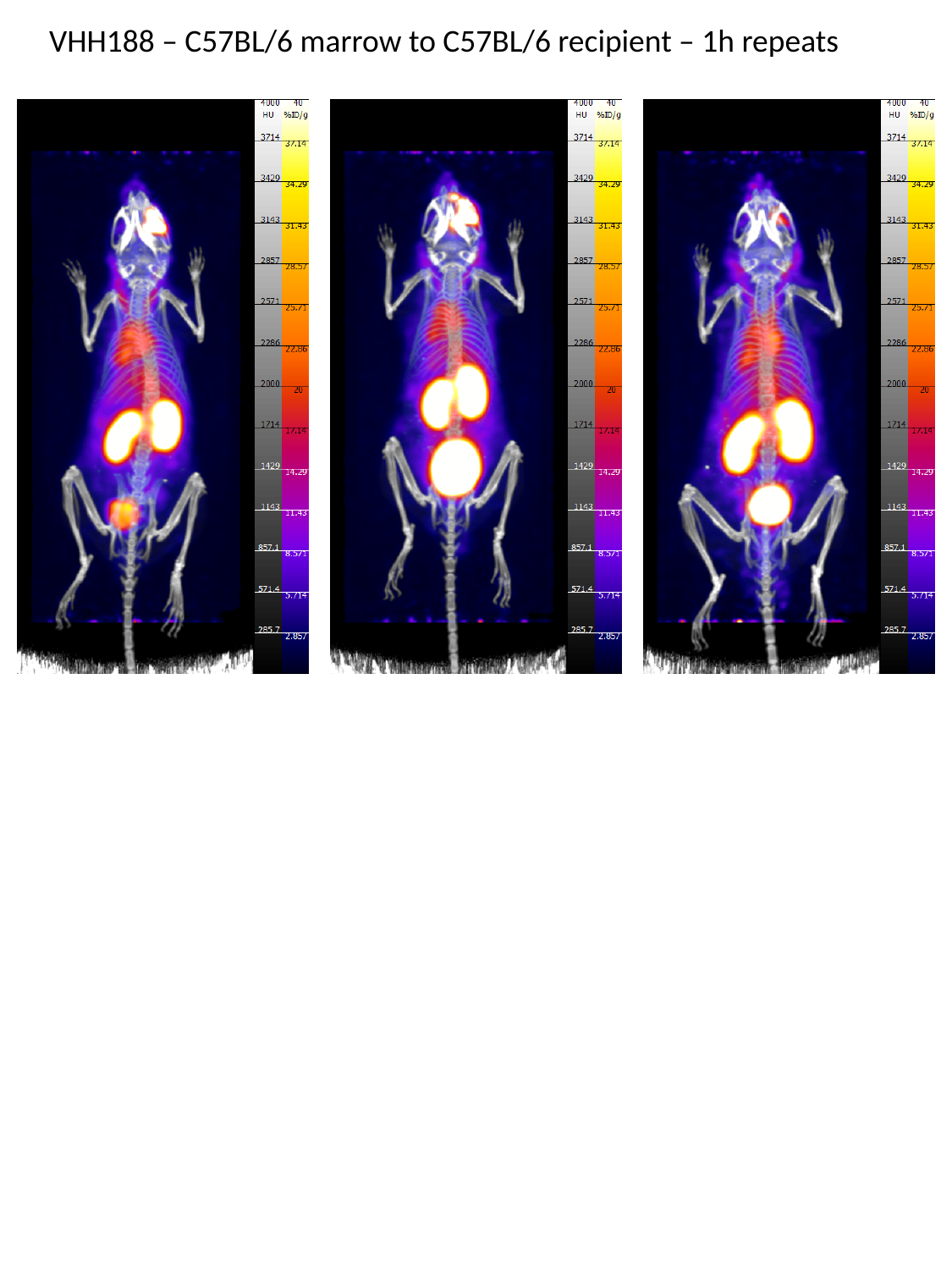

VHH188 – C57BL/6 marrow to C57BL/6 recipient – 1h repeats

### Slide 18
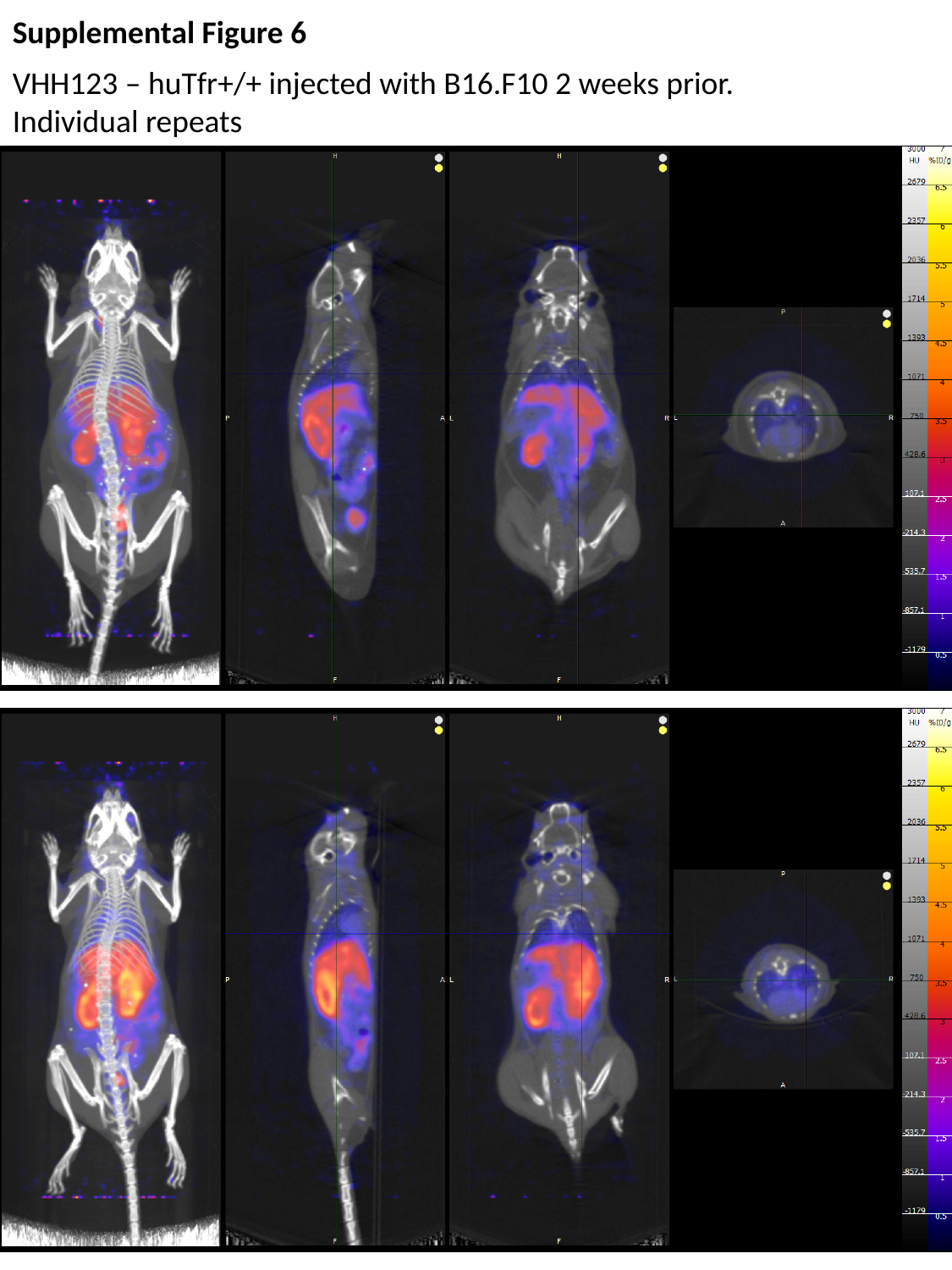

Supplemental Figure 6
VHH123 – huTfr+/+ injected with B16.F10 2 weeks prior.
Individual repeats

### Slide 19
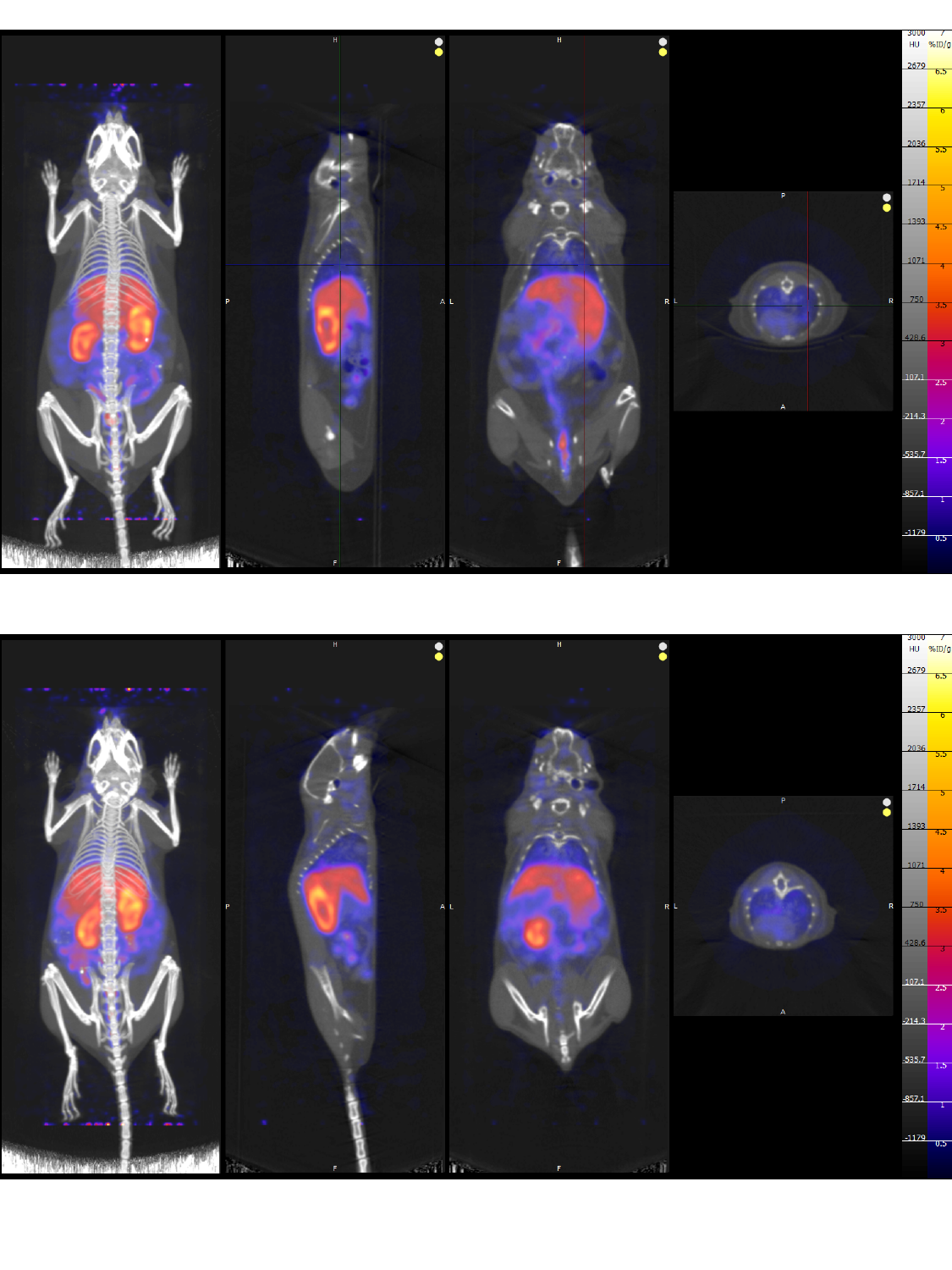

### Slide 20
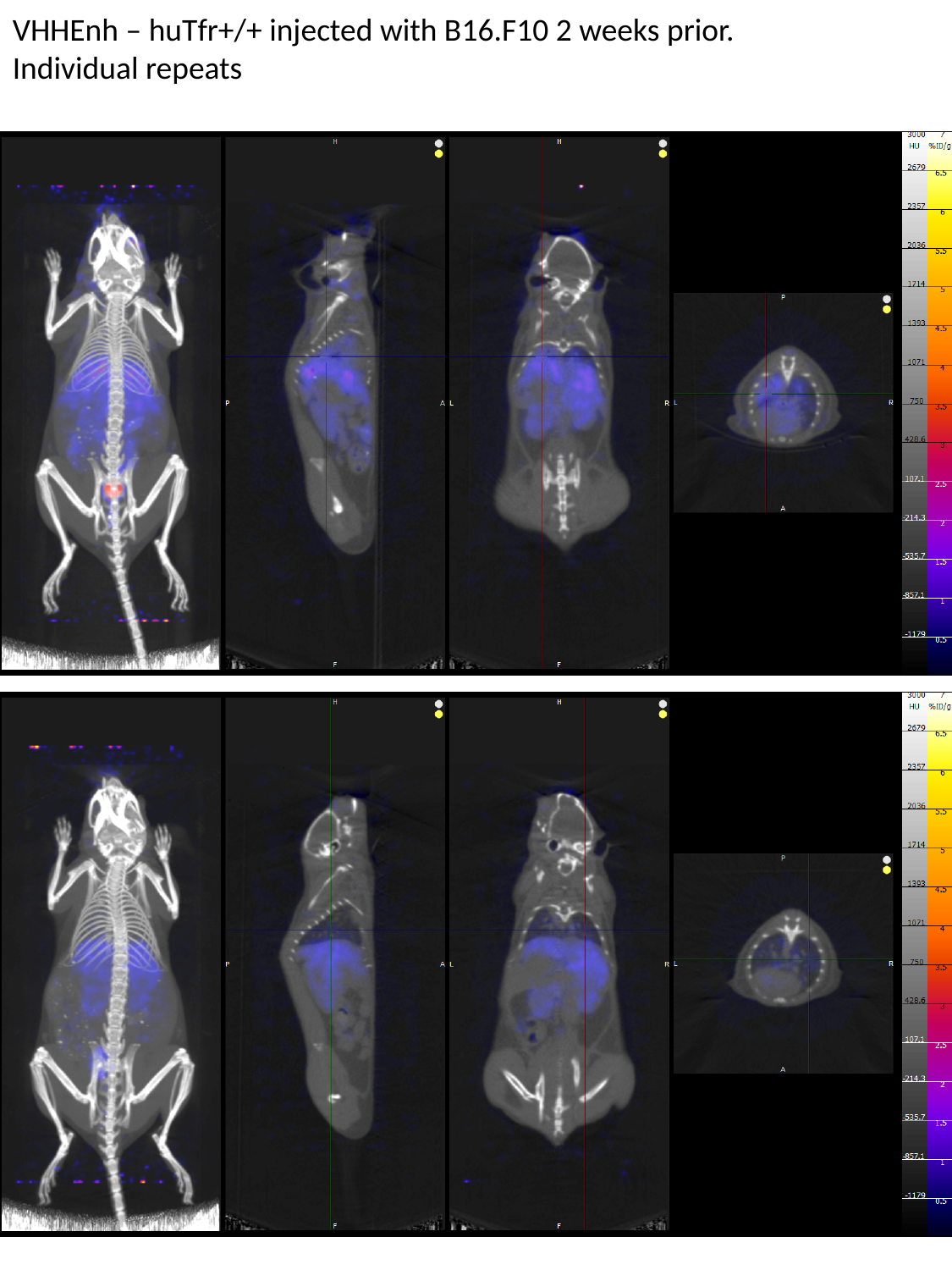

VHHEnh – huTfr+/+ injected with B16.F10 2 weeks prior.
Individual repeats

### Slide 21
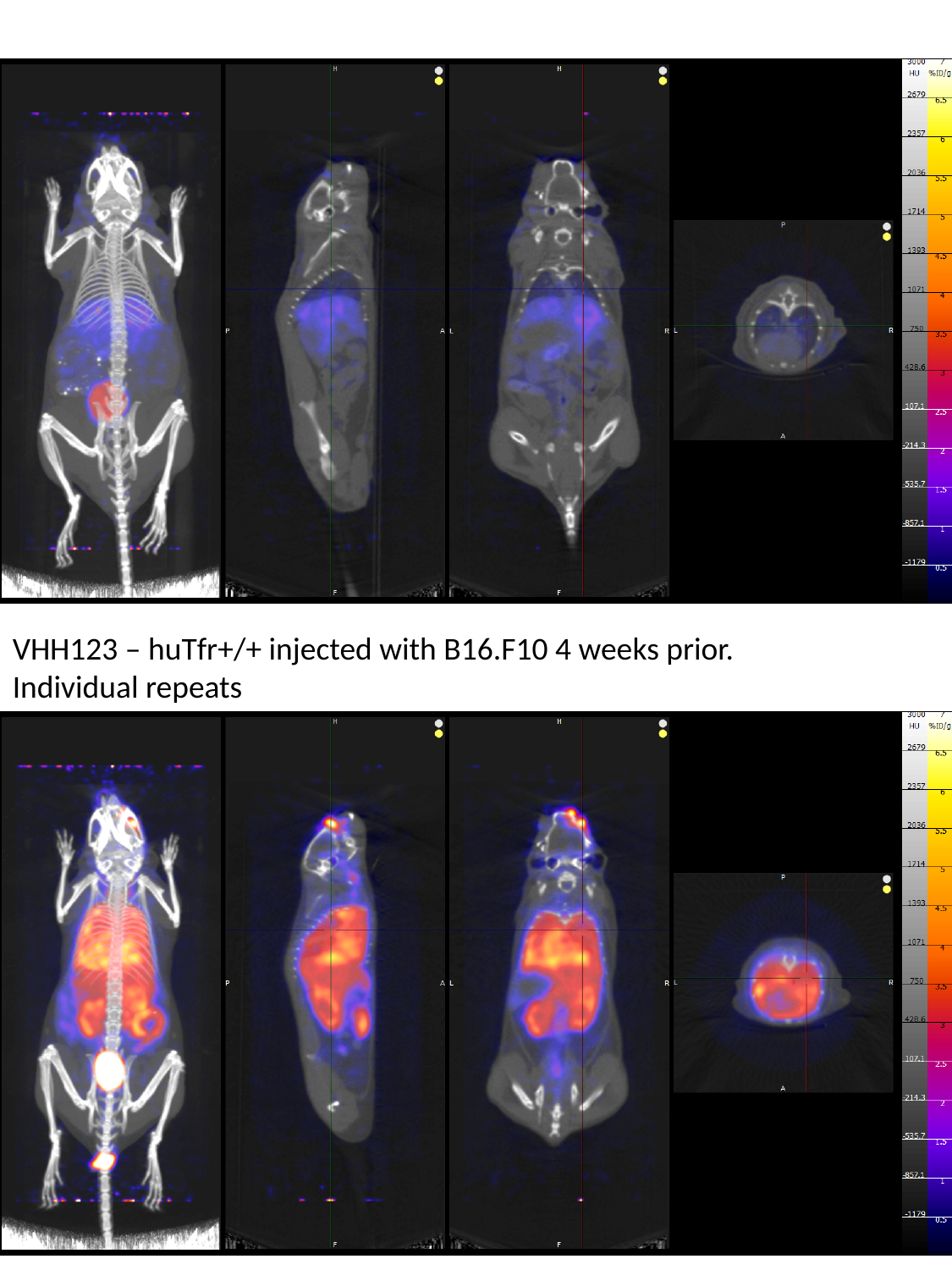

VHH123 – huTfr+/+ injected with B16.F10 4 weeks prior.
Individual repeats

### Slide 22
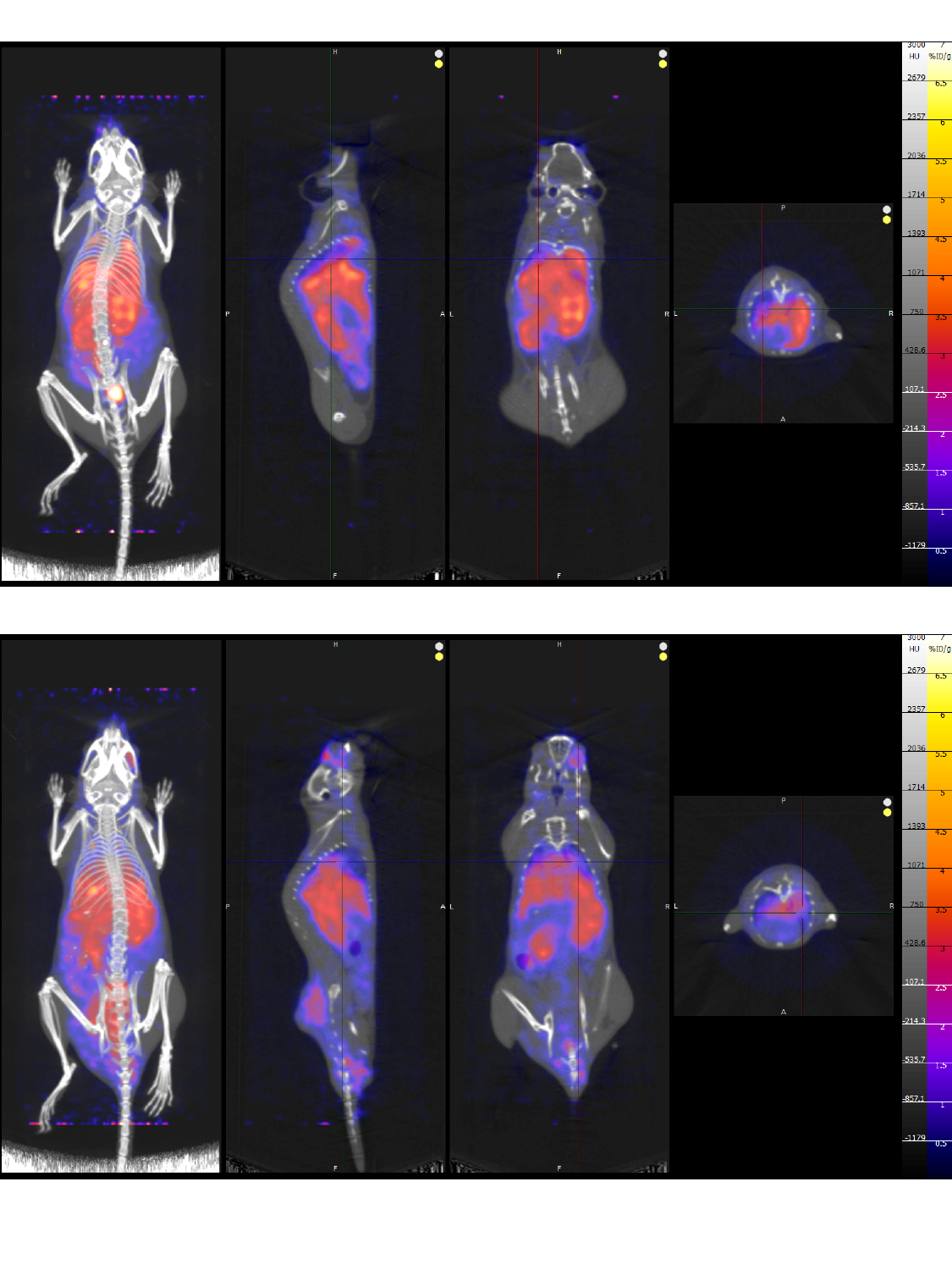

### Slide 23
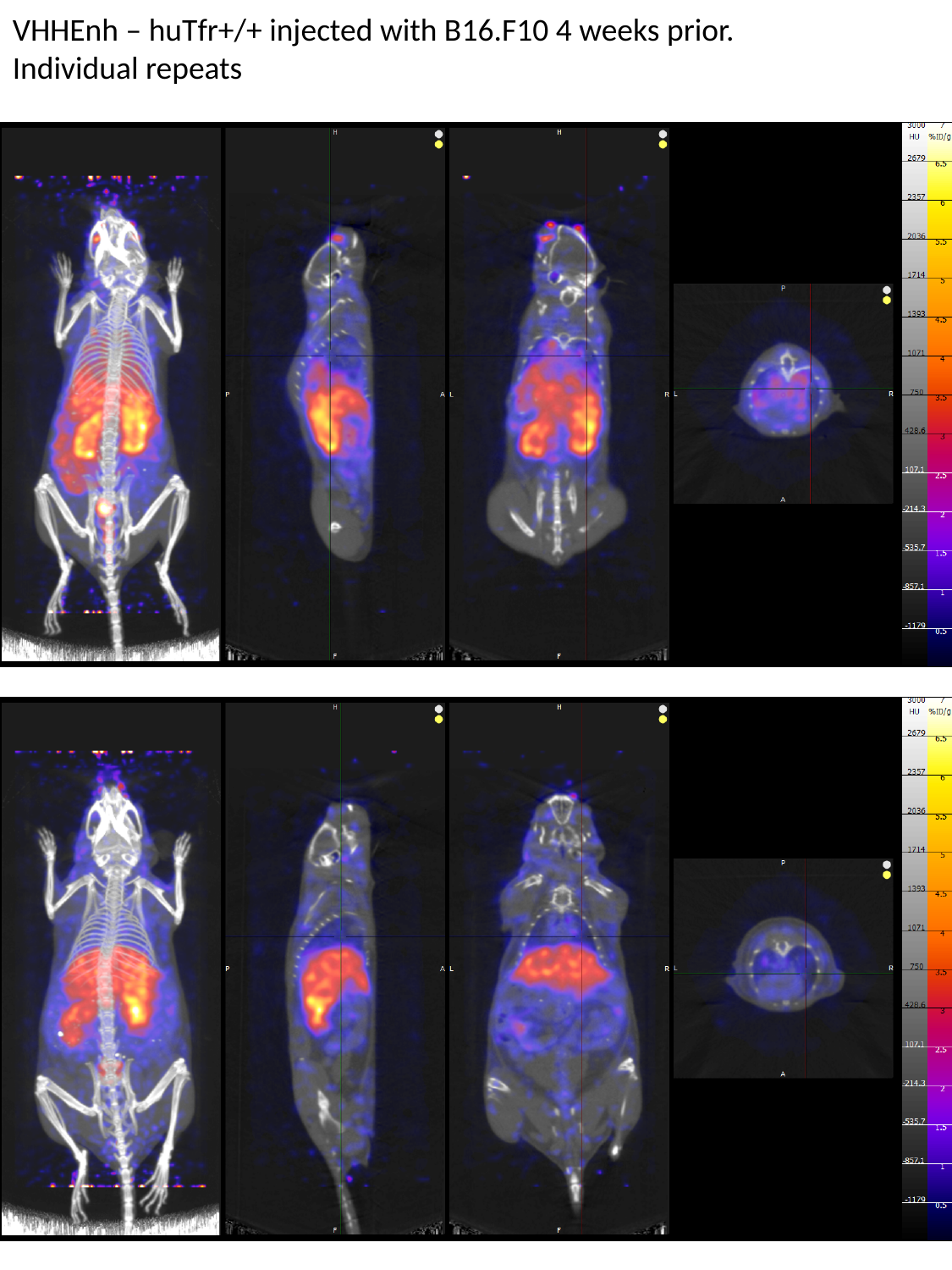

VHHEnh – huTfr+/+ injected with B16.F10 4 weeks prior.
Individual repeats

### Slide 24
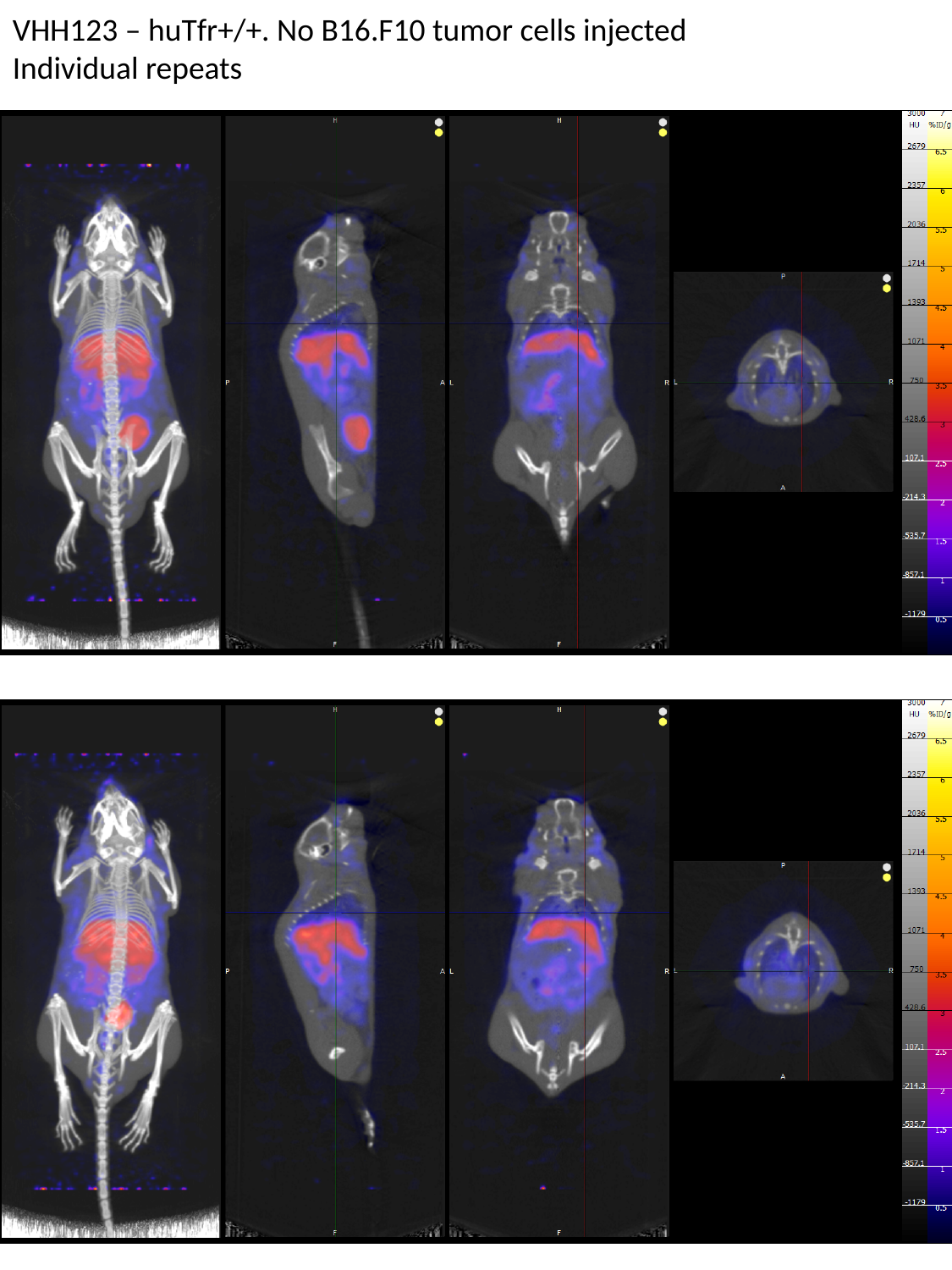

VHH123 – huTfr+/+. No B16.F10 tumor cells injected
Individual repeats

### Slide 25
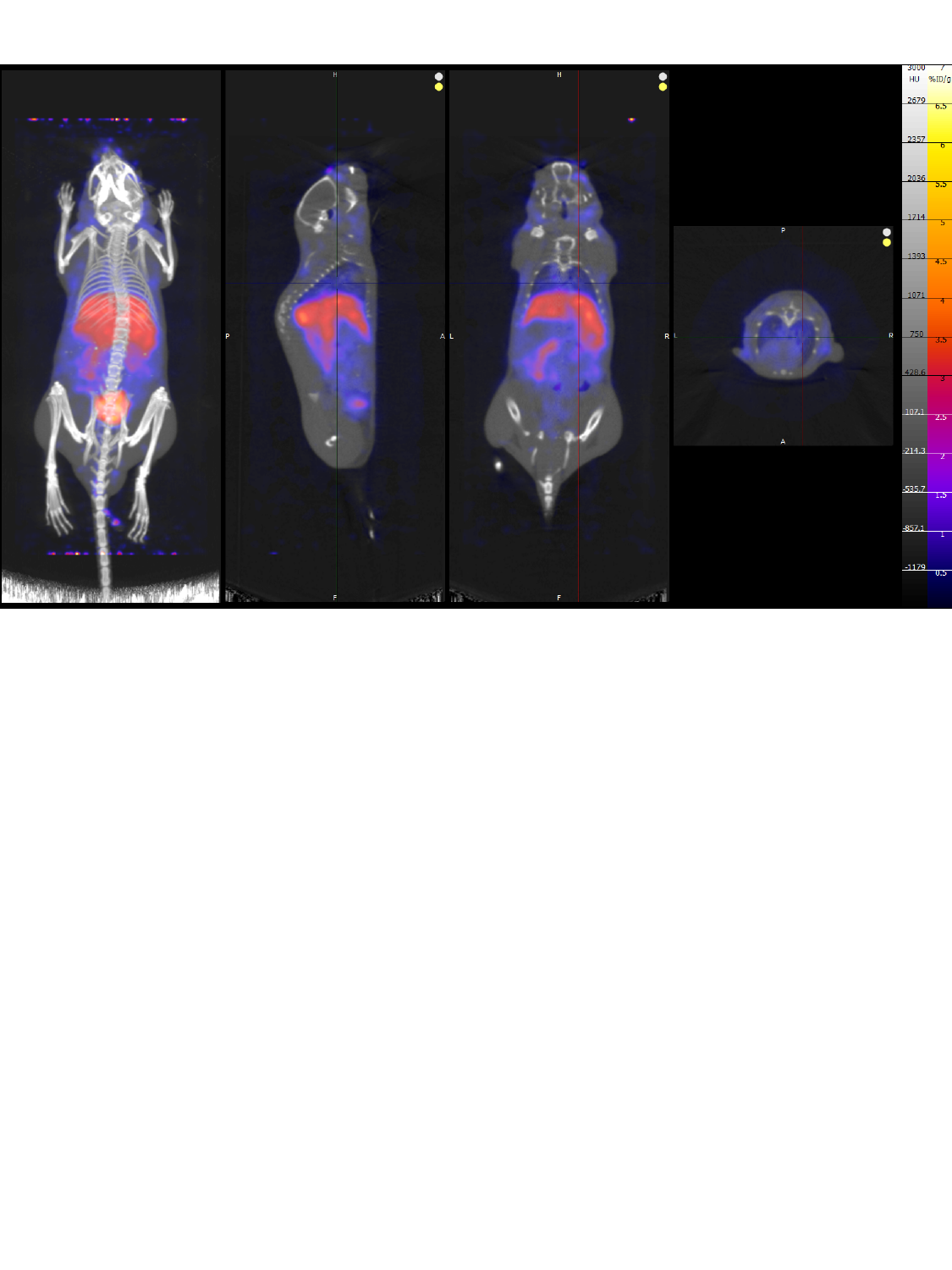

### Slide 26
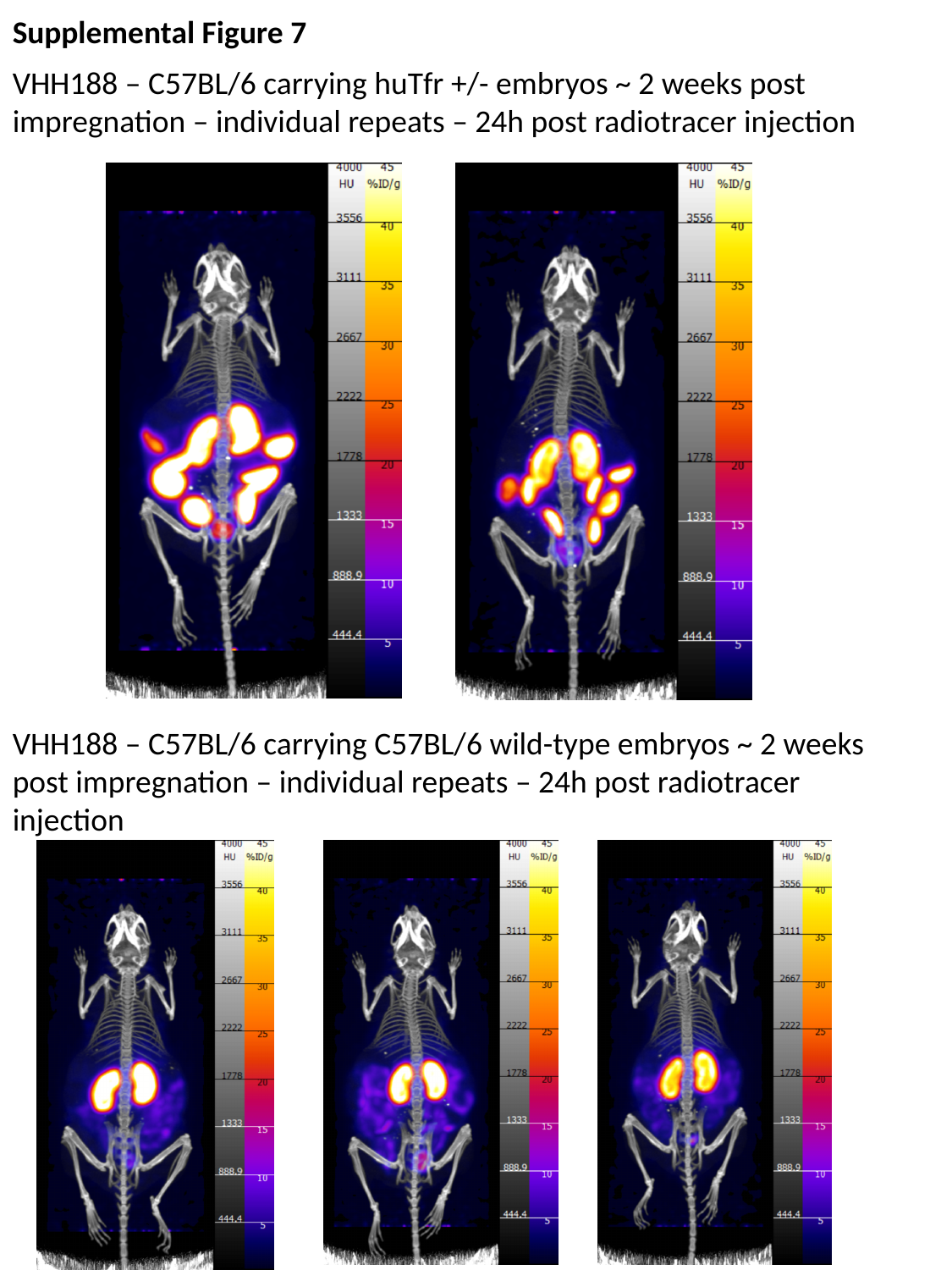

Supplemental Figure 7
VHH188 – C57BL/6 carrying huTfr +/- embryos ~ 2 weeks post impregnation – individual repeats – 24h post radiotracer injection
VHH188 – C57BL/6 carrying C57BL/6 wild-type embryos ~ 2 weeks post impregnation – individual repeats – 24h post radiotracer injection

### Slide 27
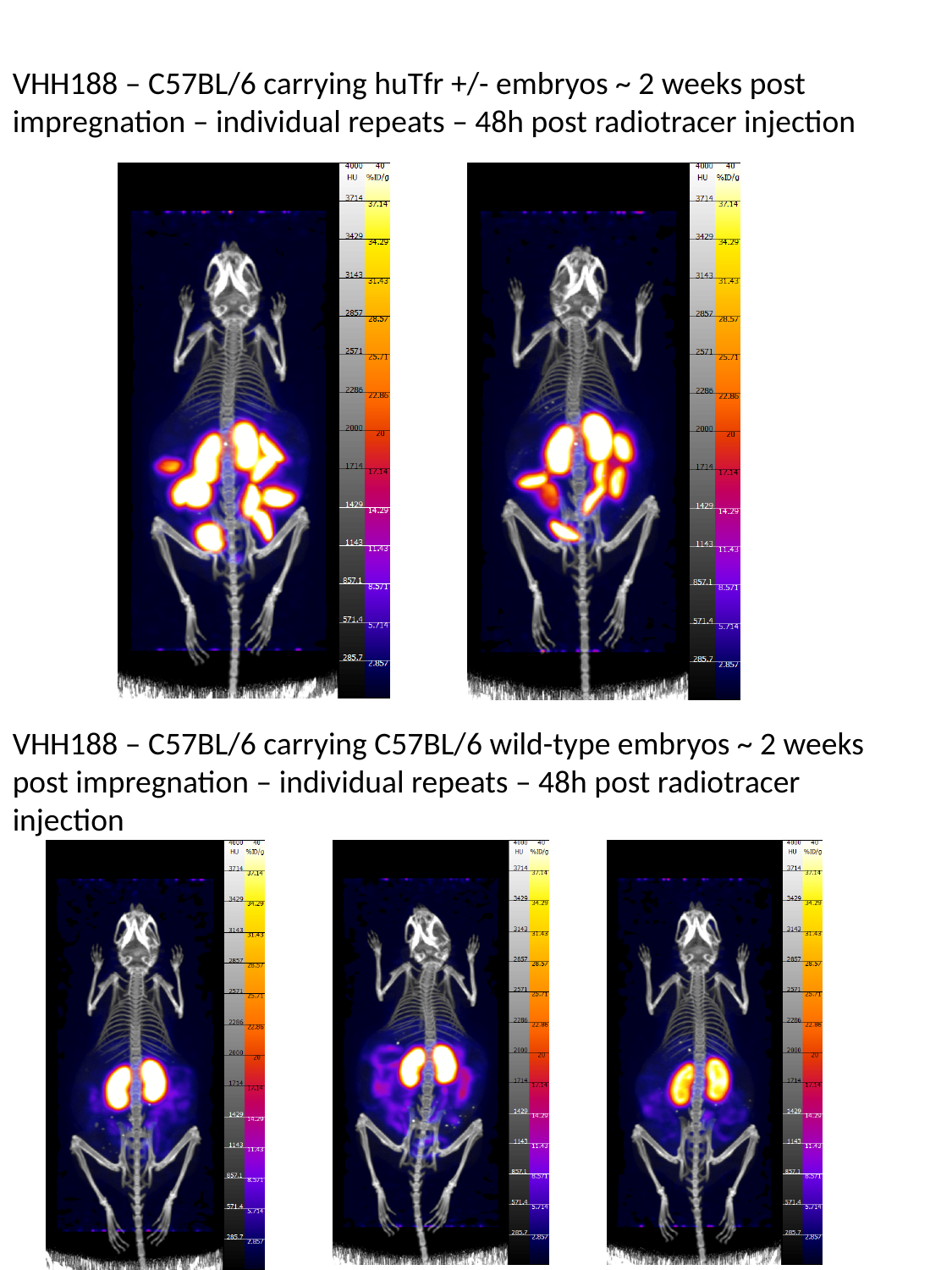

VHH188 – C57BL/6 carrying huTfr +/- embryos ~ 2 weeks post impregnation – individual repeats – 48h post radiotracer injection
VHH188 – C57BL/6 carrying C57BL/6 wild-type embryos ~ 2 weeks post impregnation – individual repeats – 48h post radiotracer injection

### Slide 28
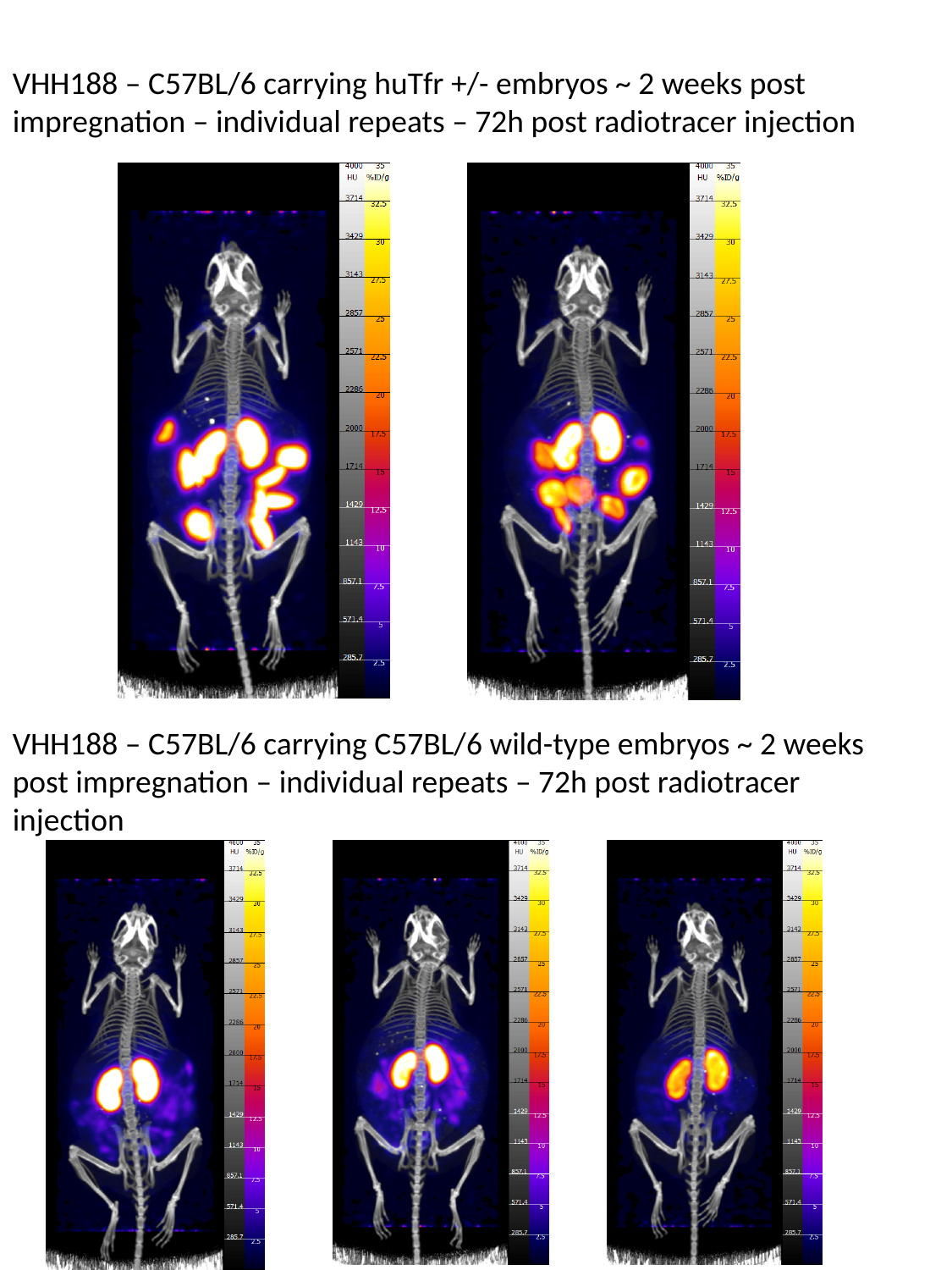

VHH188 – C57BL/6 carrying huTfr +/- embryos ~ 2 weeks post impregnation – individual repeats – 72h post radiotracer injection
VHH188 – C57BL/6 carrying C57BL/6 wild-type embryos ~ 2 weeks post impregnation – individual repeats – 72h post radiotracer injection
